# Sitetack: A Deep Learning Model that Improves PTM Prediction by Using Known PTMs

**DOI:** 10.1101/2024.06.03.596298

**Authors:** Clair S. Gutierrez, Alia A. Kassim, Benjamin D. Gutierrez, Ronald T. Raines

**Affiliations:** Department of Chemistry, Massachusetts Institute of Technology, Cambridge, Massachusetts, United States; Broad Institute of MIT and Harvard, Cambridge, Massachusetts, United States; Koch Institute for Integrated Cancer Research at MIT, Cambridge, Massachusetts, United States

## Abstract

Post-translational modifications (PTMs) increase the diversity of the proteome and are vital to organismal life and therapeutic strategies. Deep learning has been used to predict PTM locations. Still, limitations in datasets and their analyses compromise success. Here we evaluate the use of known PTM sites in prediction via sequence-based deep learning algorithms. Specifically, PTM locations were encoded as a separate amino acid before sequences were encoded via word embedding and passed into a convolutional neural network that predicts the probability of a modification at a given site. Without labeling known PTMs, our model is on par with others. With labeling, however, we improved significantly upon extant models. Moreover, knowing PTM locations can increase the predictability of a different PTM. Our findings highlight the importance of PTMs for the installation of additional PTMs. We anticipate that including known PTM locations will enhance the performance of other proteomic machine learning algorithms.

---

Posttranslational modifications (PTMs) are chemical alterations to proteins following their biosynthesis^1,2^. Typically catalyzed by enzymes, PTMs expand the size of the human proteome by orders of magnitude and are critical to the proper function of cells and organisms^3-7^. Despite advances in mass spectrometry and other experimental methods, the large-scale analysis of PTMs remains challenging and costly^8,9^. Machine learning-based methods, particularly deep learning methods, have been useful for predicting the location of some PTMs from amino acid sequences alone^10^. Many models have been developed in recent years^11-17^ Still, the accuracy of prediction is not highly reliable for the vast majority of PTMs.

We reasoned that including the location of *known* PTMs along with sequence information could improve PTM prediction. Our approach has biochemical precedent. For example, crosstalk is well-known between phosphorylation and O-linked glycosylation at the same and proximal serine or threonine residues^18,19^. Kinase-recognition sites themselves often contain phosphorylated residues^20^. Likewise, a variety of PTMs can affect the ubiquitination of a protein^21^. More broadly, PTM crosstalk pairs have been compiled from 82 human proteins and found to correlate with sequence co-evolution^22^.

Despite the importance of PTMs in directing other PTMs, computational methods that deploy these data for PTM prediction are rare and of limited scope^23^. Moreover, we are not aware of a systematic evaluation of how PTMs influence other PTM sites of the same type. Here, we present “Sitetack” and show that its inclusion of known PTMs improves the prediction of other PTM sites significantly.

## Results

### PTM data improves model performance

Given the success of convolutional neural network (CNN) architectures, we employed a simple CNN-based deep-learning model for testing. Although other architectures were tested, including long-short term memory (LSTM) models, attention-based models, and combinations thereof (Supplementary Fig. 4.2), a simple CNN tended to perform well and have reasonable training times. The overall model architecture used for this study is shown in Fig. 1. For each dataset, we trained two separate models with identical architectures, one that contained other known PTM locations (of the same type of PTM) and one that did not. The former is unique to this work. This training was done using known datasets^16^ for thirteen PTMs: phosphorylation of serine and threonine, phosphorylation of tyrosine, N-glycosylation of asparagine, O-glycosylation of serine and threonine, ubiquitination of lysine, SUMOylation of lysine, acetylation of lysine, methylation of lysine, methylation of arginine, pyroglutamylation of glutamine, palmitoylation of cysteine, hydroxylation of proline, and hydroxylation of lysine (Supplementary Fig. 1.2). For O-glycosylation, we trained additional models for subtype-specific models in which the glycan was either GlcNAc or GalNAc using datasets taken from O-GlcNAcAtlas^24^ and OGP^25^. In addition, models were trained on datasets that include PTMs known to exist only in humans or in any organism.

**Fig. 1.**
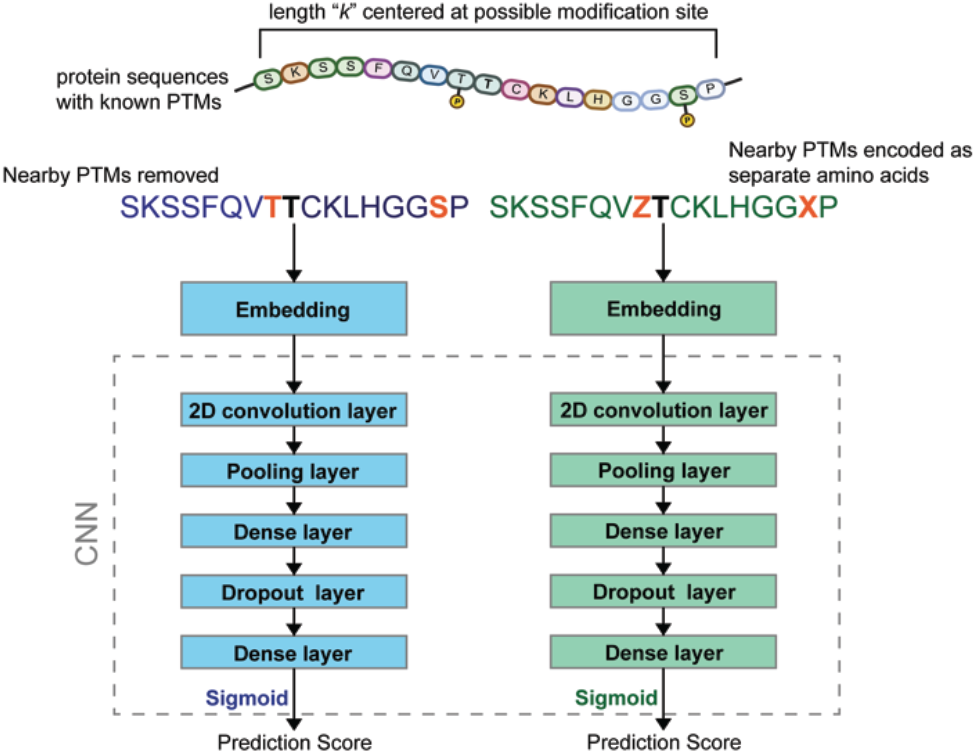
Overall framework of the Sitetack model for PTM prediction.

For a majority of the models we trained, we observed an increase in prediction performance when known PTM locations were encoded as separate amino acids. This increase was most striking for phosphorylation, which went from an AUC of 0.881 to 0.931 for the human-only model and 0.891 to 0.928 for the all-organism model. Select ROC curves are shown in Fig. 2, and detailed performance metrics for all trained models are shown in Supplementary Fig. 5.1, 5.2, and 7.1.

**Fig. 2.**
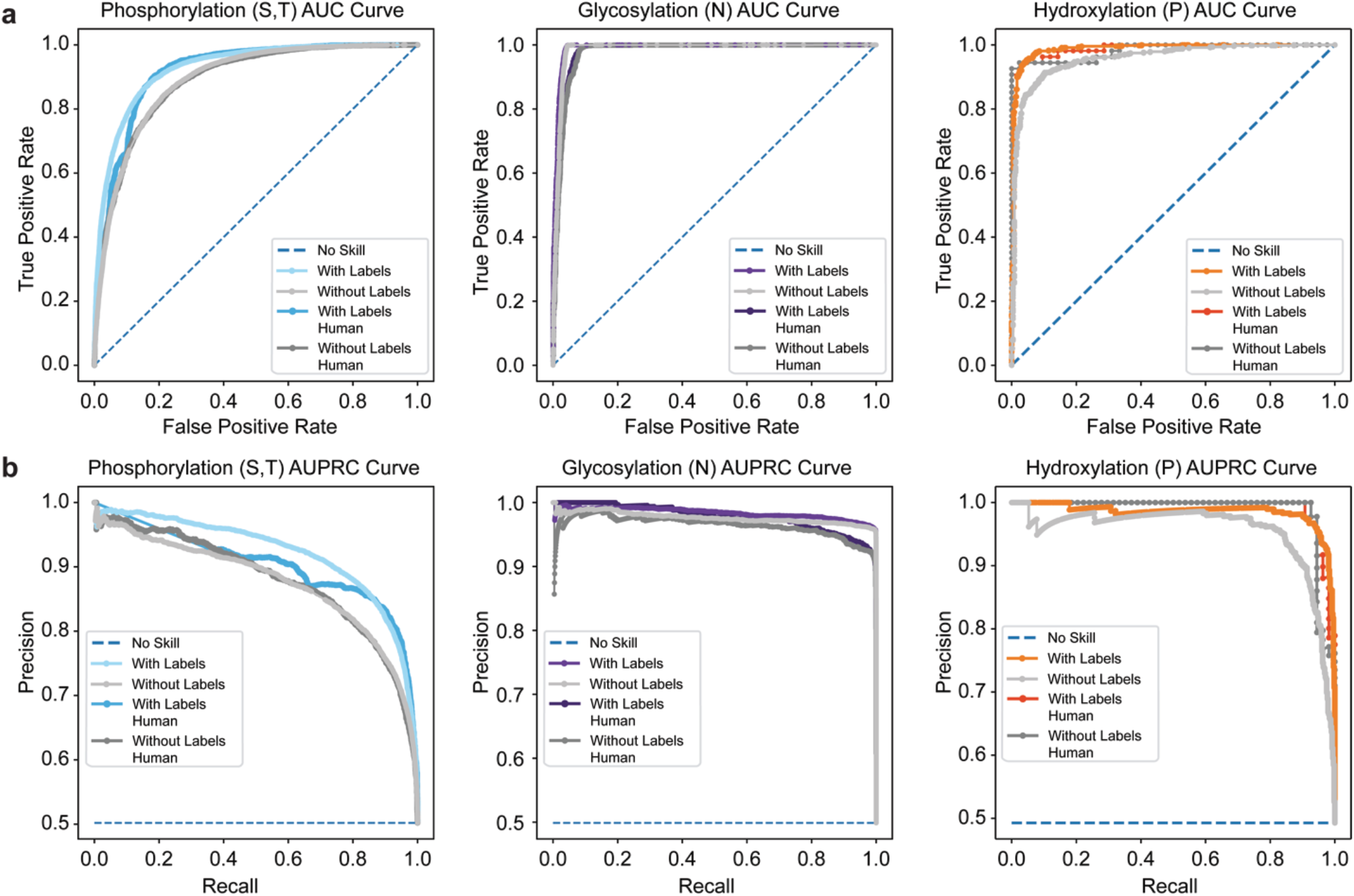
Prediction of three representative PTMs with and without the use of labels: serine or threonine phosphorylation, N-linked glycosylation, and proline hydroxylation. **a**, AUC curves. **b**, AUPRC curves.

Performance enhancements correlated with the size of the dataset of known PTMs. PTMs with larger datasets of known PTMs had larger improvements in prediction performance when known those PTMs were labeled a priori. An exception was N-glycosylation, which had a large dataset and did not have improved performance when known sites were included (Fig. 2). This anomaly was likely due to the simplicity of the N-glycosylation recognition sequence, so we examined N-X-S/T sequon-specific models explicitly. It is a relatively easy task to predict if a site *can* be N-glycosylated, but it is more difficult to distinguish which of the N-X-S/T sequons are actually N-glycosylated. For this task, we trained two sets of models using data from NGlyDE^26^ and NGlycositeAtlas^27^, which were compiled by others^28^. We saw an improvement in performance upon the inclusion of known N-glycosylation sites, indicating that known sites could be helpful in this task (Table 1 and Supplementary Fig. 8). Still, the best models for this task use structural inputs, and these models can perform better than our purely sequence-based models, suggesting that other factors are at play beyond nearby PTMs^28,29^.

**Table 1.** Comparison of PTM predictions for all-organism datasets by Sitetack and other models. Representative models for each PTM were chosen and Sitetack was trained on the same datasets as those models with and without known PTM locations. Values in bold typeface refer to the largest AUC and AUPRC value for each PTM among the three models: Sitetack − PTMs, Sitetack + PTMs, and a comparitor.

| PTM type (residue) | Subtype | AUC/AUPRC |  | Comparator | Comparator Model |
| --- | --- | --- | --- | --- | --- |
|  |  | Sitetack – PTMs | Sitetack + PTMs |  |  |
| Phosphorylation (S,T) |  | 0.869/0.853 | **0.928<sup>a</sup>/0.922<sup>a</sup>** | 0.896/0.329 | MusiteDeep<sup>16</sup> |
| Phosphorylation (Y) |  | 0.892/0.856 | 0.901/**0.869** | **0.958**/0.864 | MusiteDeep<sup>16</sup> |
| N-Glycosylation (N) |  | 0.982/0.969 | 0.989<sup>a</sup>/**0.985<sup>a</sup>** | **0.993**/0.937 | MusiteDeep<sup>16</sup> |
| O-Glycosylation (S,T) |  | 0.947/0.933 | **0.960/0.944** | 0.943/0.539 | MusiteDeep<sup>16</sup> |
| Ubiquitination (K) |  | 0.888/0.801 | **0.923<sup>a</sup>/0.862<sup>a</sup>** | 0.804/0.279 | MusiteDeep<sup>16</sup> |
| SUMOylation (K) |  | 0.957/0.930 | 0.966/**0.945** | **0.990**/0.881 | MusiteDeep<sup>16</sup> |
| Acetylation (K) |  | 0.863/0.761 | 0.904<sup>a</sup>/0.836<sup>a</sup> | **0.978/0.858** | MusiteDeep<sup>16</sup> |
| Methylation (K) |  | 0.965/0.905 | **0.968/0.917** | 0.951/0.850 | MusiteDeep<sup>16</sup> |
| Methylation (R) |  | 0.928/0.870 | **0.945<sup>a</sup>/0.906<sup>a</sup>** | 0.941/0.844 | MusiteDeep<sup>16</sup> |
| Pyroglutamylation (Q) |  | 0.961/0.941 | **0.980/0.978** | 0.979/0.947 | MusiteDeep<sup>16</sup> |
| Palmitoylation (C) |  | 0.949/0.924 | **0.972/0.949** | 0.961/0.922 | MusiteDeep<sup>16</sup> |
| Hydroxylation (P) |  | 0.901/0.856 | **0.980<sup>a</sup>/0.968<sup>a</sup>** | 0.732/0.627 | MusiteDeep<sup>16</sup> |
| Hydroxylation (K) |  | 0.918/0.968 | 0.947<sup>a</sup>/0.876<sup>a</sup> | **0.982/0.930** | MusiteDeep<sup>16</sup> |
| N-Glycosylation (N) | Sequon-specific | 0.656/0.482 | 0.687/0.521 | **0.907/0.857<sup>28</sup>** | DeepNGlyPred<sup>27</sup> |
| O-Glycosylation (S,T) |  | 0.829/0.832 | 0.953/**0.963** | **0.983**/0.915<sup>b</sup> | OGP SVM<sup>25</sup> |
| O-Glycosylation (S,T) | GalNAc | 0.859/0.872 | **0.963/0.968** | — | OGP SVM<sup>25</sup> |
| O-Glycosylation (S,T) | GlcNAc | 0.766/0.767 | **0.879/0.890** | 0.828/— | O-GlcNAcPRED-DL<sup>7,24</sup> |

<sup>a</sup>A significant difference ( $P < 0.05$ ) exists between the Sitetack models with and without known PTM locations. <sup>b</sup>This model was trained on an unspecified subset of OGP.

### Comparison with existing methods

We compared the AUC and AUPRC of Sitetack with and without known PTMs compared to representative models trained on similar datasets. Because we used the same datasets as MusiteDeep and few other models study a similar breadth of PTMs, our main comparison was with MusiteDeep. In general, for our models trained *without* known PTM locations, MusiteDeep outperformed Sitetack, which is not surprising given its use of a more complex architecture. Yet, when known PTM sites were included, Sitetack outperformed MusiteDeep (AUC or AUPRC) for most PTMs (Table 1). For O-glycosylation, we tried several other models with a larger dataset or subtype-specific models. For those models, Sitetack with labels generally outperformed corresponding models. For GalNAc-specific models, there were no models trained on a similar dataset. Nonetheless, including known PTM locations increased prediction reliability, and partitioning of the GalNAc sites was beneficial compared to an analysis of the overall OGP dataset.

### Proximity of PTMs to other PTMs

To understand why nearby PTM sites improve model performance, we examined the frequency of nearby PTM sites in a given dataset at a target residue. The results followed three patterns, as shown in Fig. 3. (All PTMs are shown in Supplementary Fig. 3.) In the first pattern, as is evident with N-glycosylation, we found a relatively even distribution of PTMs at different distances from the site of interest. In this pattern, there is little benefit to knowing the location of nearby PTMs. Accordingly, nearby PTMs are more likely due to random chance than the substrate recognition necessary for N-glycosylation. The second pattern is illustrated best by the hydroxylation of proline, which has a strongly repetitive pattern. The emergence of this pattern is not surprising because collagen is the most common protein with hydroxyproline and follows a triplet repeating pattern of Xaa-Yaa-Gly in which Yaa is hydroxyproline 38% of the time^30^. In this pattern, the inclusion of nearby PTMs improves prediction only slightly, which is likely due to the simplicity of the pattern that a model could learn. The third pattern is illustrated by the phosphorylation of serine and threonine, whereas as distance increases from a given site, there are fewer nearby phosphorylation events. In such cases, we saw that including known nearby PTMs in the dataset improved predictions significantly. We hypothesize that enzymes that catalyze these modifications are influenced by amino acid residues that have already undergone a PTM.

**Fig. 3.**
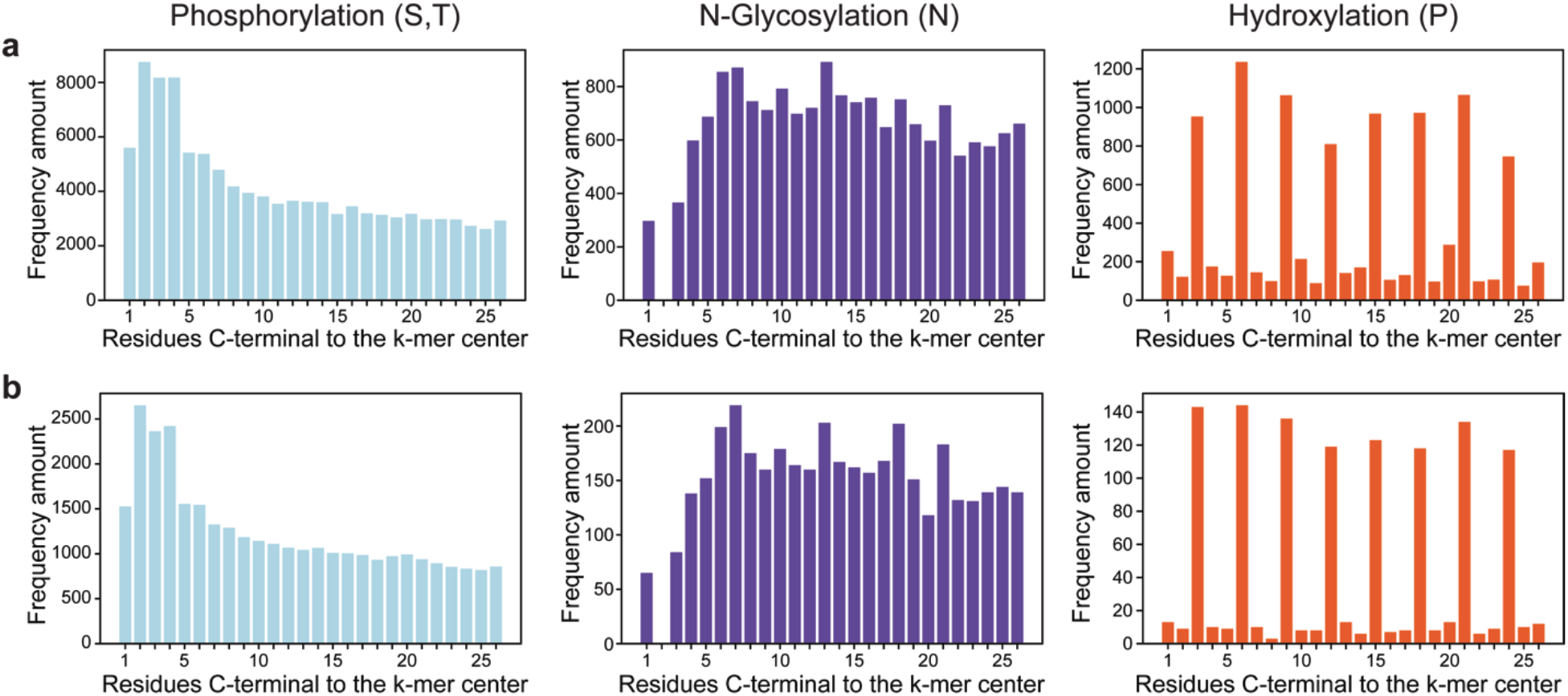
Frequency of nearby PTM sites for three representative PTMs: serine or threonine phosphorylation, asparagine N-glycosylation, and proline hydroxylation. **a**, All-organism datasets. **b**, Human-only datasets.

### Extension to kinase-specific models

The human kinome contains more than 500 kinases^31^. Accordingly, the identity of the kinase that catalyzes a particular phosphorylation event is useful information. Towards this goal, we developed kinase-specific models^11,32^ to determine whether or not nearby phosphorylation sites were important for prediction at the enzymatic level.

The datasets are necessarily much smaller for a particular kinase than for phosphorylation in general. We envisioned that a reasonable approach would be to predict sites that might be phosphorylated in a protein of interest and then to see which kinase might be responsible for the phosphorylation of each highly scoring site. We were not interested in which kinase is responsible in natural systems per se, as that question also requires a consideration of which phosphatases might be present and likely have a smaller training set. Instead, we asked a more prudent question: Which kinases could target a particular site? To answer this question, we selected a dataset from a publication by Ishihama and coworkers^33^ due to its breadth in the number of kinases assayed, number of sites reported, and use of an *in vitro* assay. From this dataset, we looked at 68 kinases that target serine or threonine residues and that had over 500 reported sites (Supplementary Fig. 9.1). For each of these kinases, we trained models with and without known phosphorylation sites (Supplementary Fig. 9.2 and 9.3). Overall, we saw that the inclusion of known phosphorylation sites improved kinase-specific prediction, with some kinases more affected than others (Fig. 4). To understand why some kinase models were influenced more by the inclusion of known sites, we looked at three examples: NEK4, which had the largest difference in AUC between the two models; IKKε, which was near the median difference in AUC between the two models; and PKA Cα, which had the smallest difference between the AUC of the two models. For these models, the number of nearby phosphorylation sites greatly influenced the performance of the labeled model (Fig. 4b).

**Fig. 4.**
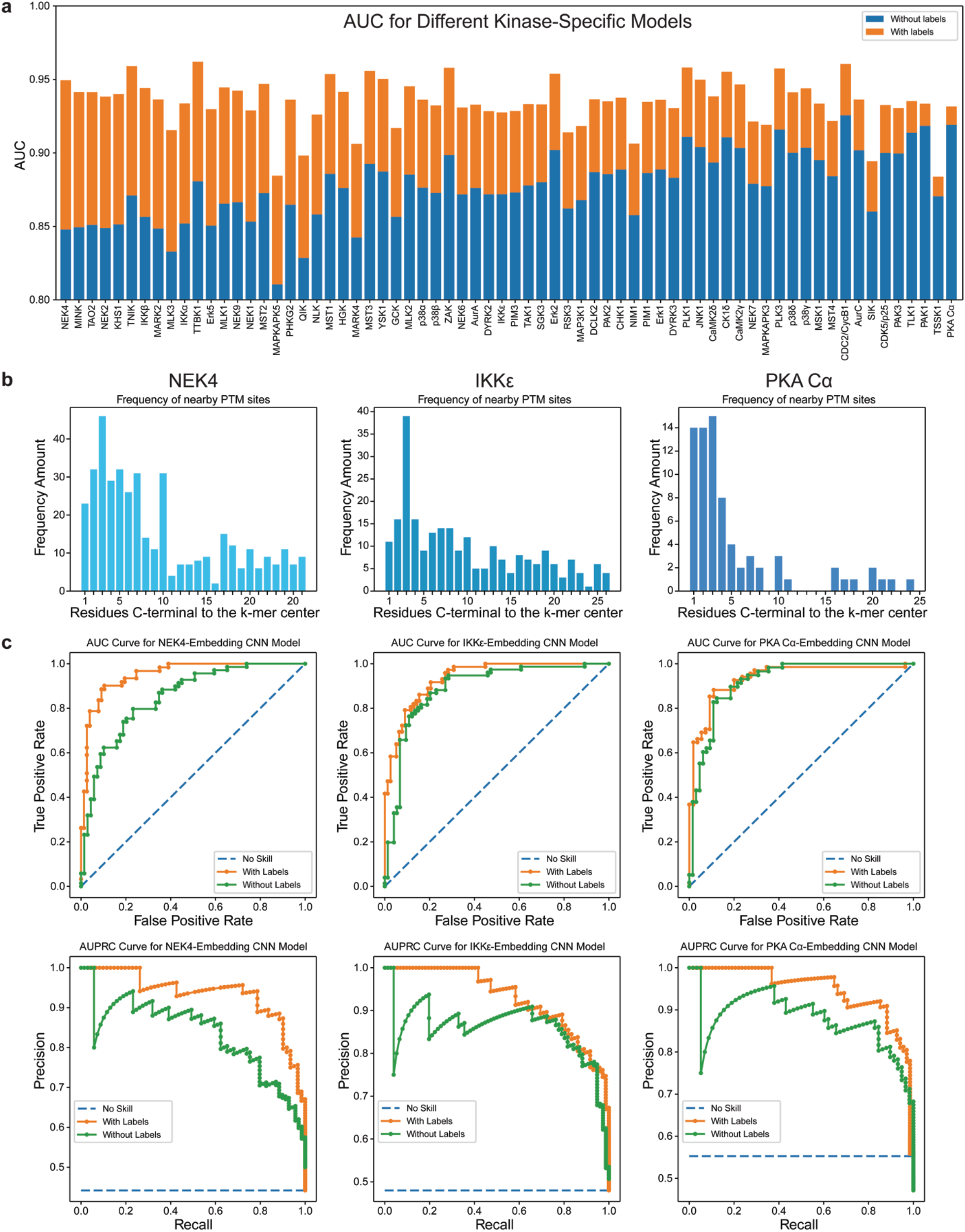
Phosphorylation (S,T) prediction using kinase-specific models. **a**, Difference in AUC with and without known site locations. **b**, Frequencies of nearby sites in the dataset. **c**, AUC and AUPRC curves.

### Cross PTM models

Having shown that encoding known PTM locations of a given type as a separate amino acid can improve prediction accuracy for that PTM, we wondered if including information about a *different* PTM can improve prediction accuracy. To explore this crosstalk, we chose O-glycosylation with GlcNAc and phosphorylation at serine and threonine, which generally occur in the same subcellular compartments and at the same amino acid residues (serine and threonine)^18,19,34^. Moreover, both had large datasets, which we suspected was important to detect an impact on prediction accuracy^24^. We chose to look at O-GlcNAc prediction using phosphorylation sites and not the converse, given that the size of the phosphorylation dataset was much larger. For this task, we trained two separate models with known PTM locations, one in which known phosphorylation sites were encoded as separate amino acids (but not known O-GlcNAc sites) and another in which known O-GlcNAc sites were encoded as a separate amino acid but not phosphorylation sites. The result was that both phosphorylation sites and O-GlcNAc sites improved prediction, though O-GlcNAc sites did so to a much higher level (Fig. 5a,c) in the all-organism datasets. There was an improvement, but not a significant one, in the human-only datasets (Supplementary Fig. 10). We found that phosphorylation sites were present near the prediction residue but not at the level that O-GlcNAc sites occurred (Fig. 5b), which offers an explanation for O-GlcNAc sites showing a larger improvement. Thus, known PTM locations are able to improve the prediction of a different PTM.

**Fig. 5.**
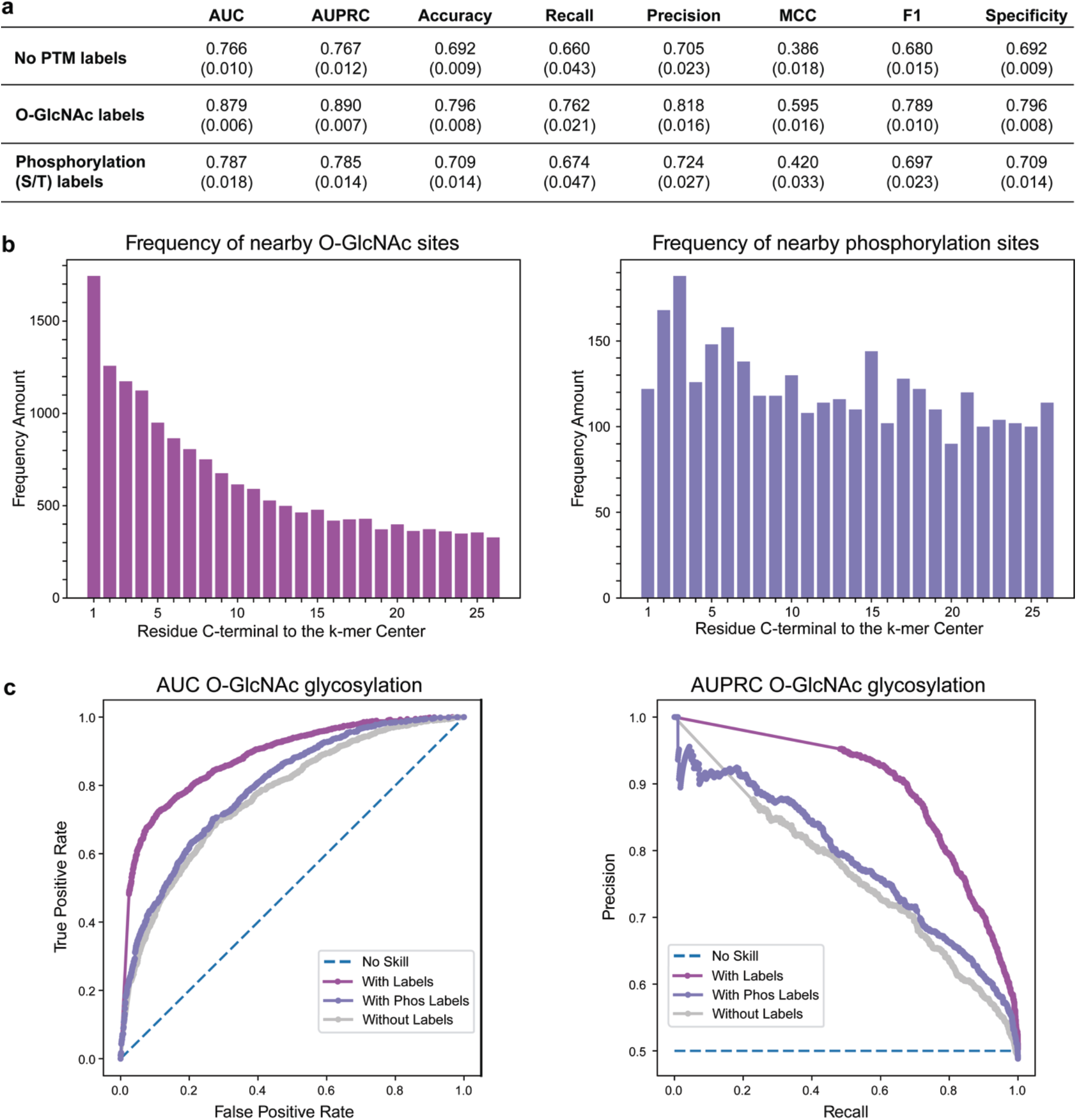
O-GlcNAc prediction using known O-GlcNAc or phosphorylation sites. **a**, Performance assessments with and without labels. **b**, Frequencies of nearby sites in the dataset. **c**, AUC and AUPRC curves.

### Accessibility of models

To make this tool accessible to researchers, we incorporated our best models into a free web tool that is accessible at https://sitetack.net. Users can input multiple sequences of interest and select a model to use for PTM prediction. For sequences with known PTM locations, there are instructions for each model on how to encode that information (*e*.*g*., a phosphoserine is encoded as @). Results are displayed on the page for a single input, and multiple inputs can be downloaded as a CSV file. The website includes models for every PTM from this work, trained with and without known PTM locations.

We also developed simple scripts for researchers to generate their own models for any PTM dataset. We envision these scripts as being useful for researchers who have compiled sites for a given enzyme or rare PTM that was not evaluated here, allowing them to train prediction models to aid in their research. This functionality is available at https://github.com/clair-gutierrez/sitetack, along with step-by-step instructions.

## Discussion

This work demonstrates that encoding known PTM locations as separate amino acids is a relatively simple way to improve the performance of deep learning PTM prediction models for many PTMs. From a biochemical perspective, this finding is not surprising, given that PTMs alter the chemical, structural, and functional landscape of proteins^1,2^. We found that the extra data were especially useful in prediction tasks with large datasets like phosphorylation, which is intelligible from a technical standpoint as a certain level of coverage is necessary for PTMs to be near each other.

We found that kinase-specific models generally benefited from the inclusion of known phosphorylation locations more so than did most other models. This benefit might be due to the sensitivity of protein kinases to amino acid sequence^20^ and that the modification installs a compact dianionic functional group. We anticipate that future work on the development of kinase- or kinase family-specific models would benefit greatly by including known phosphorylation sites. Additionally, context-specific information could be learned with models trained on phosphorylation levels in different contexts (*e*.*g*., cell types, hypoxia, or stress states).

We also showed that phosphorylation sites can improve the prediction of O-glycosylation via O-GlcNAc. These models did not perform especially well for O-GlcNAc prediction but were still significantly better than models that omitted phosphorylation data. In future work, we foresee probing which PTMs affect the prediction of other PTMs, like ubiquitination, of biological relevance. The ensuing crosstalk underscores the importance of PTM selection in prediction tasks, as the choice of which PTMs to include can affect the resulting prediction.

Including PTMs in training datasets encoded as unique amino acids has broad utility beyond PTM prediction. PTMs are known to affect the structure, function, and activity of proteins and could improve other models ranging from structure prediction to de novo protein design. Additionally, many current PTM models lack temporal information because many modifications, such as phosphorylation, O-glycosylation, and N-glycosylation, are physiologically reversible. More tools that allow researchers to see what modifications are likely in different contexts would allow for more information on proteins of interest and begin to reveal situational information. Encoding PTM locations can improve prediction and, hence, the reliability of deep learning PTM prediction models and is poised for application to different prediction tasks.

## Methods

### Benchmark datasets

In this study, we deployed several previously curated PTM datasets that had been used to train different machine-learning models. Many of our models used the various PTM datasets compiled by Xu and workers^16^ that were obtained from www.musite.net.^11,14,16^ For O-glycosylation specifically, we used the OGP dataset from Li and coworkers^25^ because the dataset from Xu and coworkers^16^ was relatively small. Additionally, to obtain a large dataset for O-GlcNAc O-glycosylation, we used a dataset processed from O-GlcNAc site atlas^24^ that was used by Jia, Ma, and coworkers^7^ in their prediction model. For N-glycosylation, the specificity for the N-X-S/T sequon led us to use two additional datasets from KC and coworkers^28^, which curated positive and negative examples specifically for the N-X-S/T sequon. For kinase-specific models, the datasets from Ishihama and coworkers^33^ were processed to remove cross sites and sites not in reference proteome UP000005640. Of these, only datasets containing over 500 sites were used in order to have better training.

To evaluate model ability, we split each dataset into training, validation, and test sets in the ratio of 80%:10%:10% selected at random. Though the composition of these sets varied with the specific models and different validation runs, the test set was always completely distinct from the training set for a given evaluation. To further validate models, 10-fold validation was performed on the best models by resplitting the datasets, retraining, and retesting separately.

Model statistics were reported as an average of the ten trials, with an error reported as the standard deviation of the trials.

### Data representation

For a given protein sequence, subsequences or k-mers of length k centered at possible PTM sites were generated with various window sizes. Optimization was performed on the best size of “k” by retraining the CNN model for serine or threonine phosphorylation with different sizes of “k” (Supplementary Fig. 4.1). For all subsequent models, optimization of k was not performed due to the number of models that needed to be trained, so a size of k = 53 was chosen. For each type of PTM in models that contained PTM information, known PTMs of the same type were encoded as a separate amino acid. For human-only models this was generally “X” (and “Z”, if there was another type of amino acid residue that could undergo modification). In the all-organism datasets, “X” and “Z” were often present as unknown amino acids or glutamine/glutamic acid respectively, so the symbols “@” and “&” were used. For each model, the alphabet of amino acids used can be found in Supplementary data 1. For all datasets the 20 canonical amino acids were encoded as standard along with “-” to denote no amino acids (for sites near the ends of a protein) and “U” for selenocysteine. For the models used, the amino acids were then embedded into a vector where each amino acid corresponded to a different number before feeding it into the model. For example, the sequence “ACD” would become <1, 2, 3>.

Given that there are many more negative examples than positive examples, for each model, we randomly selected the same number of negative sites as there were positive sites, except in the case of N-X-S/T sequon glycosylation, which already had a relatively balanced dataset. During 10-fold validation, we randomly selected negatives again, such that the negative datasets used in each validation run differed significantly. Additionally, to handle homology between the kmers themselves for our main models, we ensured that no sequence in any test set was identical to the cognate train or validation sets. For each model, we evaluated different sequence identity cutoffs down to 40% in the kmers (which is stricter than on a whole protein level) using CD-HIT,^35,36^ which can be seen in Supplementary Fig. 11. Most performances suffer only minor reductions with lower sequence similarities.

The human-only datasets were curated to contain only human proteins by taking only those in UniProt reference proteome UP000005640 for *Homo sapiens*^37^. For all-organism datasets, because of the vast size of an interrogation of all proteins from all organisms, negatives were taken from unmodified sites in proteins that had the PTM. In this case “reference proteomes” were generated by obtaining the sequences for the modified proteins from UniProt.

### Model architecture

The general deep learning architecture for this work is shown in Fig. 1. Several other architectures were tried, including an LSTM and attention-based model (Supplementary Fig. 4.2). In short, the general model used in this study took the vectorized sequences of length 53, embedded them as a word embedding, then passed them through a 2D convolution layer, a max pooling layer, and two dense layers with a dropout of 0.1 in between. ReLu activation was used for all layers except for the output that used Sigmoid.

### Training

Model parameters were tuned using the human protein phosphorylation dataset (Supplementary Fig. 4). All other models were trained by using the Adam Optimizer^38^ (learning rate = 0.001) with Binary Cross Entropy as a loss function. Early stopping by monitoring the validation set loss with a patience of 15 was used to prevent overfitting. If early fitting was not activated, then the models could train up to 400 epochs using a batch size of 100. Additionally, kernel regularization was used at the convolutional layer via L2 regularization with a weight of (1E−6).

### Performance assessment

To assess the prediction performance of Sitetack, we used several common metrics, including area under the ROC curve (AUC), area under the precision recall curve (AUPRC), accuracy (Acc), sensitivity or recall (Sn), specificity (Sp), precision (Pre), Matthew’s correlation coefficient (MCC), and F1 score (F1). Whereas AUC and AUPRC were calculated based on taking the area under the curve generated by plotting the false positive rate against the true positive rate or the precision against the recall respectively of a dataset, the remaining metrics were defined by the equations:

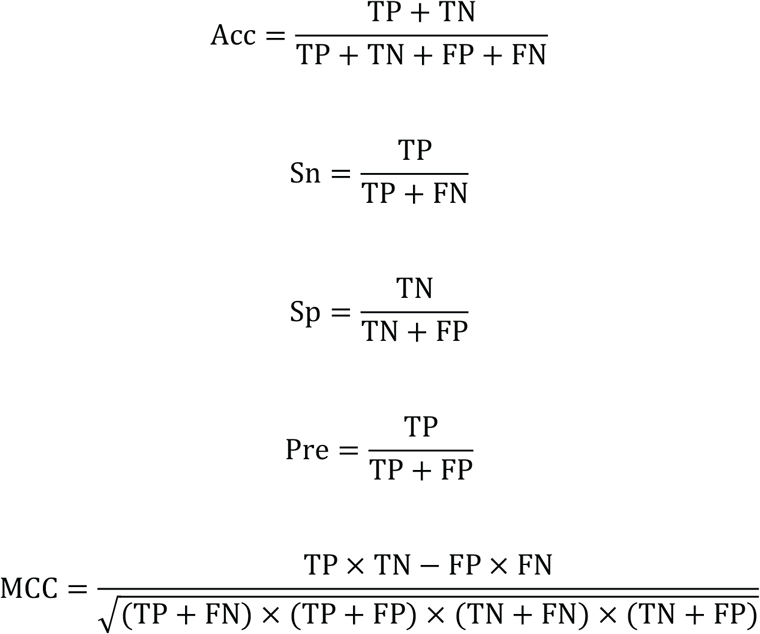

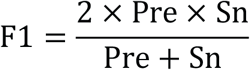

where TP, TN, FP, and FN refer to true positives, true negatives, false positives, and false negatives, respectively.

## Acknowledgements

This work was supported by the National Institutes of Health (R01 CA073808 and R35 GM148220). C.S.G. was supported by an NDSEG Fellowship sponsored by the Air Force Research Laboratory. A.A.K was supported by the MIT Undergraduate Research Opportunities Program. We thank Professors Forest M. White and Matthew D. Shoulders, and Oscar J. Molina for helpful discussions. This work would not have been possible without the MIT Engaging computational cluster and the staff that maintains it.

## Author contributions

C.S.G. conceptualized the project. A.A.K designed and optimized the general deep learning model. C.S.G and A.A.K conducted the algorithm implementation, training, and analysis. C.S.G and B.D.G. created the software and resources for general use. B.D.G. designed and implemented the webserver. C.S.G., A.A.K, and R.T.R. wrote the paper. R.T.R supervised the project. All authors reviewed and approved the paper.

## Competing interests

The authors declare no competing interests.

## Additional information

Sitetack is available as a web tool at: https://sitetack.net for protein PTM prediction using the CNN models reported herein. The source code, representative datasets, instructions for local use, and select models can be found at: https://github.com/clair-gutierrez/sitetack.

## Notes

https://sitetack.net

### Competing Interest Statement

The authors have declared no competing interest.

