## Supplementary for "Sitetack: A Deep Learning Model that Improves PTM Prediction by Using Known PTMs"

---

**Content** **Page**

**1. General Information**
- 1.1. Abbreviations ..... S3
- 1.2. Posttranslational Modifications ..... S3
- 1.3. Datasets ..... S4
- 1.4. Python Libraries and Code ..... S4
- 1.5. Website ..... S5
- 1.6. Statistics ..... S5

**2. Data Preparation**
- 2.1. MusiteDeep Datasets ..... S5
- 2.2. OGP Datasets ..... S5
- 2.3. O-GlcNAc Site Atlas Datasets ..... S5
- 2.4. LMNglyPred Datasets ..... S5
- 2.5. Sugiyama Kinase Datasets ..... S6

**3. Frequency of Nearby PTM Sites**
- 3.1. Generation of PTM Frequency Graphs ..... S7
- 3.2. Frequency of Nearby PTM Sites in Human-Only Datasets from MusiteDeep ..... S7
- 3.3. Frequency of Nearby PTM Sites in All-Organism Datasets from MusiteDeep ..... S11
- 3.4. Frequency of Other O-Glycosylation Sites from OGP and O-GlcNAc Site Atlas ..... S15
- 3.5. Frequency of Other N-Glycosylation Sites from LMNglyPred Datasets ..... S18
- 3.6. Frequency of Other S/T Phosphorylation Sites from Sugiyama Kinase Datasets ..... S19

**4. Tuning of Model Parameters**
- 4.1. Selection of Optimal “k” for k-mers ..... S28
- 4.2. Testing of Other Model Architectures ..... S28
- 4.3. Hyperparameter Tuning ..... S28

**5. Additional Model Results**
- 5.1. Model Results in Human-Only Datasets from MusiteDeep ..... S29
- 5.2. Model Results in All-Organism Datasets from MusiteDeep ..... S32

**6. Additional ROC curves**
6.1. AUC for Models Trained on MusiteDeep Datasets ..... S35
6.2. AUPRC for Models Trained on MusiteDeep Datasets ..... S40
**7. Additional O-Glycosylation Models**
7.1. Model Results ..... S45
7.2. AUC ROC Curves ..... S47
7.3. AUPRC ROC Curves ..... S49
**8. N-Glycosylation Sequon-Specific Models**
8.1. Model Results ..... S51
8.2. AUC ROC Curves ..... S51
8.3. AUPRC ROC Curves ..... S52
**9. Additional Kinase-Specific Model Results**
9.1. List of Kinases Used ..... S52
9.2. Kinase-Specific Model Results ..... S53
**10. Additional Cross-Model Figures**
10.1. Human O-GlcNAc Prediction with Phosphorylation (S/T) Sites ..... S65
**11. Effect of Homology on Model Performance**
11.1. Evaluation of Sequence Identity ..... S67
11.2. Using CD-Hit to Lower Test-Set Identity ..... S68
11.3. Using CD-Hit to Lower Test-Set Identity in OGP Datasets ..... S74
11.4. Using CD-Hit to Lower Test-Set Identity in O-GlcNAc Datasets ..... S74
11.5. Using CD-Hit to Lower Test-Set Identity in N-Glycosylation Sequon-Specific Datasets ..... S74
**12. References** ..... S75

### 1. General Information

#### 1.1. Abbreviations

| | | | |
|---|---|---|---|
| Acc | accuracy | MCC | Matthew's correlation coefficient |
| AUC | area under (ROC) curve | MLP | multilayer perceptron |
| AUPRC | area under precision recall curve | Pre | precision |
| CNN | convolutional neural network | ReLu | rectified linear unit |
| DNN | deep neural network | RF | random forest |
| F1 | F1 score | PTM | posttranslational modification |
| FN | false negatives | ROC | receiver operating characteristic |
| FP | false positives | Sn | sensitivity or recall |
| GalNAc | *N*-acetylgalactosamine | Sp | specificity |
| GlcNAc | *N*-acetylglucosamine | SVM | support vector machine |
| HexNAc | *N*-acetylhexosamine | TN | true negatives |
| LSTM | long short-term memory | TP | true positives |

#### 1.2. Posttranslational Modifications

The chemical structures of the thirteen posttranslational modifications considered in this work are indicated in red below. Note that only monoglycosylation is shown for N- and O-glycosylation.

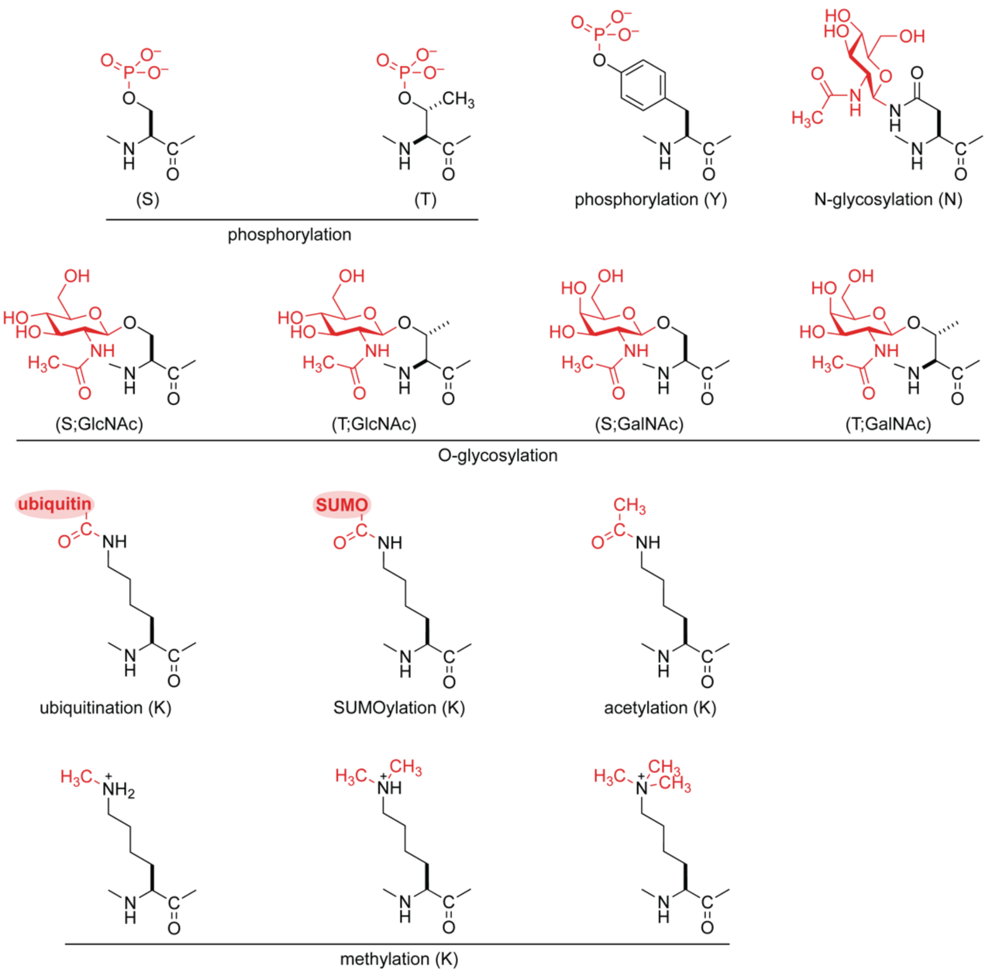

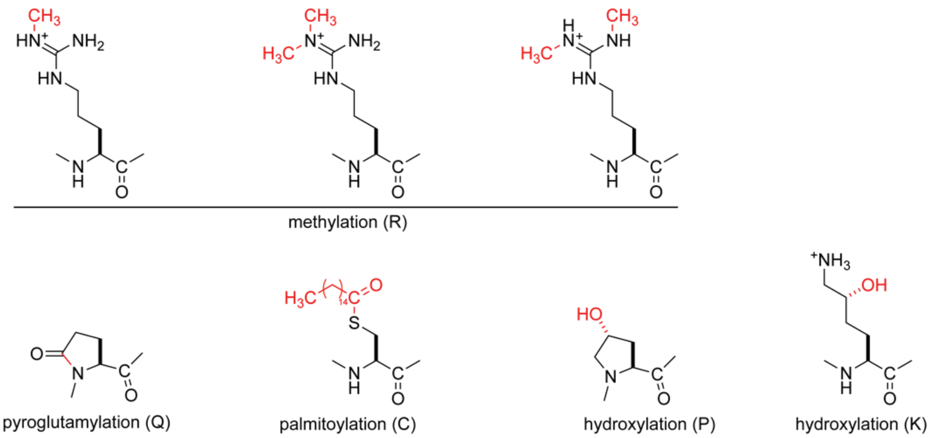


##### 1.3. Datasets

For information specific to the datasets used herein, see Section 2. Unless specified otherwise, the datasets were processed as follows. For each dataset, a list of Uniprot IDs of the modified proteins with a list of sites for each protein was generated. Additionally, a FASTA file of reference sequences was obtained. For human protein models, this file was the UP000005640 reference proteome. For all-organism models, this file was generated by using the ID-mapping function on Uniprot (https://www.uniprot.org/id-mapping; accessed 7 February 2024) and inputting a list of all of the Uniprot IDs of the modified proteins. Generally, isoforms and archaic proteins were removed automatically.

Using these files, a list of positive k-mers was generated from the reference sequences and the list of protein IDs and sites. Negative k-mers were generated by taking a list of all the possible PTM sites (based on what amino acid is modified) and removing sites contained in the list of known PTM sites. From these sites, the same number of negative k-mers as positive k-mers were chosen randomly. In the case of PTMs that could occur at two residues (S and T), then the ratio of sites between the two residues in the positive k-mers was maintained in the negative k-mers. Adding the locations of known PTMs to the k-mers was done by modifying a dictionary of Uniprot IDs and the cognate sequences from the sequence file to change PTM locations to a separate amino acid. Then, positive and negative k-mers with known PTM locations could be generated from this modified dictionary. For these the center amino acid, for which prediction occurs, was converted back to the unmodified version in the positive k-mers (otherwise, we would already be giving the model the correct answer).

For 10-fold validation, a new set of negative k-mers was selected for each repeat and combined with the positive k-mers (with labels as to whether they were negative or positive). These were then shuffled and split into training (80%), validation (10%), and test (10%) datasets. To ensure the test datasets were distinct, k-mers that were identical to any k-mer in the cognate train and validation sets were removed before evaluation.

##### 1.4. Python Libraries and Code

All module versions for python, libraries versions, and other programming information can be found in the Github repository for this work at: https://github.com/clair-gutierrez/sitetack. In short, Jupyter Notebook was used for many of the model training and testing scripts. The TensorFlow library, often with the Keras API, was used for model creation and training. Numpy and pandas libraries were used for dataset manipulation. The Sci-kit learn library was used to analyze model results. The matplotlib library was used to generate figures in Python, which were then further refined with Adobe Illustrator. Microsoft Excel was used to visualize data. Models were trained either locally or on the MIT Engaging cluster.

##### 1.5. Website

The Sitetack webserver consists of a three−layer architecture: logic layers, backend, and the front end. The prediction is done using the models written in python. The backend was written in the FastAPI python library for integration of the deep-learning models with the webserver. The front end and user interface were built using JavaScript and the JavaScript library Bootstrap. The server is hosted using Google Cloud Platform (GCP).

##### 1.6. Statistics

Reported values were the average of 10 independent validation replicates, each with its own train, test, and validation set. Error is reported as ± standard deviation (SD) of these replicates. *P* Values were calculated by a two-tailed Student's t-test using a paired test in Microsoft Excel or the Scipy library in python. In the case of Cross models, since the data was no longer paired, an unpaired t-test with unequal variance was used. A cutoff of $P < 0.05$ was used for significance.

#### 2. Dataset Preparation

##### 2.1. MusiteDeep Datasets

The datasets used by MusiteDeep [1] were downloaded from musite.net. These datasets consisted of a FASTA file with amino acids that had the PTM followed by a "#". We then generated a file of the Uniprot ID of the sequence in one column with all of the modified site locations in the next column, along with a FASTA file without the "#", which was used as the reference sequence file for the all-organism models.

##### 2.2. OGP Datasets

The dataset from the OGP[2] was downloaded from http://www.oglyp.org/download.php (accessed 7 February 2024). To access the entire database (specifically for information on HexNAc versus GalNAc glycans), we contacted the authors directly. For the whole glycosylation datasets, the datasets downloaded were already a file with the UniProt ID with the site location. For the HexNAc and GalNAc datasets, we split the whole database (which includes that information and less validated sites) into subsets of sites with HexNAc or GalNAc. These were then cross-referenced with the main set to only include validated sites.

##### 2.3. O-GlcNAc Site Atlas Datasets

The dataset from O-GlcNAcAtlas [3] was downloaded from https://oglcnac.org/atlas/download/ (accessed: 12 February 2024). Only the Unambiguous sites dataset was used, and the dataset downloaded was already a file with the UniProt ID with the site location. This dataset had duplicates (same Uniprot ID and site location), so k-mers were only added to the set of positive k-mers if they did not already exist in the set of positive k-mers.

##### 2.4. LMGglyPred Datasets

The datasets from LMNglyPred[4] were downloaded from their Github at https://github.com/KCLabMTU/LMNglyPred (accessed 2 February 2024). These datasets included both the NGlycosite Atlas[5] and NGlyDE[6] sets. The datasets downloaded were already a file with the UniProt ID having the site location. Given that the NGlyDE dataset was already for human proteins and the NGlycosite Atlas dataset was for all organisms, we made a human protein model only from the NGlyDE dataset and an all-organism model from the NGlycosite dataset. For the NGlycosite dataset, there were many isoforms, so all of the isoform sequences on Uniprot were downloaded, and isoforms in the dataset were added to the reference sequences.

Because this set already had negatives in a similar ratio to positives, no negatives were generated as in the other datasets, additional balancing of the ratio of negatives to positives was performed. These datasets were already split into test and train sets. For the first replicate, this split was preserved, but for all subsequent validation replicates, the sets were combined, shuffled, and split again.

##### 2.5. Sugiyama Kinase Datasets

For the datasets from Sugiyama [7] the data were downloaded from https://www.nature.com/articles/s41598-019-46385-4#Sec21 (accessed, 29 November 2023). These data included the kinase and its potential substrates and the modified sites. These data were then imported into Python, and the number of sites per kinase were tabulated. A set of kinases with over 500 sites was generated, as a large number of sites is necessary for successful training and there were no S/T kinases with over 1000 sites. From this set of kinases with over 500 sites, the kinases, which mainly phosphorylated at tyrosine (using a cutoff of 50% of the sites), were then removed from the set, resulting in a set of 68 kinases that phosphorylated primarily at S/T. In the remaining kinase datasets, all of the tyrosine sites were then removed.

Given that all of the kinases were human, the reference proteome UP000005640 was used to get the sequences and generate negatives. Once the 68 kinase datasets were prepared, the training, test, and validation sets in 10-fold were prepared for each kinase, as in Section 1.

#### 3. Frequency of Nearby PTM Sites

##### 3.1. Generation of PTM frequency graphs

To determine how often PTMs appeared next to other PTMs, we pooled all of the positive k-mers for a given PTM and tabulated the pairwise distances. These distances are represented as frequency graphs from the prediction amino acid (k-mer center).

To capture every pairwise distance between PTMs only once, we considered only PTMs that were C-terminal to the k-mer center, thereby avoiding double-counting of residues N-terminal to the k-mer center.

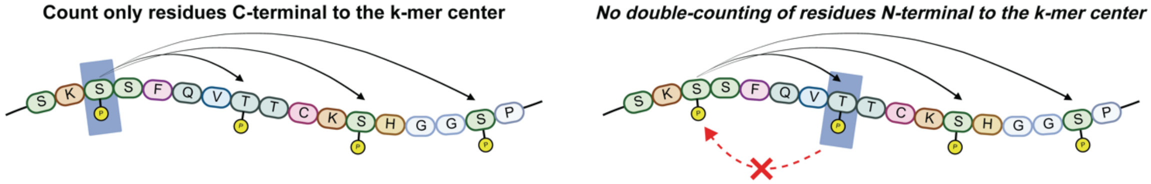

#### 3.2. Frequency of nearby PTM sites in human-only datasets taken from MusiteDeep

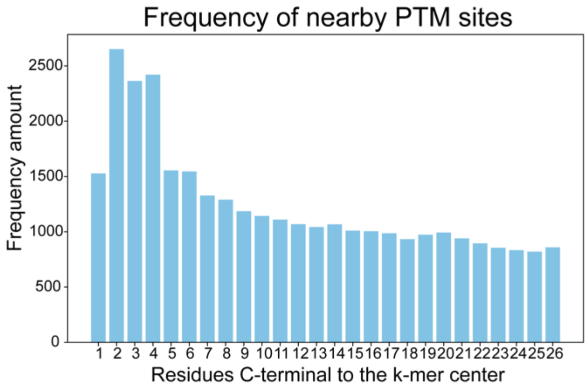


Phosphorylation (S,T)

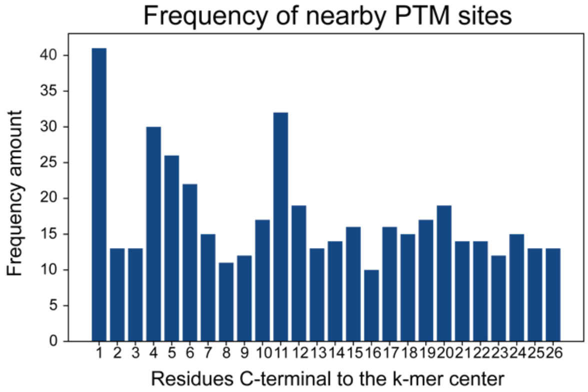


Phosphorylation (Y)

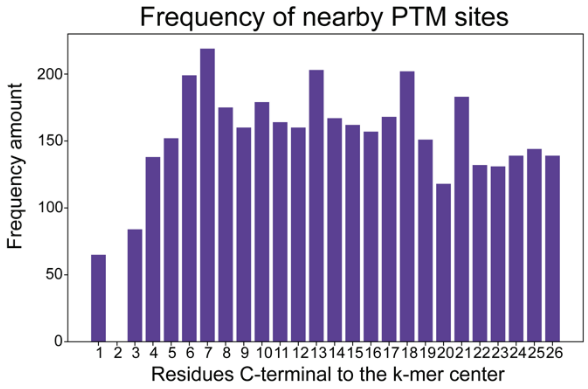


N-Glycosylation (N)

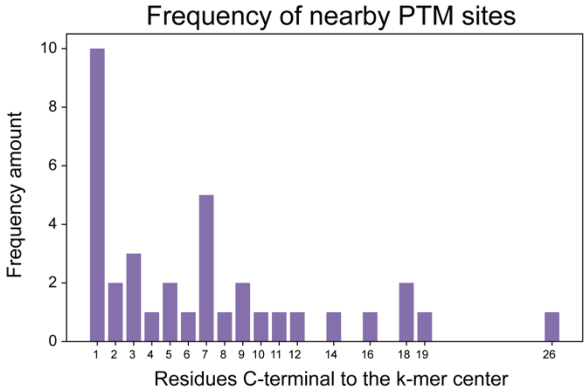


O-Glycosylation (S,T)

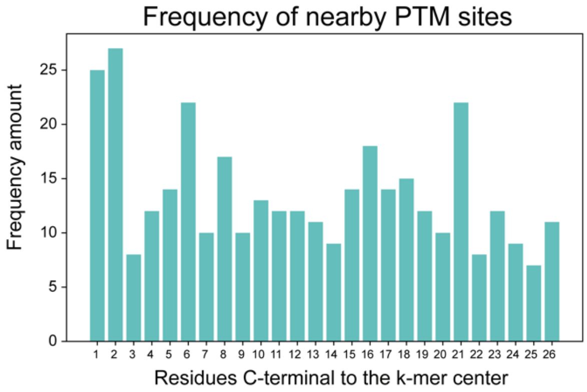


Ubiquitination (K)

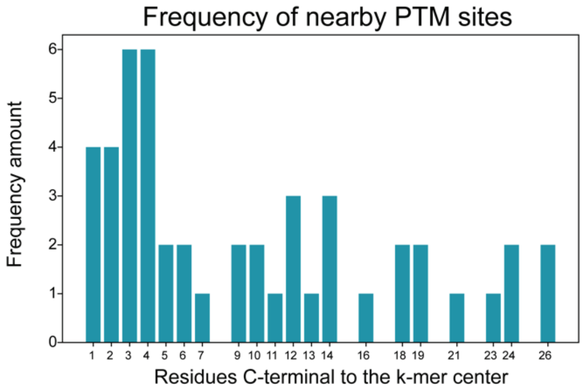


SUMOylation (K)

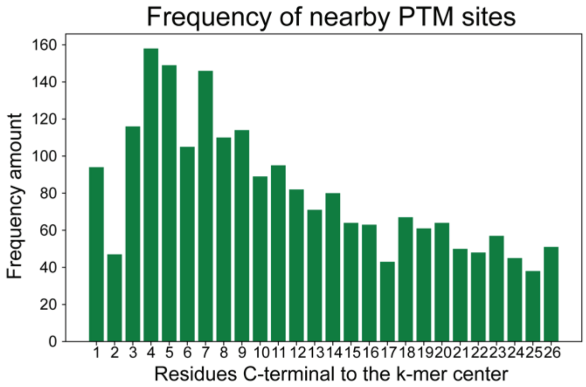


Acetylation (K)

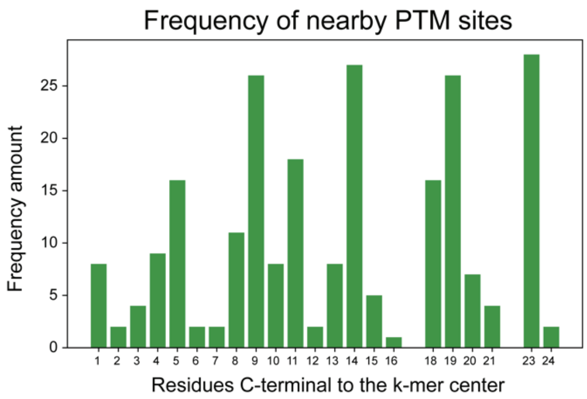


Methylation (K)

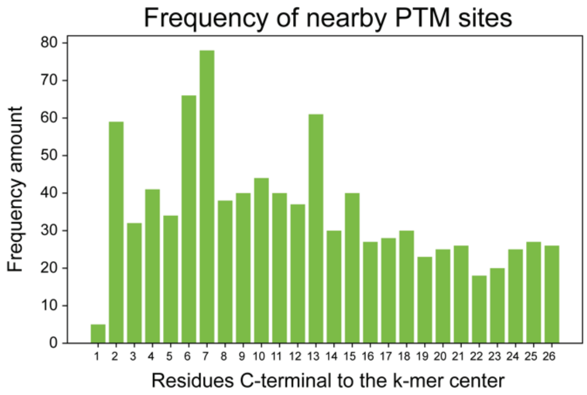


Methylation (R)

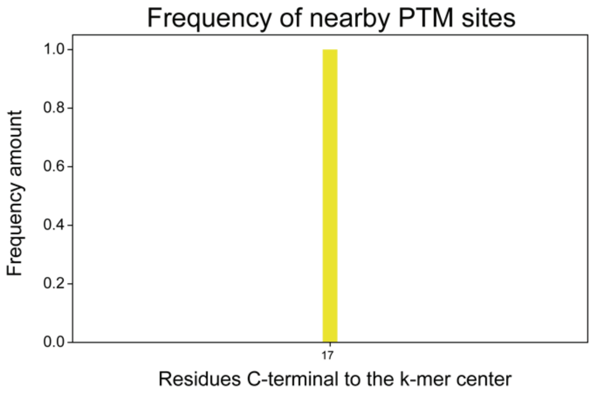


Pyroglutamylation (Q)

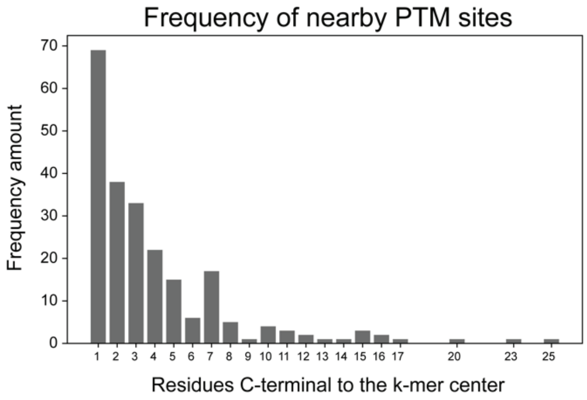


Palmitoylation (C)

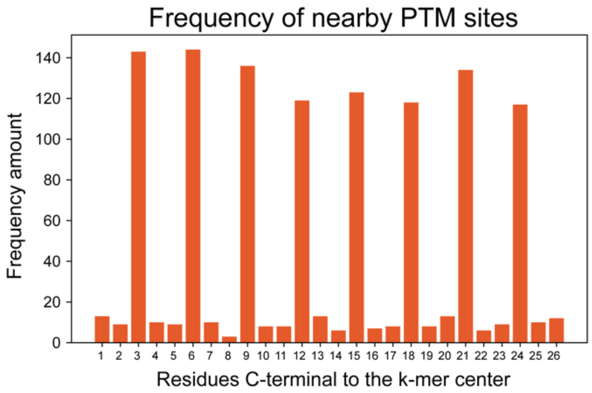


Hydroxylation (P)

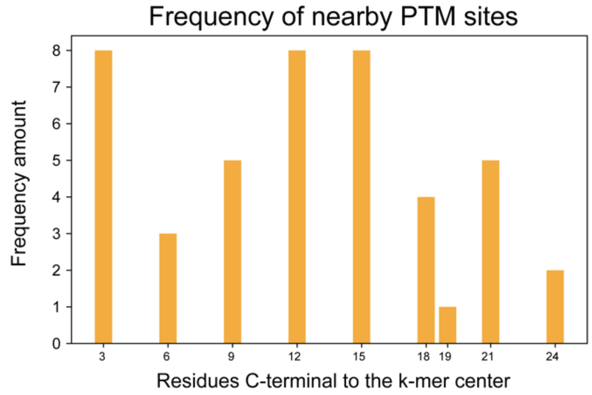


Hydroxylation (K)

##### 3.3. Frequency of Other Sites in All-Organisms from MusiteDeep

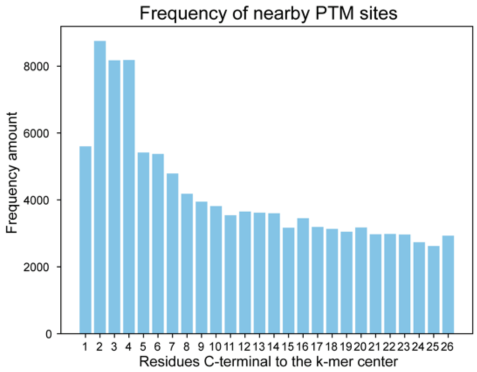


Phosphorylation (S,T)

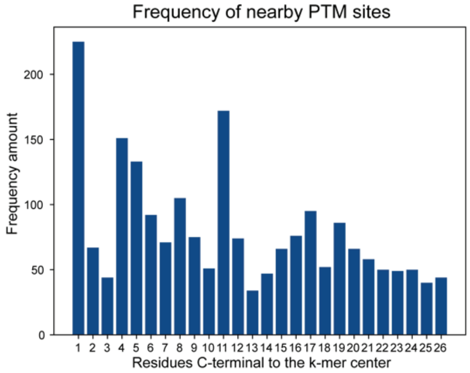


Phosphorylation (Y)

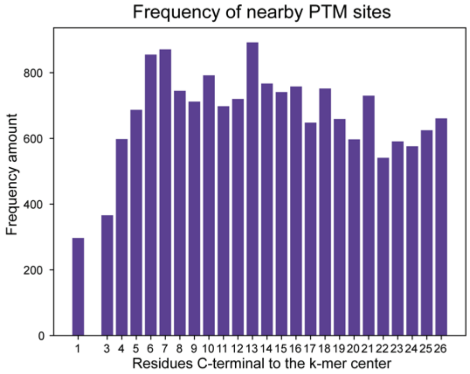


N-Glycosylation (N)

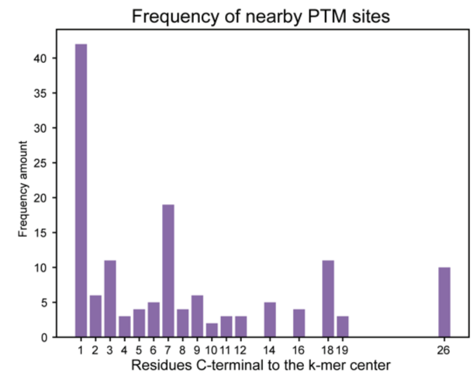


Glycosylation (O)

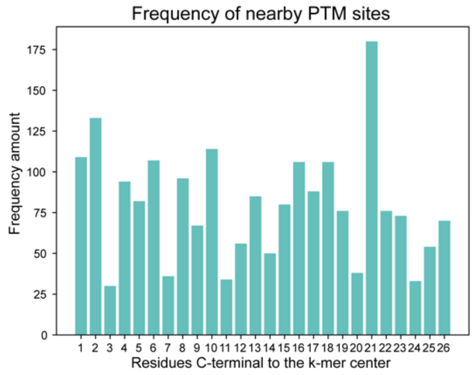


Ubiquitination (K)

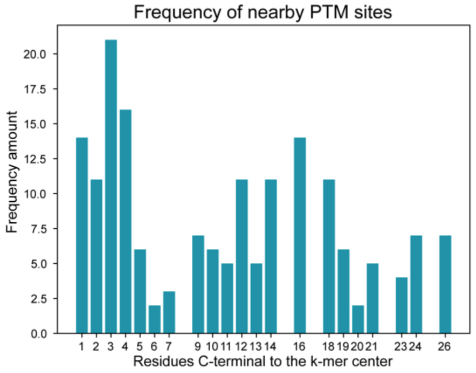


SUMOylation (K)

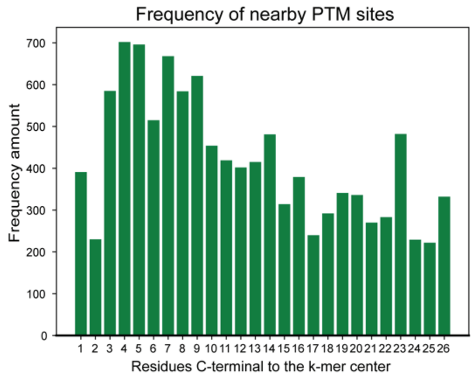


Acetylation (K)

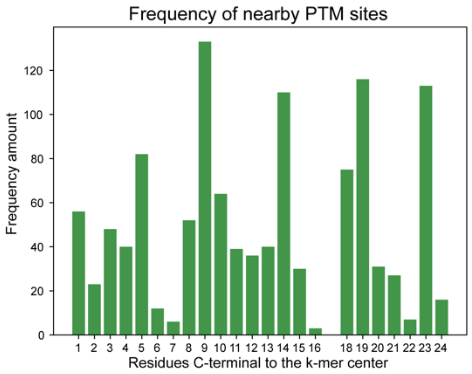


Methylation (K)

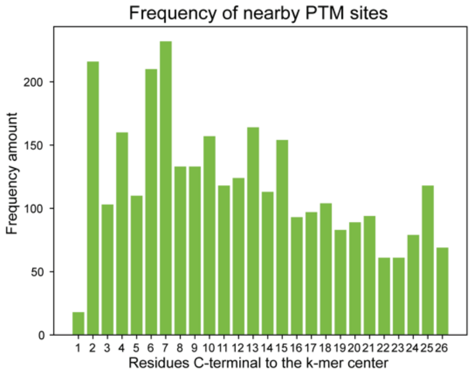


Methylation (R)

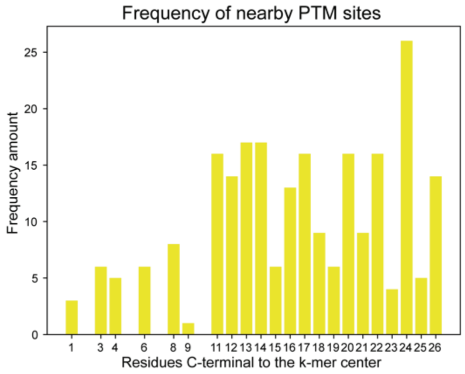


Pyroglutamylation (Q)

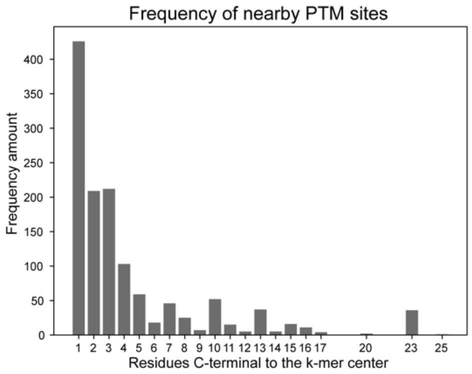


Palmitoylation (C)

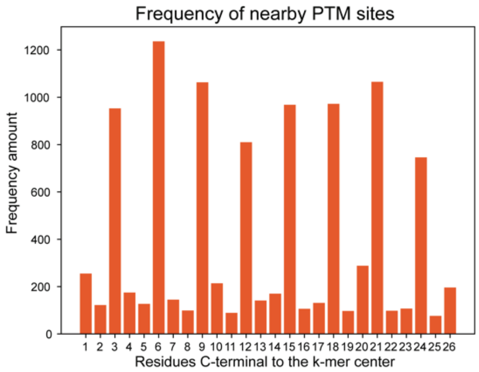


Hydroxylation (P)

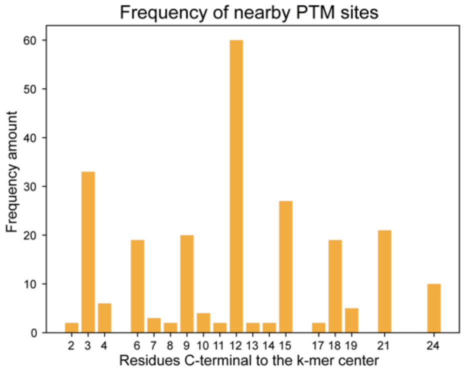


Hydroxylation (K)

##### 3.4. Frequency of Other O-Glycosylation Sites Taken from OGP and O-GlcNAc Site Atlas

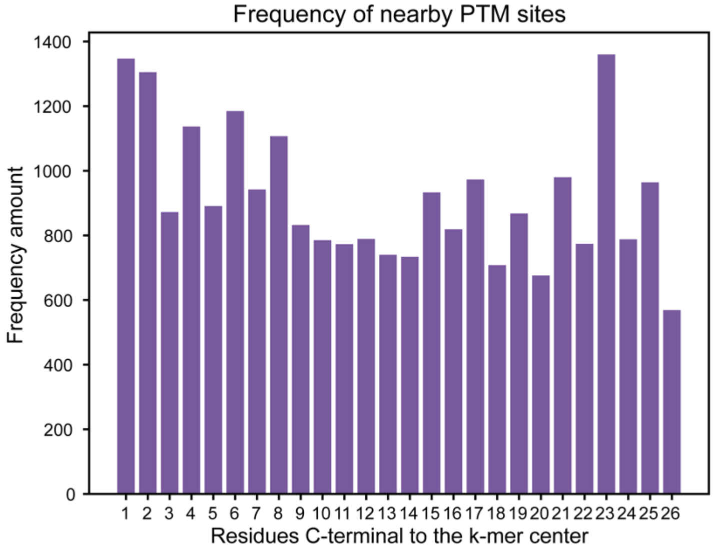


OGP O-glycosylation
Human-only proteins

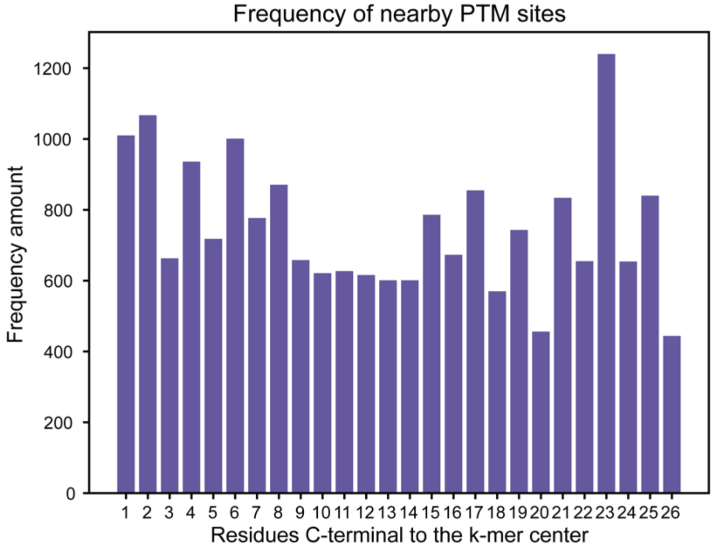


OGP GalNAc
Human-only proteins

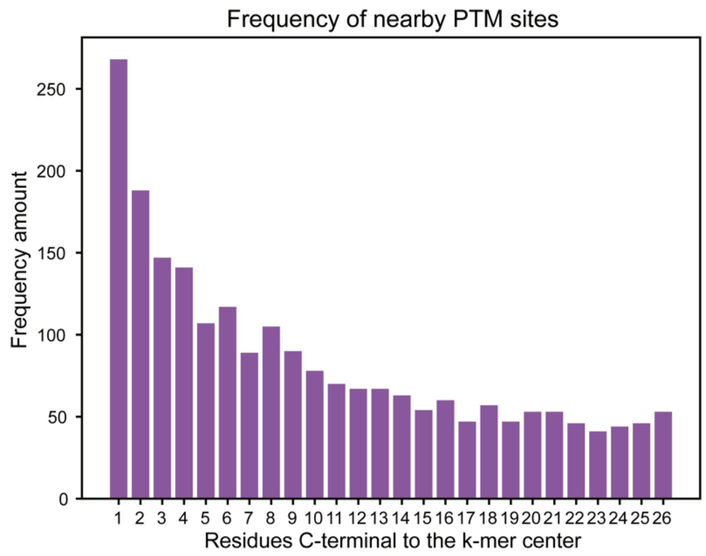


OGP HexNAc
Human-only proteins

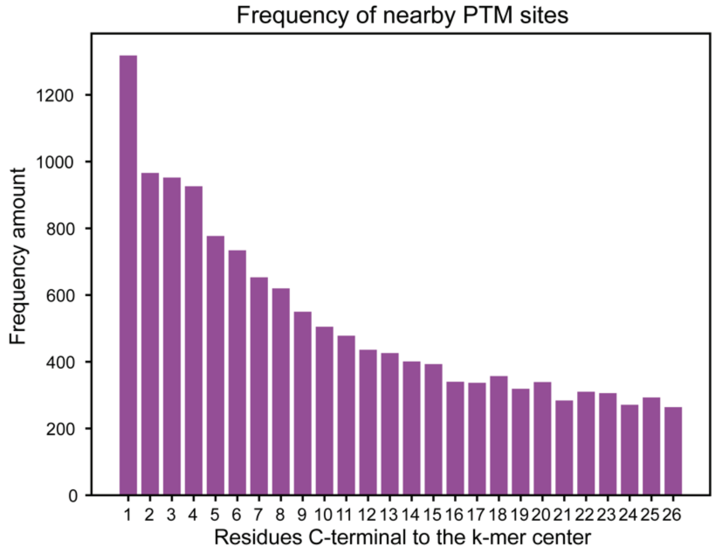


O-GlcNAc Site Atlas
Human-only proteins

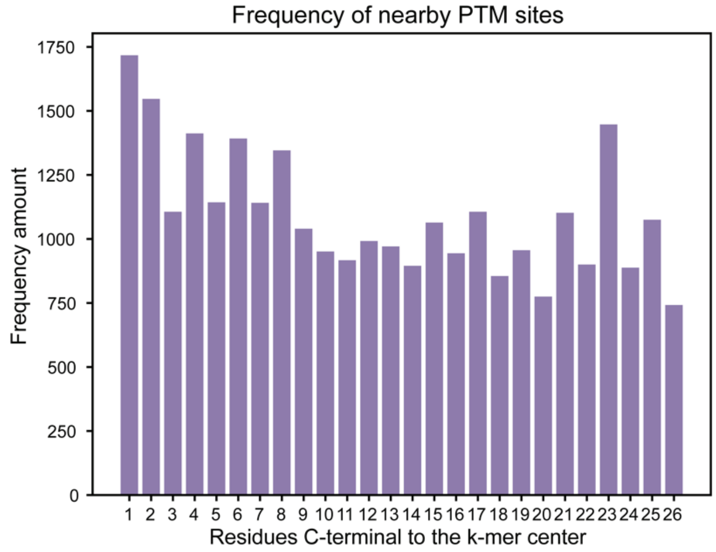


OGP O-glycosylation
All-organism proteins

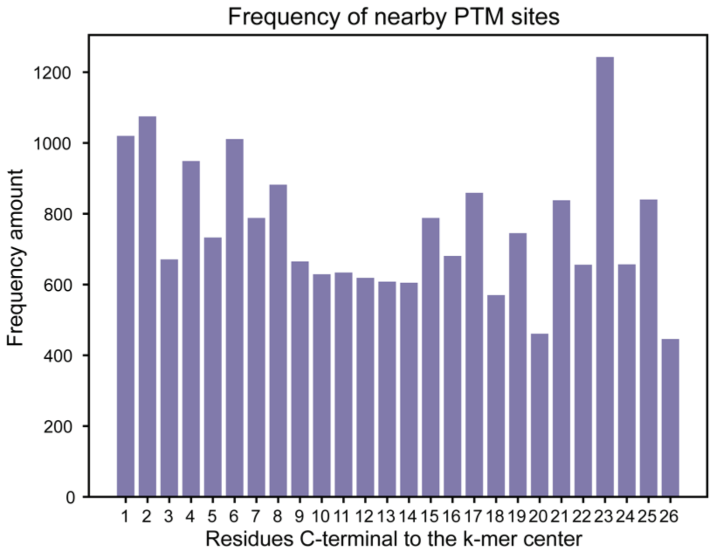


OGP GalNAc
All-organism proteins

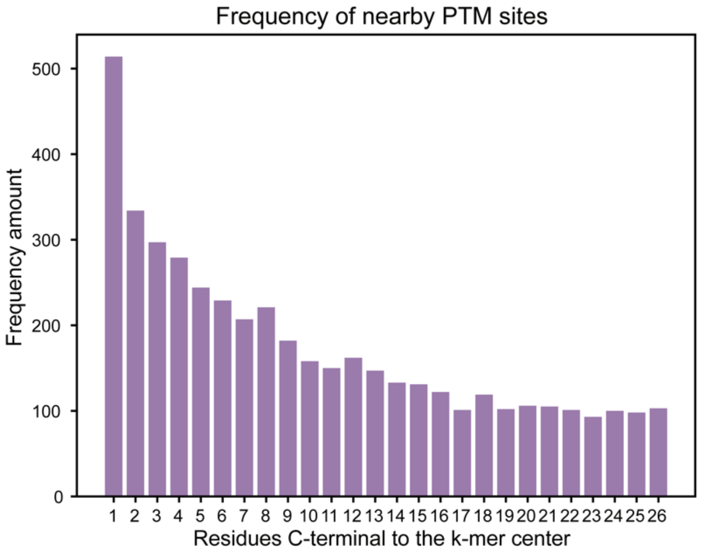


OGP HexNAc
All-organism proteins

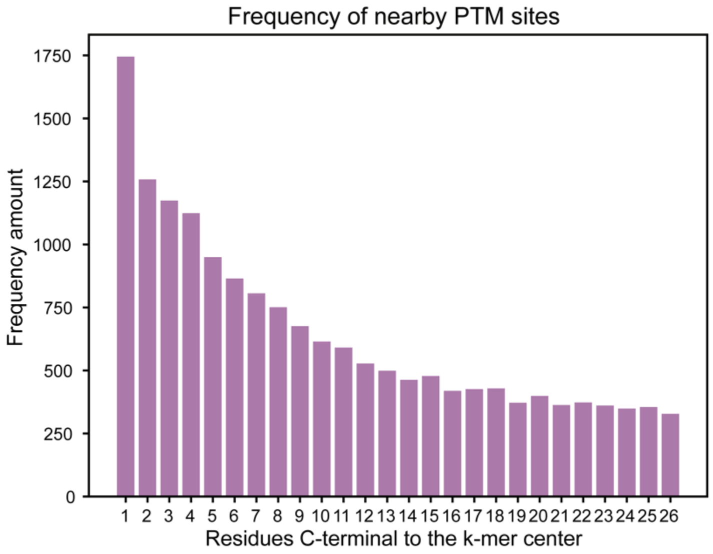


O-GlcNAc Site Atlas
All-organism proteins

##### 3.5. Frequency of Other N-Glycosylation Sites from LMNglyPred Datasets

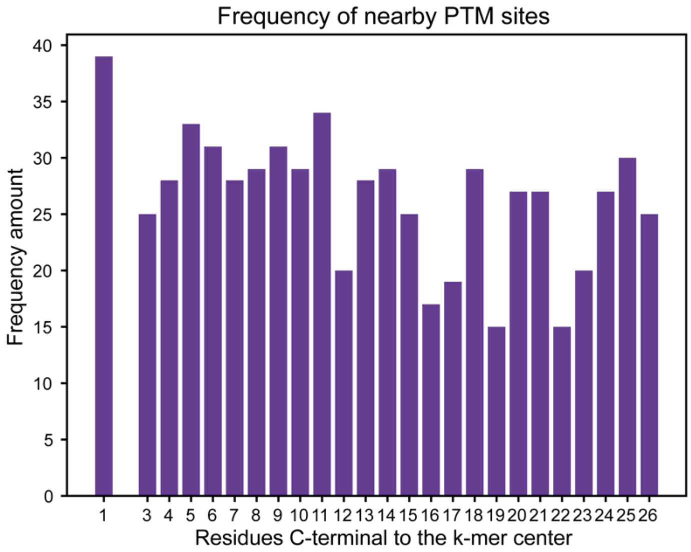


N-Glycosite Atlas dataset

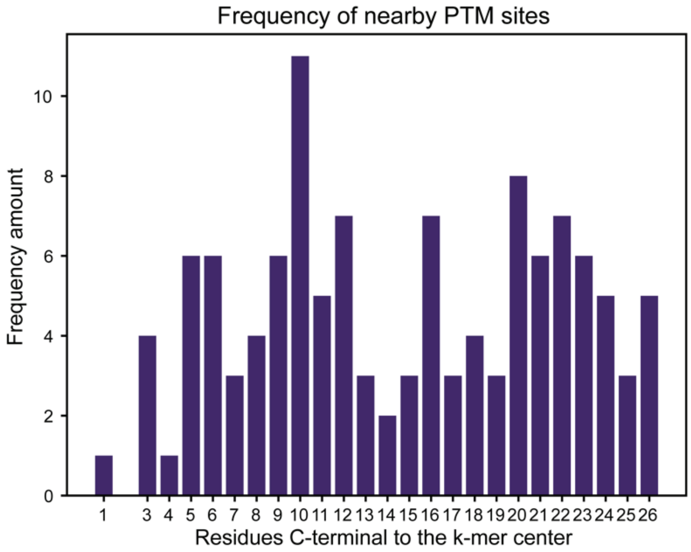


NGlyDE dataset

##### 3.6. Frequency of Other S/T Phosphorylation Sites from Sugiyama Kinase Datasets

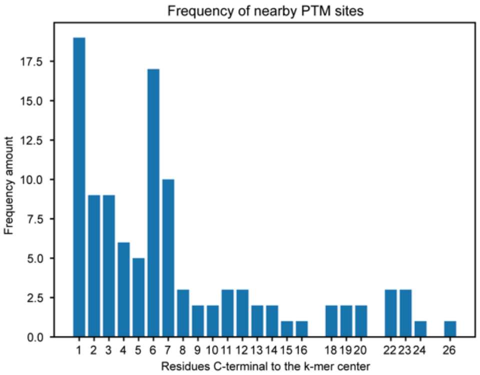


AurA

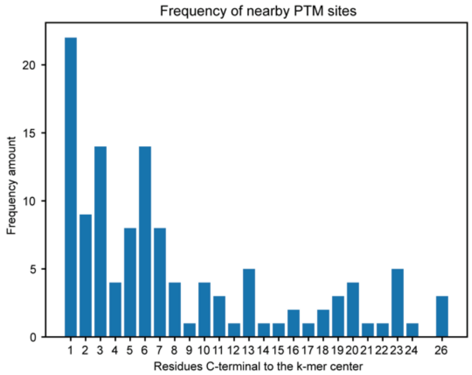


AurC

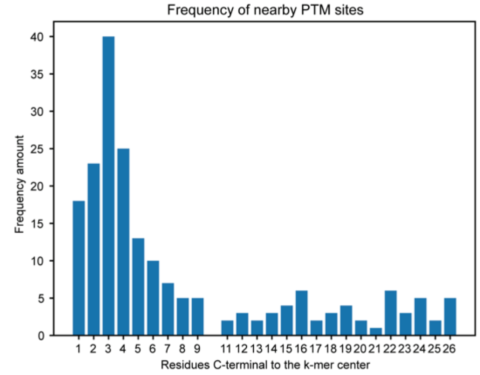


CaMK2δ

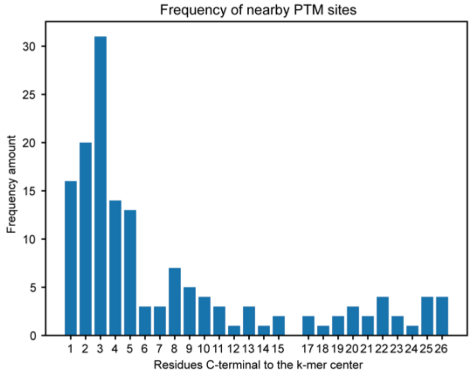


CaMK2γ

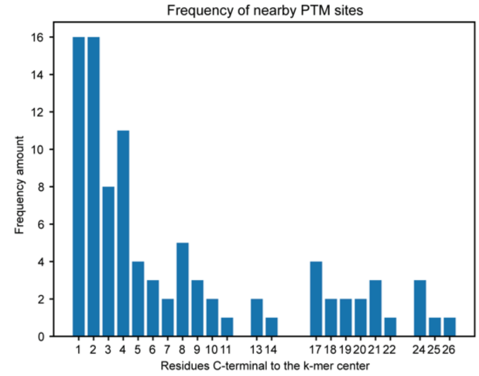


CDC2/CycB1

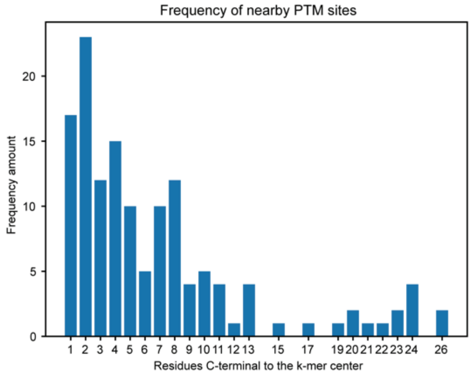


CDK5/p25

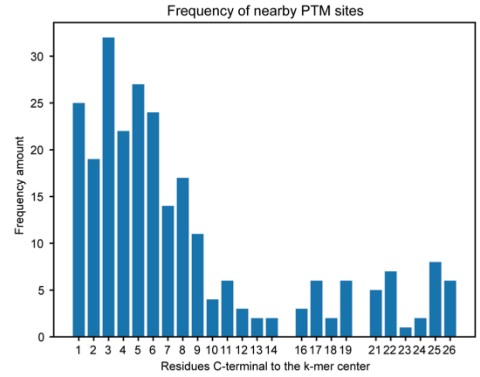


CHK1

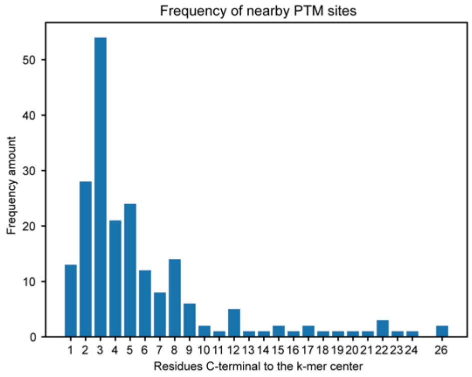


CK1δ

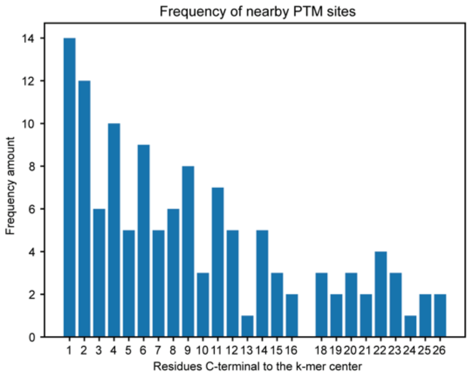


DCLK2

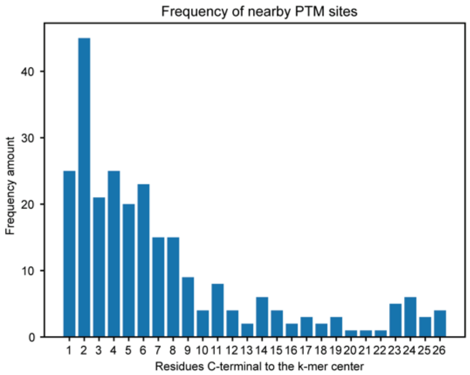


DYRK2

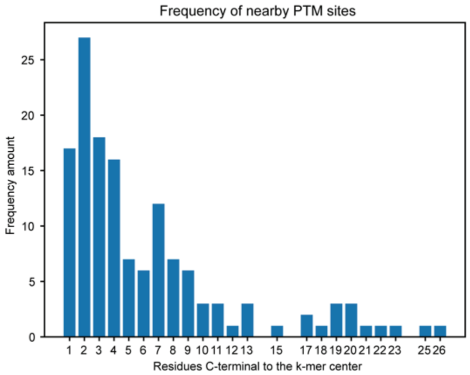


DYRK3

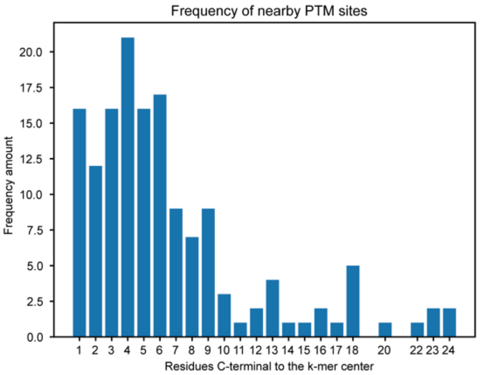


Erk1

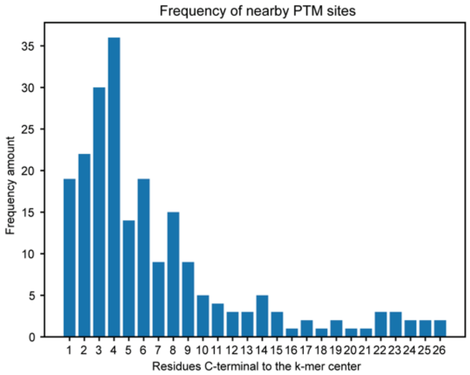


Erk2

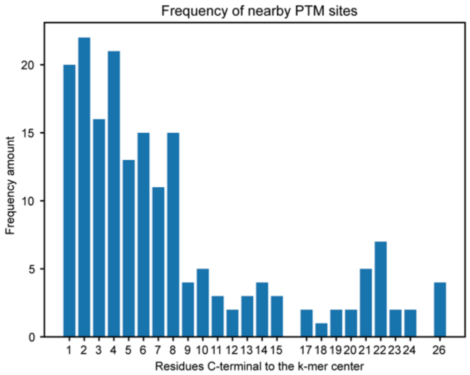


Erk5

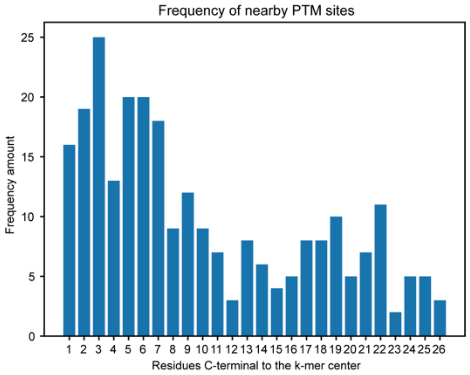


GCK

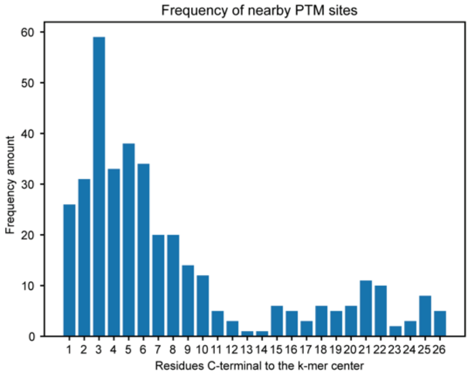


HGK

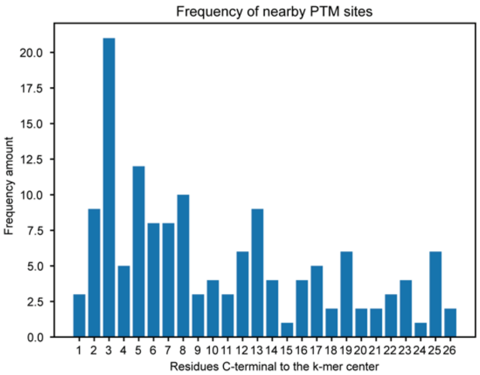


IKKα

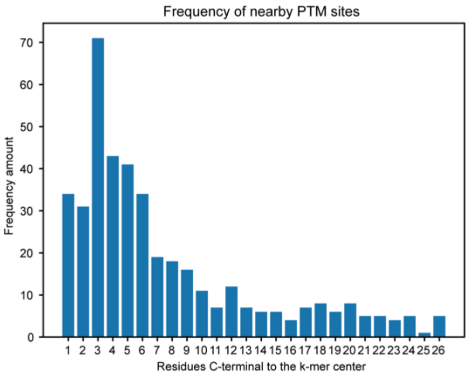


IKKβ

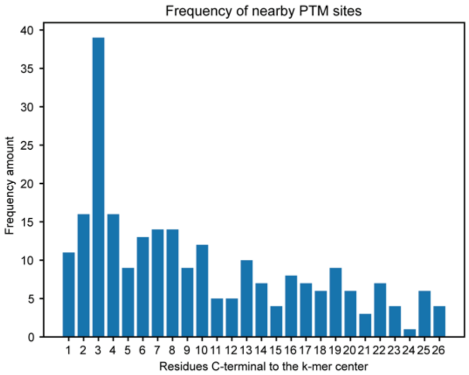


IKKε

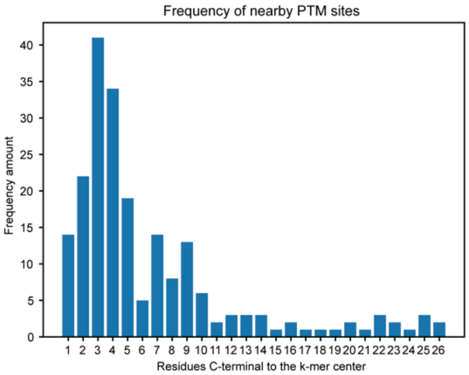


JNK1

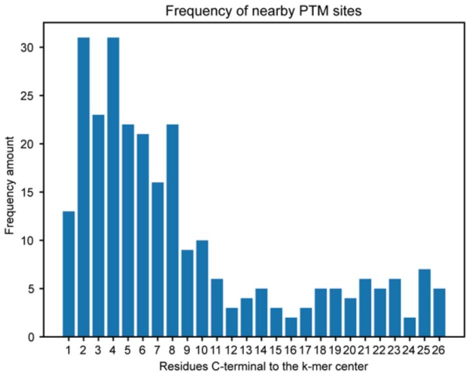


KHS1

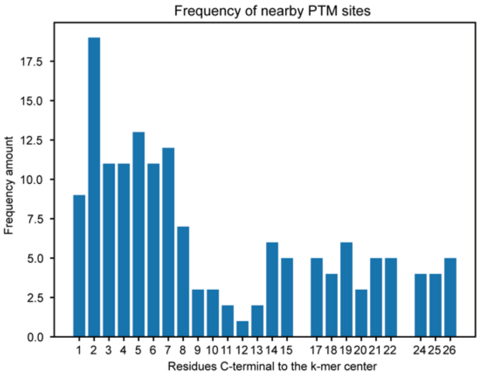


MAP3K1

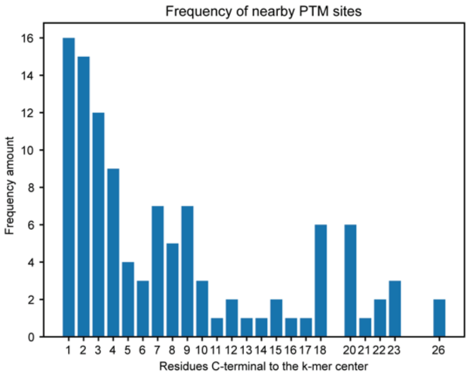


MAPKAPK3

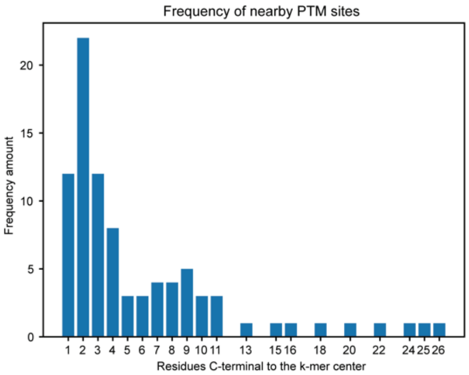


MAPKAPK5

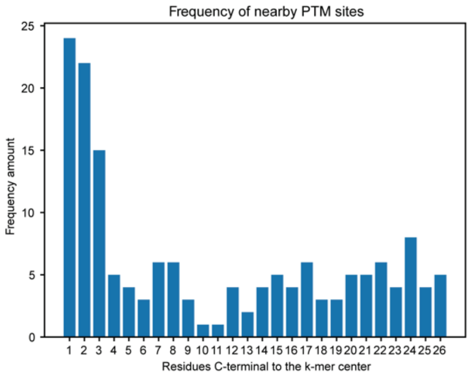


MARK2

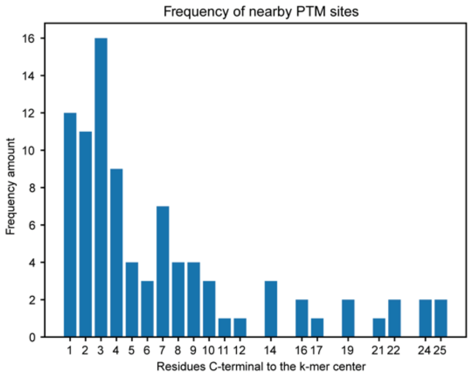


MARK4

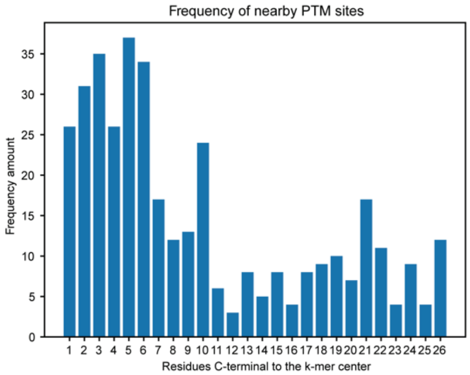


MINK

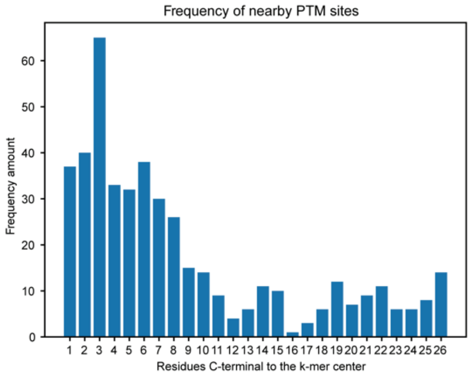


MLK1

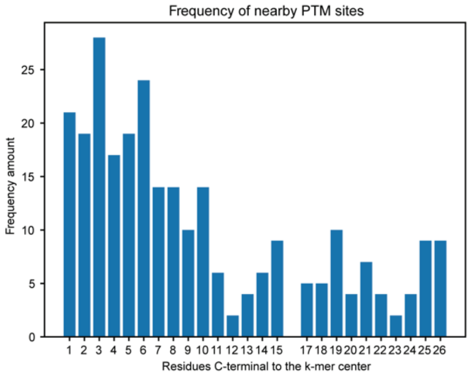


MLK2

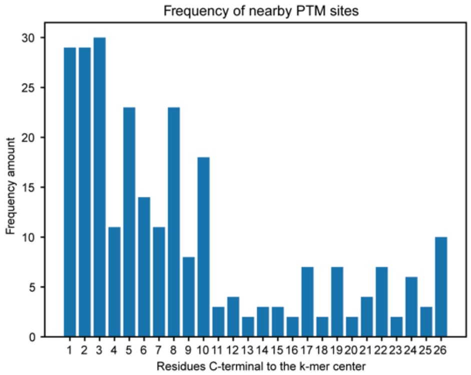


MLK3

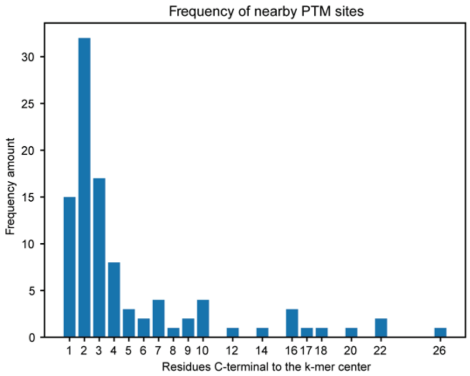


MSK1

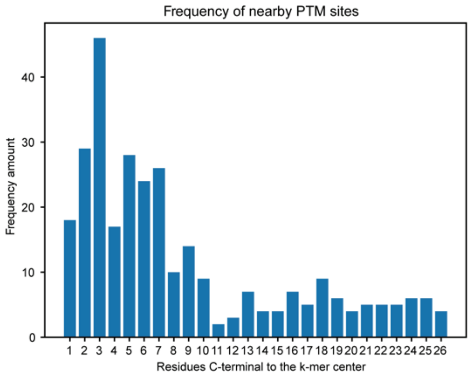


MST1

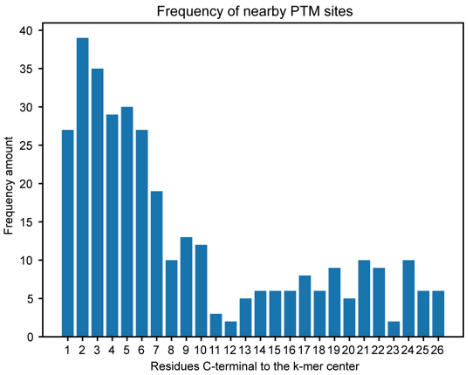


MST2

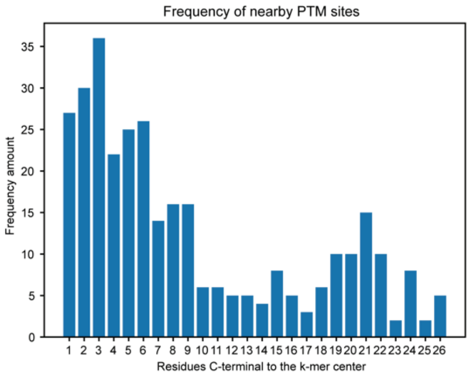


MST3

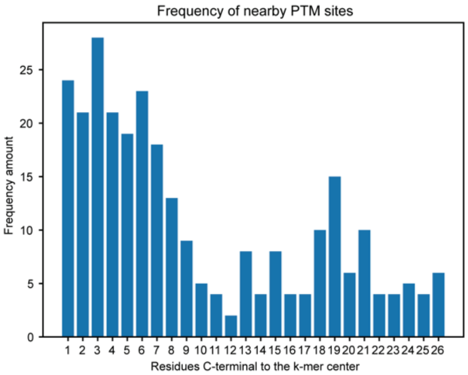


MST4

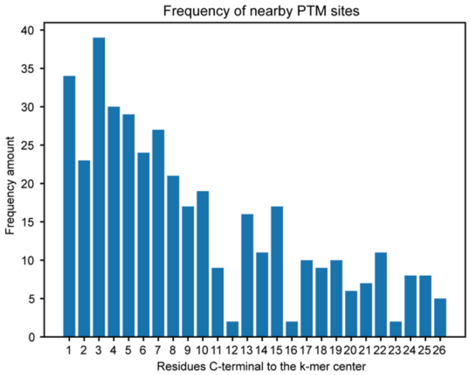


NEK1

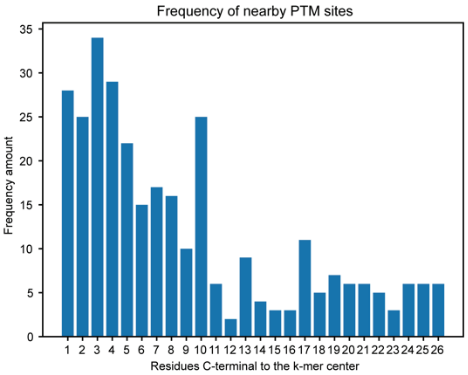


NEK2

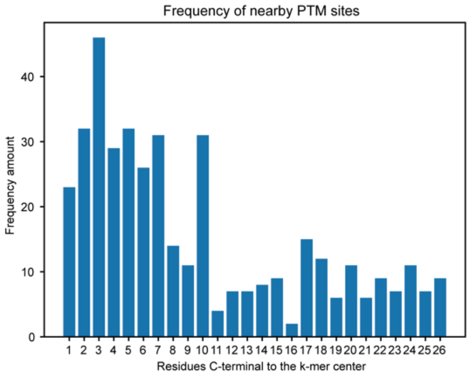


NEK4

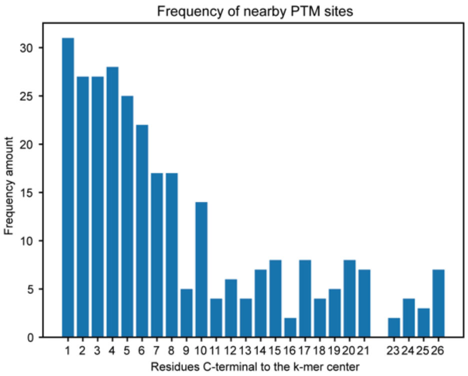


NEK6

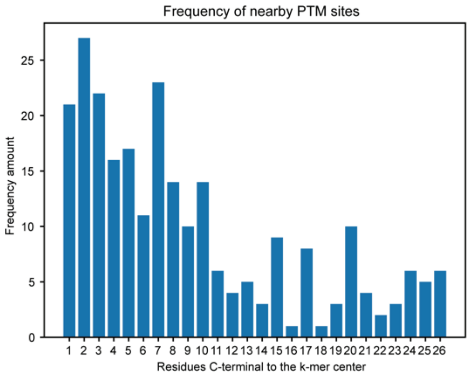


NEK7

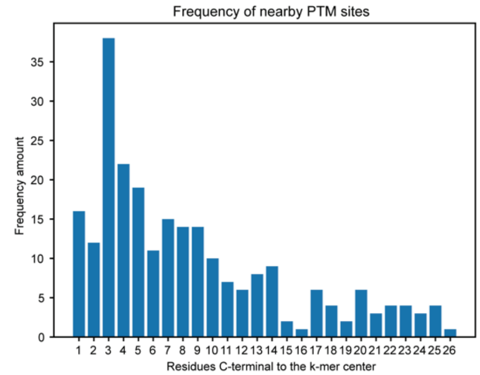


NEK9

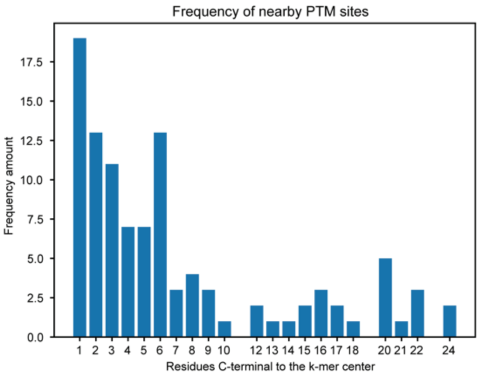


NIM1

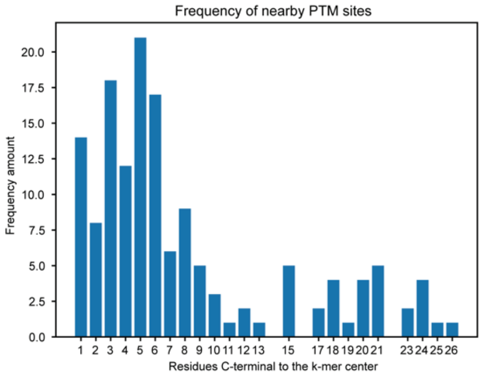


NLK

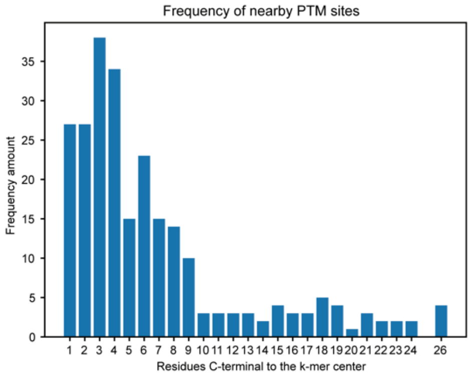


p38α

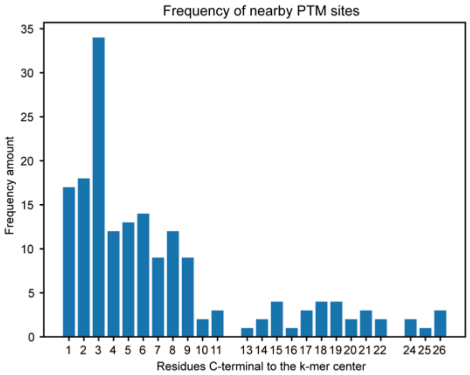


p38β

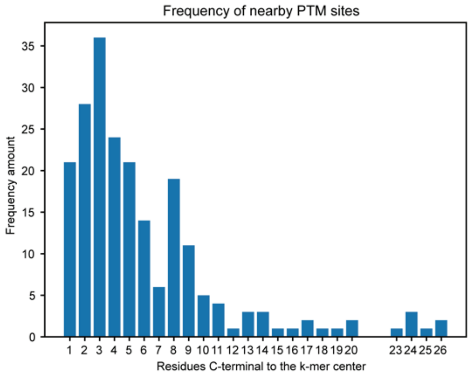


p38δ

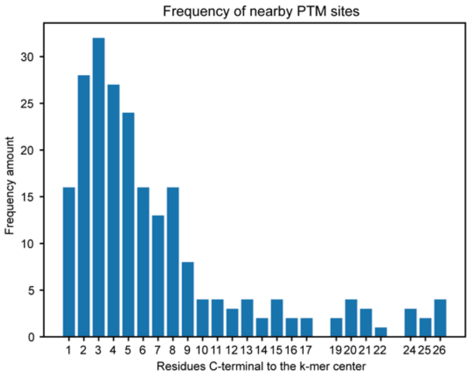


p38γ

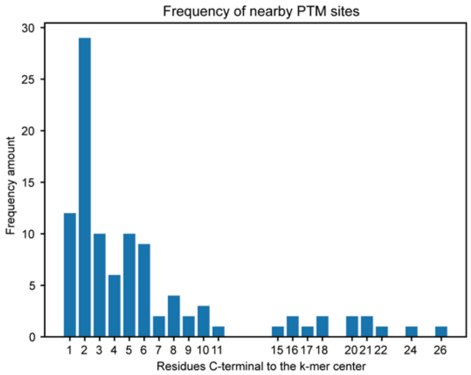


p70S6K

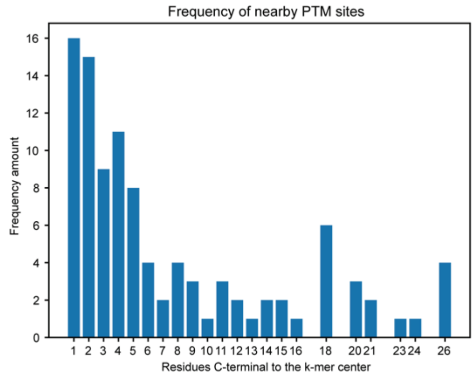


PAK1

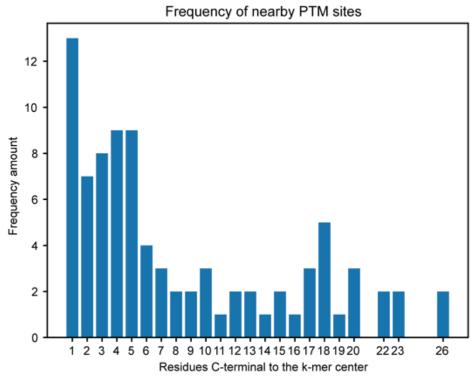


PAK2

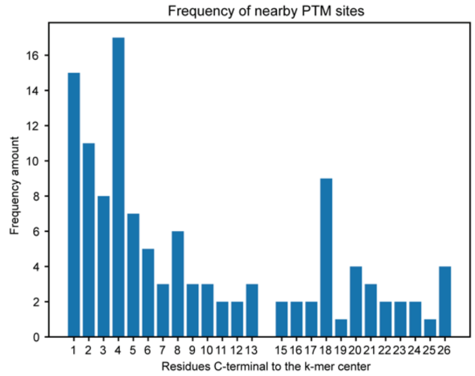


PAK3

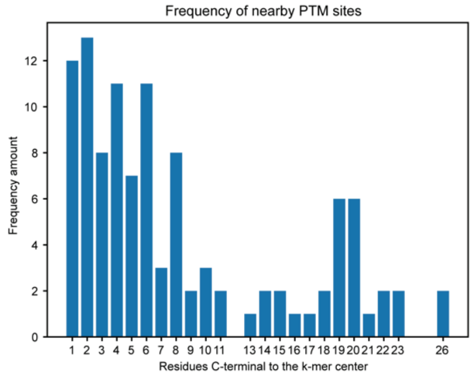


PHKG2

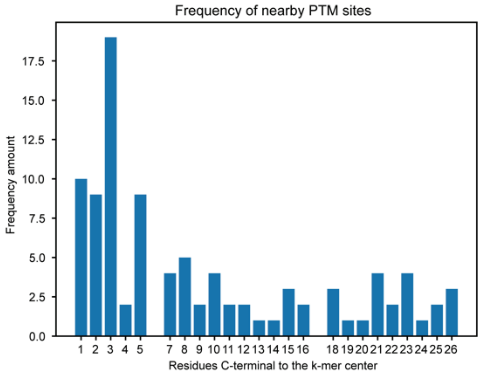


PIM1

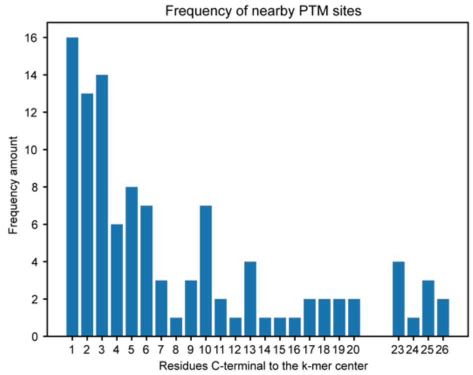


PIM3

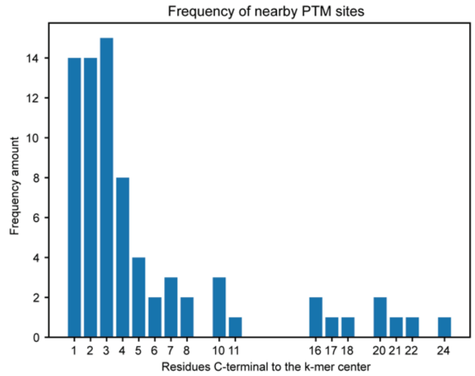


PKA Cα

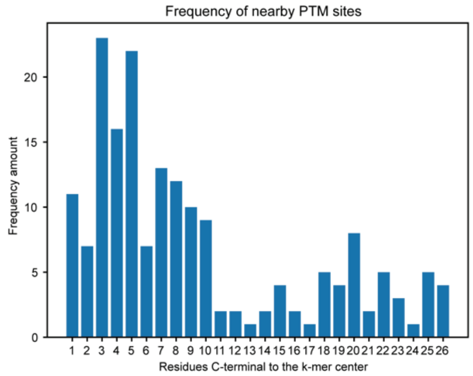


PLK1

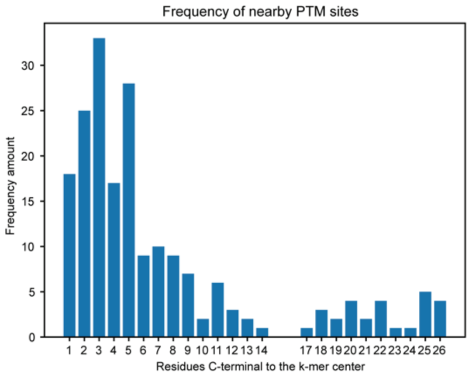


PLK3

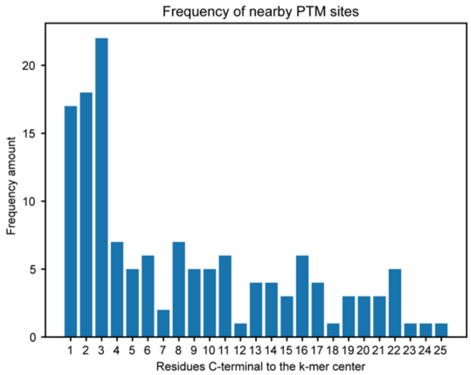


QIK

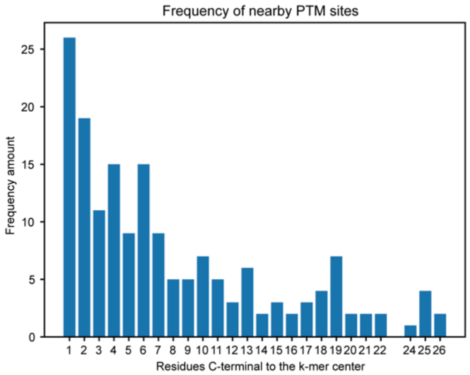


RSK3

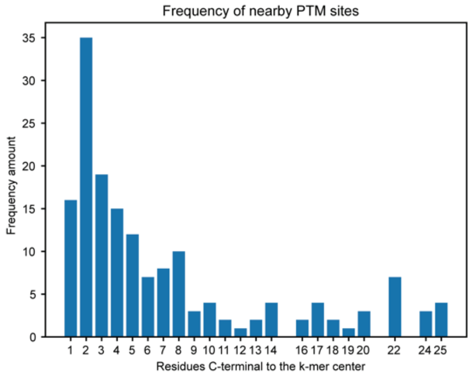


S9K3

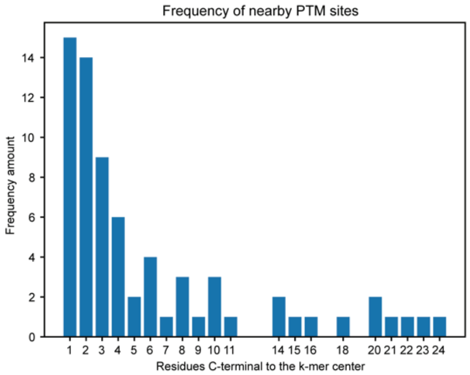


SIK

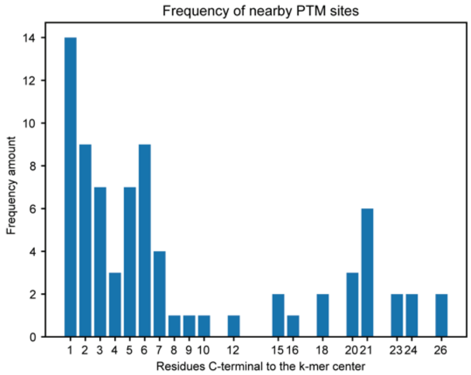


TAK1

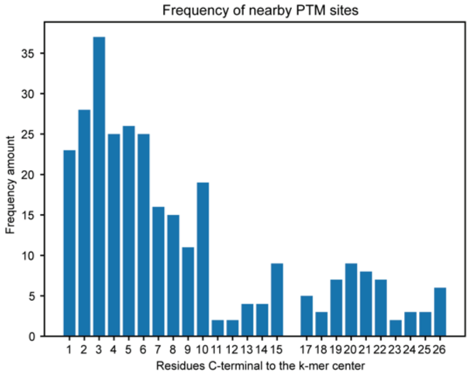


TAO2

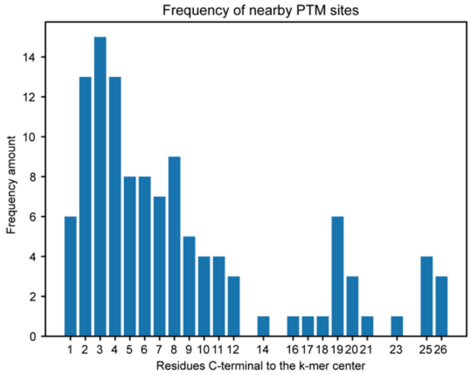


TLK1

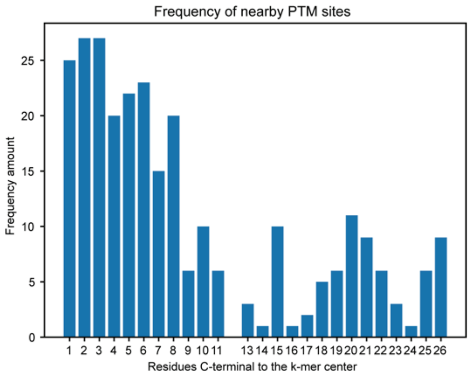


TNIK

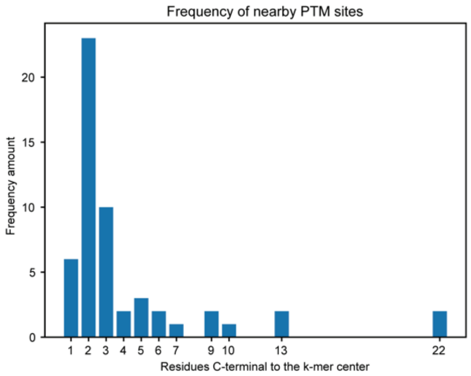


TSSK1

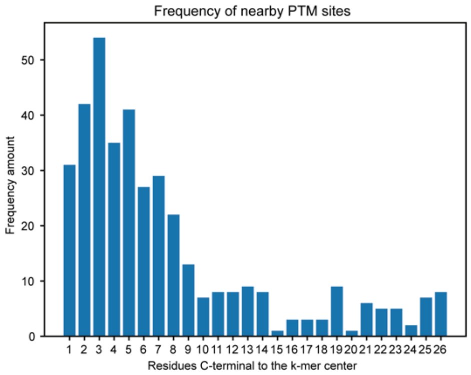


TTBK1

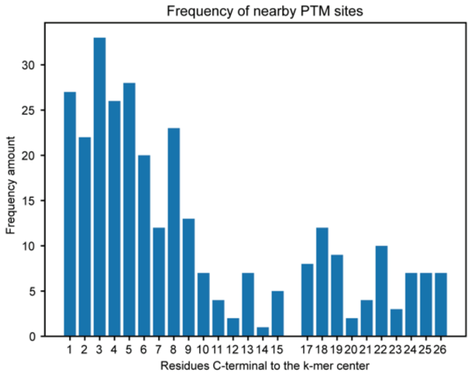


YSK1

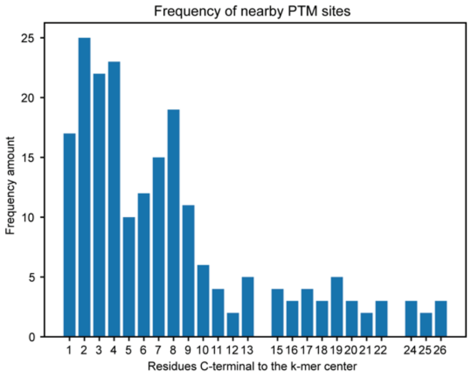


ZAK

#### 4. Tuning of Model Parameters

##### 4.1. Determination of Optimal "k" for k-mers

To ensure the best possible performance for our model, we tuned deep learning hyperparameters such as the learning rate, batch size, and epoch. We also tested different k-mer lengths which resulted in an optimal value of k = 53 that results in a high accuracy and reasonable size for running in models.

| k-mer length | Accuracy |
|---|---|
| 21 | 0.8430 |
| 31 | 0.8522 |
| 41 | 0.8583 |
| 51 | 0.8575 |
| 53 | 0.8624 |
| 55 | 0.8587 |
| 61 | 0.8610 |
| 71 | 0.8654 |

##### 4.2. Testing of Other Model Architectures

Different deep learning model architectures were tested for performance and run time. We found the best to be the embedding CNN model, which not only produces the highest accuracy for phosphorylation and lysine acetylation but is also a relatively simple model that runs on the order of minutes rather than hours as the LSTM and Attention models tend to do.

| Model | Embedding CNN | LSTM | Attention | OneHot Encoding CNN |
|---|---|---|---|---|
| Phosphorylation with labels | 0.8624 | 0.8591 | 0.8371 | 0.8563 |
| Phosphorylation without labels | 0.8083 | 0.8115 | 0.7848 | 0.7899 |
| N-Glycosylation with labels | 0.9534 | 0.9543 | 0.9315 | 0.9479 |
| N-Glycosylation without labels | 0.9527 | 0.9440 | 0.9353 | 0.9485 |
| Lysine acetylation with labels | 0.7870 | 0.7811 | 0.6757 | 0.7527 |
| Lysine acetylation without labels | 0.7243 | 0.7420 | 0.6260 | 0.6769 |

##### 4.3. Hyperparameter Tuning

Further tuning was done with the hyperparameters learning rate, dropout rate, L2 rate, batch size, and kernel size. We tried learning rates of 0.1, 0.01, 1E−03, and 1E−04. We tried dropout rates of 0.9, 0.5, 0.2, and 0.1. We tried L2 rates of 1E−03, 1E−06, and 0. We tried batch sizes of 10, 30, 50, and 100. We tried kernel sizes of 3×3 and 5×5. We tested all possible combinations of these hyperparameters with a small k-mer size of 35 to have a reasonable run time. Our best results were a learning rate of 1E−03, a dropout rate of 0.1, and L2 rate of 1E−06, a batch size of 50, and a kernel size of 3×3, allowing our accuracy to improve from 79% to 85%. In our actual model, we have changed the learning rate to 1E−04 and the batch size to 256.

### 5. Additional Model Results

#### 5.1. Model results in human-only datasets taken from MusiteDeep

| PTM | | | AUC | AUPRC | Accuracy | Recall | Precision | MCC | F1 | Specificity |
|---|---|---|---|---|---|---|---|---|---|---|
| **Phosphorylation (S,T)** | No labels | Avg | 0.8790 | 0.8709 | 0.7963 | 0.7897 | 0.8029 | 0.5932 | 0.7959 | 0.8031 |
| | | STD | 0.0062 | 0.0069 | 0.0074 | 0.0246 | 0.0133 | 0.0147 | 0.0101 | 0.0193 |
| | Labels | Avg | 0.9301 | 0.9304 | 0.8575 | 0.8505 | 0.8646 | 0.7154 | 0.8573 | 0.8648 |
| | | STD | 0.0015 | 0.0018 | 0.0028 | 0.0121 | 0.0102 | 0.0056 | 0.0031 | 0.0129 |
| | | *P* value | 3.51E−10 | 1.75E−10 | 5.65E−11 | 1.51E−05 | 3.87E−09 | 4.57E−11 | 3.56E−09 | 6.59E−07 |
| **Phosphorylation (Y)** | No labels | Avg | 0.8030 | 0.7888 | 0.7250 | 0.7116 | 0.7331 | 0.4507 | 0.7207 | 0.7377 |
| | | STD | 0.0150 | 0.0320 | 0.0153 | 0.0460 | 0.0339 | 0.0317 | 0.0236 | 0.0499 |
| | Labels | Avg | 0.8368 | 0.8533 | 0.7538 | 0.7243 | 0.7706 | 0.5100 | 0.7453 | 0.7846 |
| | | STD | 0.0187 | 0.0275 | 0.0191 | 0.0481 | 0.0398 | 0.0375 | 0.0306 | 0.0301 |
| | | *P* value | 0.0006 | 0.0002 | 0.0026 | 0.5725 | 0.0452 | 0.0020 | 0.0728 | 0.0292 |
| **N-Glycosylation (N)** | No labels | Avg | 0.9681 | 0.9453 | 0.9446 | 0.9856 | 0.9103 | 0.8922 | 0.9464 | 0.9040 |
| | | STD | 0.0043 | 0.0088 | 0.0039 | 0.0040 | 0.0073 | 0.0074 | 0.0039 | 0.0075 |
| | Labels | Avg | 0.9806 | 0.9751 | 0.9480 | 0.9840 | 0.9173 | 0.8984 | 0.9495 | 0.9124 |
| | | STD | 0.0028 | 0.0045 | 0.0038 | 0.0037 | 0.0075 | 0.0071 | 0.0039 | 0.0076 |
| | | *P* value | 2.46E−06 | 4.45E−07 | 7.54E−02 | 3.94E−01 | 6.00E−02 | 8.75E−02 | 1.12E−01 | 3.15E−02 |
| **O-Glycosylation (S,T)** | No labels | Avg | 0.7035 | 0.7634 | 0.6591 | 0.8471 | 0.6527 | 0.2695 | 0.7309 | 0.4095 |
| | | STD | 0.1014 | 0.0646 | 0.0717 | 0.1138 | 0.0706 | 0.1911 | 0.0553 | 0.2252 |
| | Labels | Avg | 0.7220 | 0.7662 | 0.6814 | 0.8672 | 0.6751 | 0.2851 | 0.7487 | 0.4081 |
| | | STD | 0.1323 | 0.1058 | 0.1008 | 0.1472 | 0.0918 | 0.2709 | 0.0840 | 0.3170 |
| | | *P* value | 0.7435 | 0.9457 | 0.5957 | 0.7491 | 0.5691 | 0.8898 | 0.6040 | 0.9915 |
| **Ubiquitination (K)** | No labels | Avg | 0.6513 | 0.6147 | 0.6134 | 0.5674 | 0.5738 | 0.2185 | 0.5535 | 0.6439 |
| | | STD | 0.0473 | 0.0704 | 0.0412 | 0.1744 | 0.0632 | 0.0899 | 0.1158 | 0.1352 |
| | Labels | Avg | 0.8258 | 0.8356 | 0.7290 | 0.7215 | 0.7130 | 0.4691 | 0.7061 | 0.7383 |
| | | STD | 0.0347 | 0.0288 | 0.0532 | 0.0980 | 0.0906 | 0.0855 | 0.0390 | 0.1419 |
| | | *P* value | 1.00E−07 | 1.63E−06 | 8.13E−05 | 3.65E−02 | 1.63E−03 | 1.01E−05 | 3.21E−03 | 1.66E−01 |

| | | | | | | | | | | |
|---|---|---|---|---|---|---|---|---|---|---|
| **SUMOylation (K)** | No labels | Avg | 0.8522 | 0.8619 | 0.7843 | 0.7664 | 0.7942 | 0.5763 | 0.7748 | 0.8057 |
| | | STD | 0.0510 | 0.0557 | 0.0271 | 0.0829 | 0.0623 | 0.0519 | 0.0344 | 0.0768 |
| | Labels | Avg | 0.8688 | 0.8756 | 0.7863 | 0.7774 | 0.7873 | 0.5773 | 0.7775 | 0.7944 |
| | | STD | 0.0616 | 0.0626 | 0.0484 | 0.1048 | 0.0606 | 0.0969 | 0.0595 | 0.0708 |
| | | *P* value | 0.5418 | 0.6303 | 0.9121 | 0.8071 | 0.8141 | 0.9801 | 0.9058 | 0.7487 |
| **Acetylation (K)** | No labels | Avg | 0.7320 | 0.7069 | 0.6717 | 0.6286 | 0.6802 | 0.3462 | 0.6512 | 0.7151 |
| | | STD | 0.0248 | 0.0281 | 0.0200 | 0.0540 | 0.0360 | 0.0406 | 0.0263 | 0.0453 |
| | Labels | Avg | 0.8530 | 0.8675 | 0.7708 | 0.7161 | 0.7967 | 0.5443 | 0.7532 | 0.8239 |
| | | STD | 0.0102 | 0.0086 | 0.0115 | 0.0379 | 0.0214 | 0.0198 | 0.0164 | 0.0258 |
| | | *P* value | 1.33E−08 | 6.48E−09 | 2.85E−09 | 1.06E−03 | 6.08E−07 | 6.60E−09 | 5.73E−08 | 1.91E−05 |
| **Methylation (K)** | No labels | Avg | 0.7600 | 0.7454 | 0.7239 | 0.5804 | 0.7132 | 0.4209 | 0.6225 | 0.8097 |
| | | STD | 0.0661 | 0.0924 | 0.0587 | 0.1514 | 0.1052 | 0.1118 | 0.0968 | 0.1348 |
| | Labels | Avg | 0.7815 | 0.7725 | 0.7383 | 0.5998 | 0.7206 | 0.4504 | 0.6464 | 0.8300 |
| | | STD | 0.0601 | 0.0667 | 0.0649 | 0.1141 | 0.1392 | 0.1466 | 0.1011 | 0.0930 |
| | | *P* value | 0.4805 | 0.4867 | 0.6273 | 0.7635 | 0.9008 | 0.6376 | 0.6133 | 0.7152 |
| **Methylation (R)** | No labels | Avg | 0.9216 | 0.9197 | 0.8599 | 0.8418 | 0.8656 | 0.7211 | 0.8523 | 0.8770 |
| | | STD | 0.0109 | 0.0162 | 0.0194 | 0.0431 | 0.0359 | 0.0384 | 0.0233 | 0.0387 |
| | Labels | Avg | 0.9472 | 0.9528 | 0.8861 | 0.8596 | 0.9015 | 0.7739 | 0.8789 | 0.9114 |
| | | STD | 0.0088 | 0.0098 | 0.0100 | 0.0360 | 0.0297 | 0.0190 | 0.0123 | 0.0298 |
| | | *P* value | 3.83E−05 | 1.06E−04 | 3.06E−03 | 3.55E−01 | 3.36E−02 | 2.60E−03 | 9.08E−03 | 5.00E−02 |
| **Pyroglutamylation (Q)** | No labels | Avg | 0.9205 | 0.9352 | 0.8312 | 0.7940 | 0.8438 | 0.6704 | 0.8016 | 0.8782 |
| | | STD | 0.0848 | 0.0771 | 0.0568 | 0.1415 | 0.1609 | 0.1284 | 0.0954 | 0.1073 |
| | Labels | Avg | 0.9173 | 0.9262 | 0.8363 | 0.7995 | 0.8381 | 0.6695 | 0.8078 | 0.8718 |
| | | STD | 0.0875 | 0.0795 | 0.0701 | 0.1232 | 0.1609 | 0.1511 | 0.1102 | 0.1233 |
| | | *P* value | 0.9388 | 0.8105 | 0.8670 | 0.9302 | 0.9411 | 0.9899 | 0.9001 | 0.9080 |
| **Palmitoylation (C)** | No labels | Avg | 0.8388 | 0.8346 | 0.7213 | 0.5779 | 0.8133 | 0.4631 | 0.6668 | 0.8633 |
| | | STD | 0.0580 | 0.0685 | 0.0685 | 0.1410 | 0.0649 | 0.1251 | 0.1073 | 0.0547 |
| | Labels | Avg | 0.9353 | 0.9436 | 0.8527 | 0.8380 | 0.8681 | 0.7068 | 0.8514 | 0.8688 |
| | | STD | 0.0260 | 0.0248 | 0.0467 | 0.0575 | 0.0568 | 0.0932 | 0.0466 | 0.0599 |
| | | *P* value | 0.0006 | 0.0009 | 0.0002 | 0.0003 | 0.0728 | 0.0002 | 0.0005 | 0.8411 |

| | | | | | | | | | | |
|---|---|---|---|---|---|---|---|---|---|---|
| **Hydroxylation (P)** | No labels | Avg | 0.9813 | 0.9807 | 0.9635 | 0.9429 | 0.9851 | 0.9274 | 0.9633 | 0.9847 |
| | | STD | 0.0184 | 0.0273 | 0.0229 | 0.0364 | 0.0175 | 0.0457 | 0.0249 | 0.0165 |
| | Labels | Avg | 0.9893 | 0.9928 | 0.9750 | 0.9606 | 0.9898 | 0.9499 | 0.9748 | 0.9889 |
| | | STD | 0.0118 | 0.0081 | 0.0141 | 0.0272 | 0.0103 | 0.0282 | 0.0161 | 0.0112 |
| | | *P* value | 0.2905 | 0.2299 | 0.2214 | 0.2594 | 0.4938 | 0.2266 | 0.2621 | 0.5454 |
| **Hydroxylation (K)** | No labels | Avg | 0.9710 | 0.9815 | 0.9765 | 0.9600 | 0.9933 | 0.9423 | 0.9749 | 0.9667 |
| | | STD | 0.0547 | 0.0418 | 0.0288 | 0.0663 | 0.0200 | 0.0751 | 0.0358 | 0.1000 |
| | Labels | Avg | 0.9752 | 0.9785 | 0.9647 | 0.9400 | 0.9790 | 0.9165 | 0.9561 | 0.9576 |
| | | STD | 0.0399 | 0.0366 | 0.0288 | 0.0800 | 0.0452 | 0.0732 | 0.0425 | 0.1007 |
| | | *P* value | 0.8517 | 0.8760 | 0.3979 | 0.5711 | 0.4027 | 0.4702 | 0.3237 | 0.8497 |

#### 5.2. Model results in All-Organism Datasets from MusiteDeep

| PTM | | | AUC | AUPRC | Accuracy | Recall | Precision | MCC | F1 | Specificity |
|---|---|---|---|---|---|---|---|---|---|---|
| **Phosphorylation (S,T)** | No labels | Avg | 0.8690 | 0.8527 | 0.7928 | 0.8017 | 0.7882 | 0.5862 | 0.7945 | 0.7838 |
| | | STD | 0.0129 | 0.0117 | 0.0120 | 0.0301 | 0.0115 | 0.0239 | 0.0143 | 0.0202 |
| | Labels | Avg | 0.9275 | 0.9217 | 0.8564 | 0.8714 | 0.8463 | 0.7132 | 0.8586 | 0.8414 |
| | | STD | 0.0038 | 0.0042 | 0.0054 | 0.0072 | 0.0098 | 0.0106 | 0.0049 | 0.0126 |
| | | *P* value | 7.25E−08 | 2.78E−09 | 3.27E−09 | 5.06E−05 | 1.27E−09 | 3.62E−09 | 5.74E−08 | 2.63E−06 |
| **Phosphorylation (Y)** | No labels | Avg | 0.8921 | 0.8556 | 0.8342 | 0.8027 | 0.7970 | 0.6370 | 0.7993 | 0.8270 |
| | | STD | 0.0319 | 0.0302 | 0.0164 | 0.0595 | 0.0282 | 0.0588 | 0.0409 | 0.1123 |
| | Labels | Avg | 0.9008 | 0.8693 | 0.8356 | 0.8336 | 0.7857 | 0.6497 | 0.8080 | 0.8211 |
| | | STD | 0.0263 | 0.0320 | 0.0138 | 0.0297 | 0.0480 | 0.0453 | 0.0305 | 0.0706 |
| | | *P* value | 0.5396 | 0.3609 | 0.8549 | 0.1863 | 0.5528 | 0.6147 | 0.6141 | 0.8955 |
| **N-Glycosylation (N)** | No labels | Avg | 0.9823 | 0.9691 | 0.9753 | 0.9967 | 0.9538 | 0.9478 | 0.9747 | 0.9509 |
| | | STD | 0.0013 | 0.0109 | 0.0076 | 0.0013 | 0.0151 | 0.0040 | 0.0082 | 0.0027 |
| | Labels | Avg | 0.9887 | 0.9842 | 0.9767 | 0.9957 | 0.9574 | 0.9494 | 0.9761 | 0.9539 |
| | | STD | 0.0019 | 0.0056 | 0.0070 | 0.0022 | 0.0141 | 0.0026 | 0.0076 | 0.0061 |
| | | *P* value | 4.10E−07 | 2.50E−03 | 6.86E−01 | 2.55E−01 | 6.03E−01 | 3.08E−01 | 7.09E−01 | 1.99E−01 |
| **O-Glycosylation (S,T)** | No labels | Avg | 0.9472 | 0.9332 | 0.8691 | 0.8698 | 0.8188 | 0.7326 | 0.8412 | 0.8704 |
| | | STD | 0.0183 | 0.0284 | 0.0230 | 0.0366 | 0.0746 | 0.0519 | 0.0409 | 0.0453 |
| | Labels | Avg | 0.9598 | 0.9435 | 0.8921 | 0.8930 | 0.8545 | 0.7811 | 0.8708 | 0.8929 |
| | | STD | 0.0191 | 0.0248 | 0.0412 | 0.0598 | 0.0646 | 0.0778 | 0.0432 | 0.0628 |
| | | *P* value | 0.1722 | 0.4237 | 0.1660 | 0.3370 | 0.2917 | 0.1395 | 0.1523 | 0.3973 |
| **Ubiquitination (K)** | No labels | Avg | 0.8876 | 0.8014 | 0.8256 | 0.7980 | 0.7138 | 0.6210 | 0.7519 | 0.8370 |
| | | STD | 0.0302 | 0.0545 | 0.0304 | 0.0478 | 0.0598 | 0.0609 | 0.0416 | 0.0494 |
| | Labels | Avg | 0.9228 | 0.8623 | 0.8740 | 0.8397 | 0.7967 | 0.7216 | 0.8165 | 0.8910 |
| | | STD | 0.0215 | 0.0291 | 0.0289 | 0.0424 | 0.0426 | 0.0546 | 0.0299 | 0.0340 |
| | | *P* value | 0.0114 | 0.0106 | 0.0028 | 0.0664 | 0.0037 | 0.0017 | 0.0016 | 0.0159 |
| **SUMOylation (K)** | No labels | Avg | 0.9566 | 0.9302 | 0.9067 | 0.8912 | 0.8687 | 0.8033 | 0.8773 | 0.9194 |
| | | STD | 0.0165 | 0.0321 | 0.0262 | 0.0392 | 0.0736 | 0.0557 | 0.0379 | 0.0387 |

| | | | | | | | | | | |
|---|---|---|---|---|---|---|---|---|---|---|
| | Labels | Avg | 0.9661 | 0.9448 | 0.9135 | 0.9096 | 0.8699 | 0.8185 | 0.8867 | 0.9191 |
| | | STD | 0.0167 | 0.0239 | 0.0277 | 0.0480 | 0.0725 | 0.0598 | 0.0408 | 0.0373 |
| | | *P* value | 0.2418 | 0.2872 | 0.5972 | 0.3849 | 0.9717 | 0.5837 | 0.6182 | 0.9865 |
| **Acetylation (K)** | No labels | Avg | 0.8631 | 0.7605 | 0.7937 | 0.7567 | 0.7023 | 0.5637 | 0.7281 | 0.8147 |
| | | STD | 0.0125 | 0.0204 | 0.0144 | 0.0180 | 0.0243 | 0.0243 | 0.0123 | 0.0270 |
| | Labels | Avg | 0.9044 | 0.8361 | 0.8357 | 0.8215 | 0.7541 | 0.6559 | 0.7850 | 0.8441 |
| | | STD | 0.0185 | 0.0239 | 0.0192 | 0.0342 | 0.0389 | 0.0331 | 0.0195 | 0.0380 |
| | | *P* value | 4.73E−05 | 1.18E−06 | 6.96E−05 | 2.01E−04 | 4.05E−03 | 4.03E−06 | 2.06E−06 | 7.65E−02 |
| **Methylation (K)** | No labels | Avg | 0.9648 | 0.9046 | 0.9213 | 0.8915 | 0.8252 | 0.8039 | 0.8542 | 0.9338 |
| | | STD | 0.0161 | 0.0475 | 0.0228 | 0.0514 | 0.0755 | 0.0565 | 0.0420 | 0.0287 |
| | Labels | Avg | 0.9677 | 0.9167 | 0.9224 | 0.8851 | 0.8293 | 0.8038 | 0.8538 | 0.9367 |
| | | STD | 0.0104 | 0.0367 | 0.0222 | 0.0494 | 0.0753 | 0.0586 | 0.0466 | 0.0267 |
| | | *P* value | 0.6586 | 0.5539 | 0.9228 | 0.7921 | 0.9078 | 0.9993 | 0.9862 | 0.8318 |
| **Methylation (R)** | No labels | Avg | 0.9282 | 0.8695 | 0.8643 | 0.8672 | 0.7930 | 0.7191 | 0.8277 | 0.8625 |
| | | STD | 0.0171 | 0.0247 | 0.0195 | 0.0389 | 0.0317 | 0.0404 | 0.0250 | 0.0266 |
| | Labels | Avg | 0.9451 | 0.9062 | 0.8842 | 0.8802 | 0.8245 | 0.7579 | 0.8512 | 0.8865 |
| | | STD | 0.0134 | 0.0161 | 0.0147 | 0.0237 | 0.0219 | 0.0298 | 0.0180 | 0.0186 |
| | | *P* value | 0.0321 | 0.0019 | 0.0254 | 0.4076 | 0.0263 | 0.0334 | 0.0360 | 0.0413 |
| **Pyroglutamylation (Q)** | No labels | Avg | 0.9610 | 0.9414 | 0.9072 | 0.8850 | 0.9049 | 0.8128 | 0.8927 | 0.9239 |
| | | STD | 0.0132 | 0.0307 | 0.0306 | 0.0812 | 0.0242 | 0.0630 | 0.0468 | 0.0189 |
| | Labels | Avg | 0.9797 | 0.9781 | 0.9408 | 0.9346 | 0.9338 | 0.8808 | 0.9337 | 0.9472 |
| | | STD | 0.0080 | 0.0085 | 0.0126 | 0.0309 | 0.0249 | 0.0259 | 0.0167 | 0.0142 |
| | | *P* value | 0.0025 | 0.0059 | 0.0100 | 0.1135 | 0.0223 | 0.0112 | 0.0304 | 0.0093 |
| **Palmitoylation (C)** | No labels | Avg | 0.9492 | 0.9236 | 0.8832 | 0.8061 | 0.8963 | 0.7570 | 0.8471 | 0.9375 |
| | | STD | 0.0168 | 0.0216 | 0.0328 | 0.0629 | 0.0282 | 0.0600 | 0.0358 | 0.0197 |
| | Labels | Avg | 0.9721 | 0.9493 | 0.9339 | 0.9314 | 0.9064 | 0.8629 | 0.9180 | 0.9369 |
| | | STD | 0.0085 | 0.0218 | 0.0158 | 0.0304 | 0.0342 | 0.0322 | 0.0189 | 0.0210 |
| | | *P* value | 0.0029 | 0.0216 | 0.0011 | 0.0001 | 0.5031 | 0.0004 | 0.0001 | 0.9491 |
| **Hydroxylation (P)** | No labels | Avg | 0.9088 | 0.8562 | 0.8319 | 0.7685 | 0.8086 | 0.6497 | 0.7874 | 0.8750 |
| | | STD | 0.0125 | 0.0267 | 0.0129 | 0.0350 | 0.0247 | 0.0282 | 0.0198 | 0.0207 |

| | | | | | | | | | | |
|---|---|---|---|---|---|---|---|---|---|---|
| | Labels | Avg | 0.9796 | 0.9676 | 0.9426 | 0.9404 | 0.9196 | 0.8815 | 0.9298 | 0.9442 |
| | | STD | 0.0048 | 0.0088 | 0.0123 | 0.0217 | 0.0162 | 0.0263 | 0.0166 | 0.0091 |
| | | *P* value | 3.15E−09 | 1.35E−07 | 3.44E−13 | 2.31E−09 | 7.19E−09 | 6.14E−13 | 4.20E−12 | 7.00E−07 |
| **Hydroxylation (K)** | No labels | Avg | 0.9186 | 0.8368 | 0.8470 | 0.7803 | 0.7842 | 0.6602 | 0.7734 | 0.8915 |
| | | STD | 0.0330 | 0.0517 | 0.0465 | 0.1016 | 0.0871 | 0.0761 | 0.0473 | 0.0401 |
| | Labels | Avg | 0.9472 | 0.8763 | 0.8687 | 0.8777 | 0.7683 | 0.7154 | 0.8016 | 0.8732 |
| | | STD | 0.0211 | 0.0985 | 0.0440 | 0.0606 | 0.1731 | 0.1101 | 0.1283 | 0.0562 |
| | | *P* value | 0.0450 | 0.3054 | 0.3222 | 0.0262 | 0.8097 | 0.2336 | 0.5476 | 0.4374 |

### 6. Additional ROC Curves

#### 6.1. AUC for Models Trained on MusiteDeep Datasets

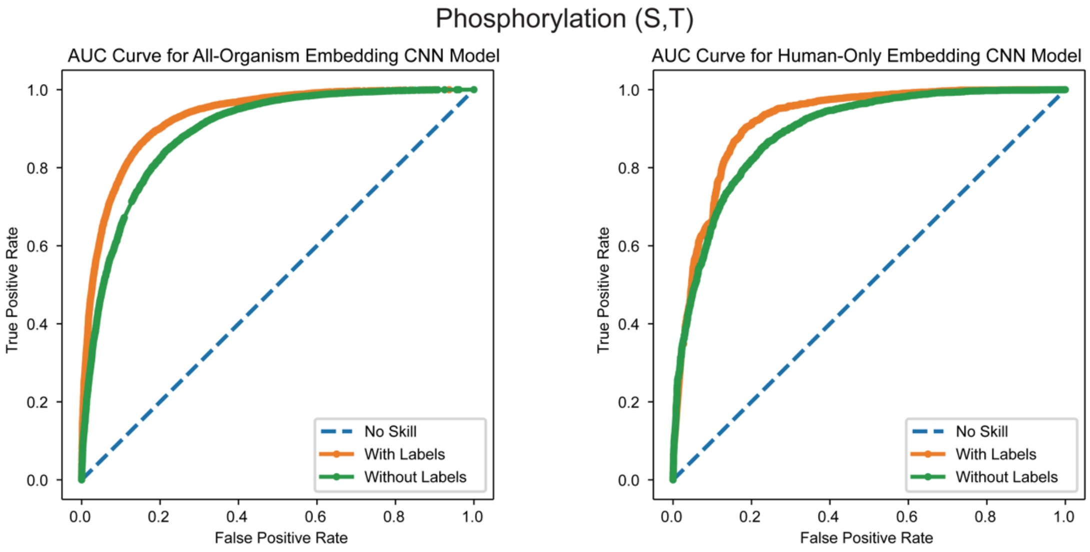


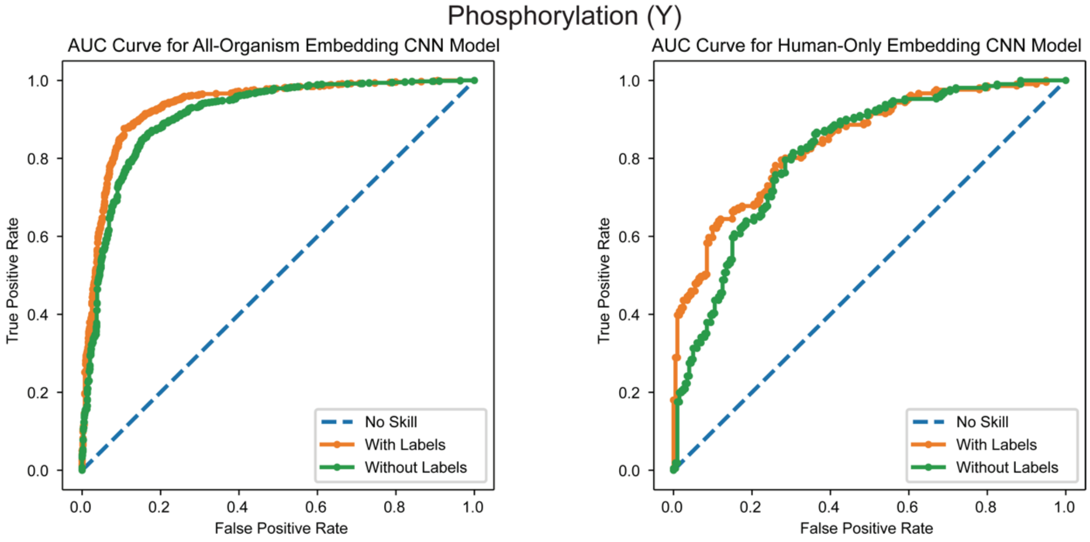

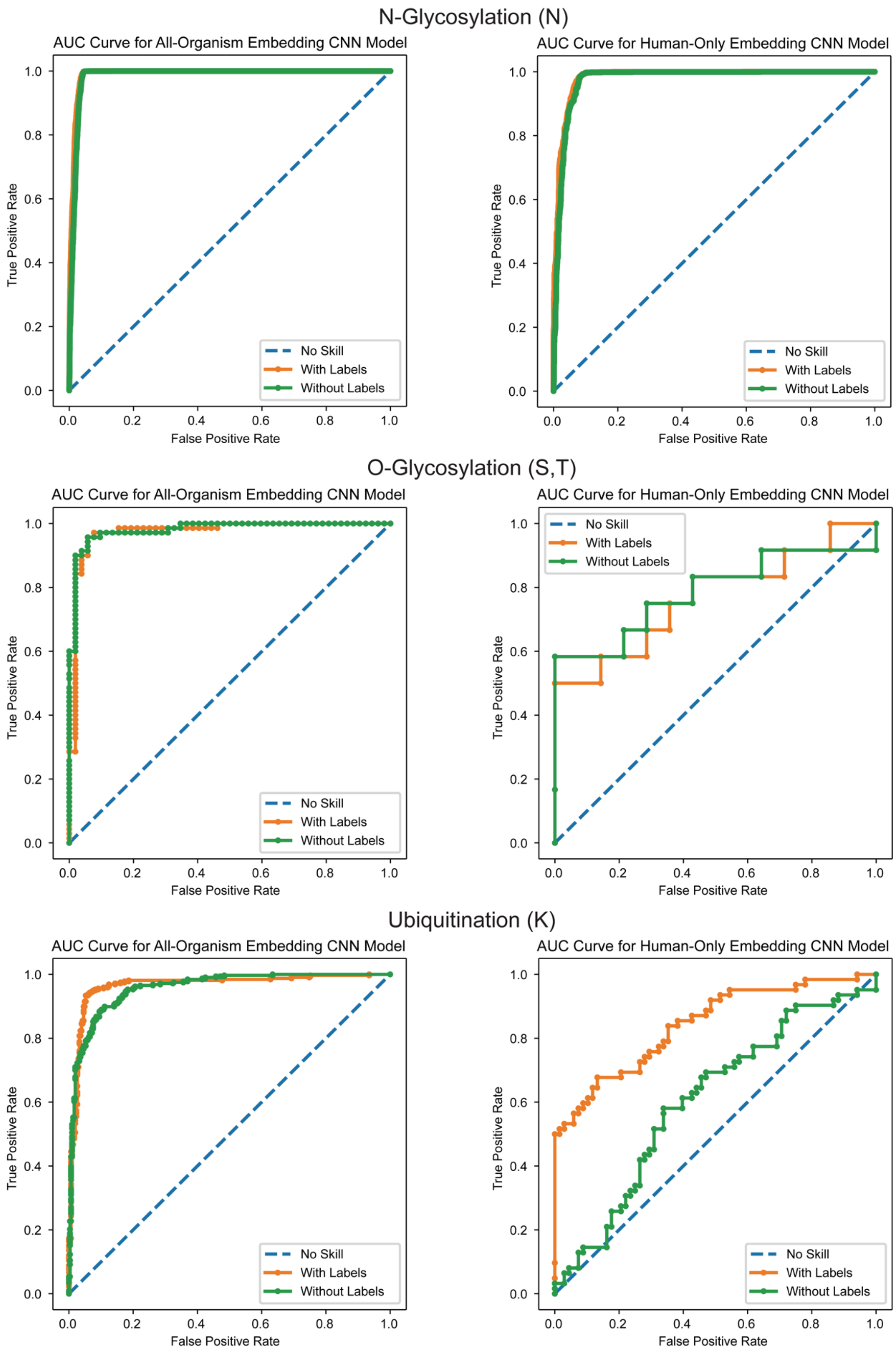
N-Glycosylation (N)
AUC Curve for All-Organism Embedding CNN Model
AUC Curve for Human-Only Embedding CNN Model
O-Glycosylation (S,T)
AUC Curve for All-Organism Embedding CNN Model
AUC Curve for Human-Only Embedding CNN Model
Ubiquitination (K)
AUC Curve for All-Organism Embedding CNN Model
AUC Curve for Human-Only Embedding CNN Model
True Positive Rate
False Positive Rate
No Skill
With Labels
Without Labels
0.0
0.2
0.4
0.6
0.8
1.0

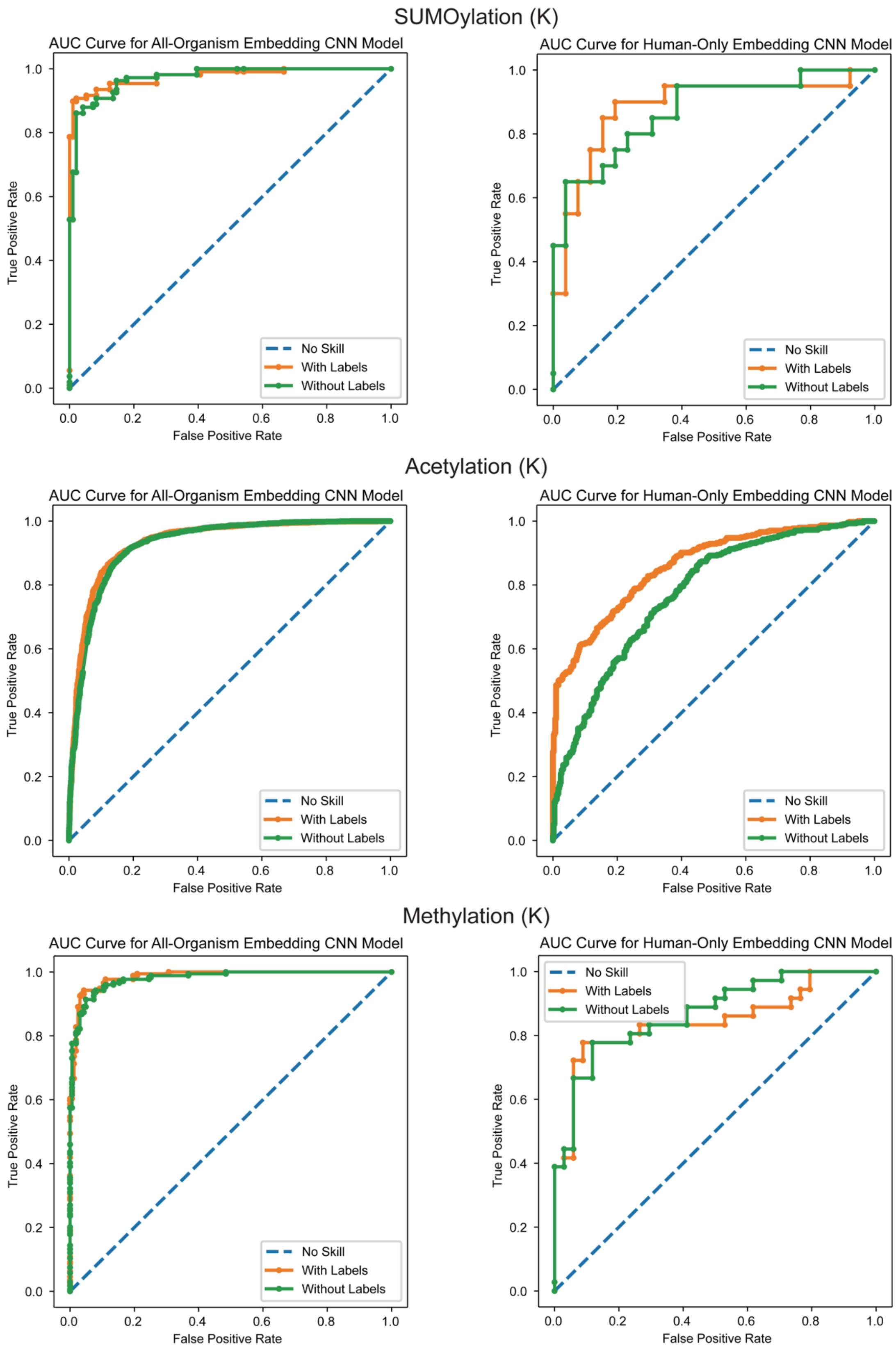
SUMOylation (K)
AUC Curve for All-Organism Embedding CNN Model
AUC Curve for Human-Only Embedding CNN Model
Acetylation (K)
AUC Curve for All-Organism Embedding CNN Model
AUC Curve for Human-Only Embedding CNN Model
Methylation (K)
AUC Curve for All-Organism Embedding CNN Model
AUC Curve for Human-Only Embedding CNN Model
True Positive Rate
False Positive Rate
No Skill
With Labels
Without Labels
0.0
0.2
0.4
0.6
0.8
1.0

#### Methylation (R)

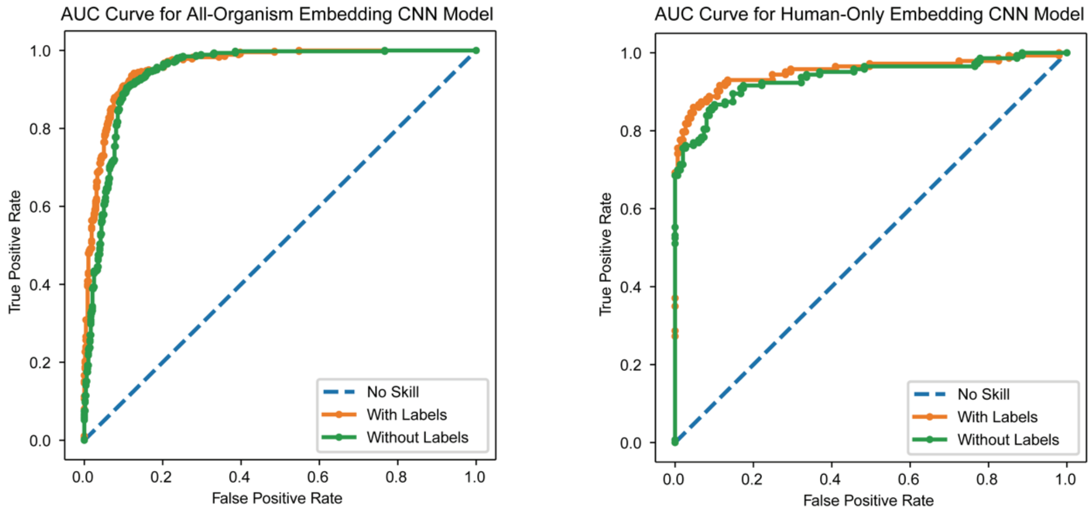


#### Pyroglutamylation (Q)

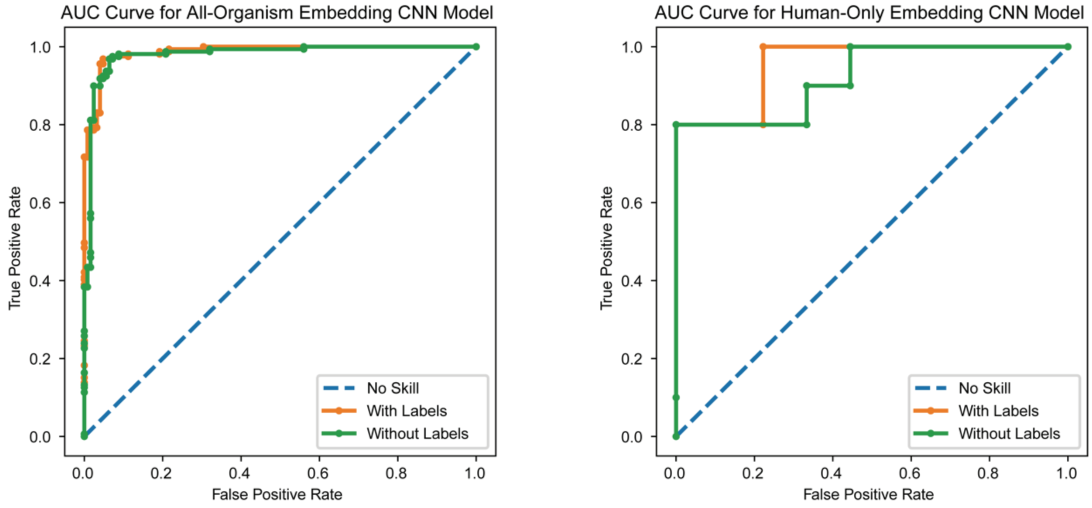


#### Palmitoylation (C)

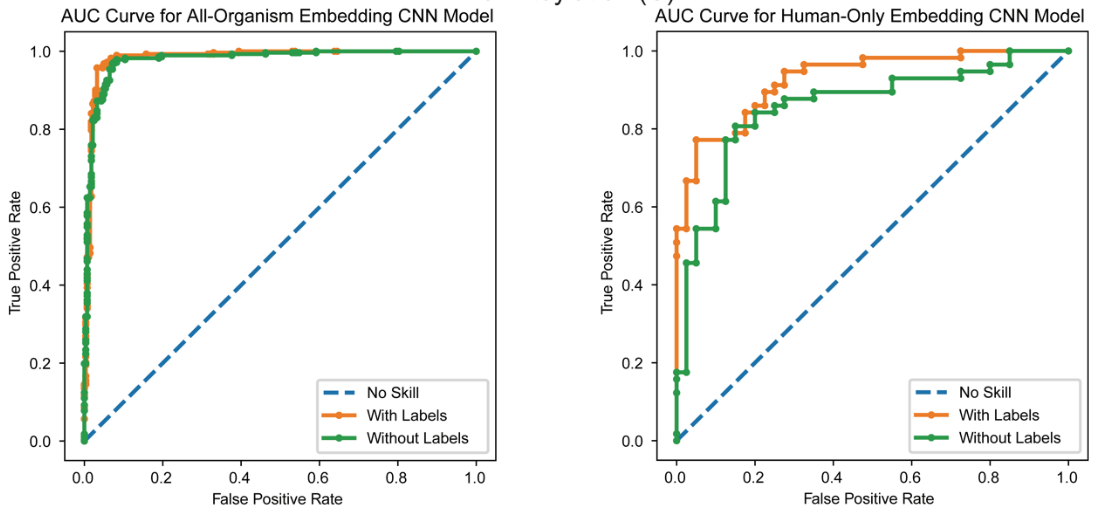

#### Hydroxylation (P)

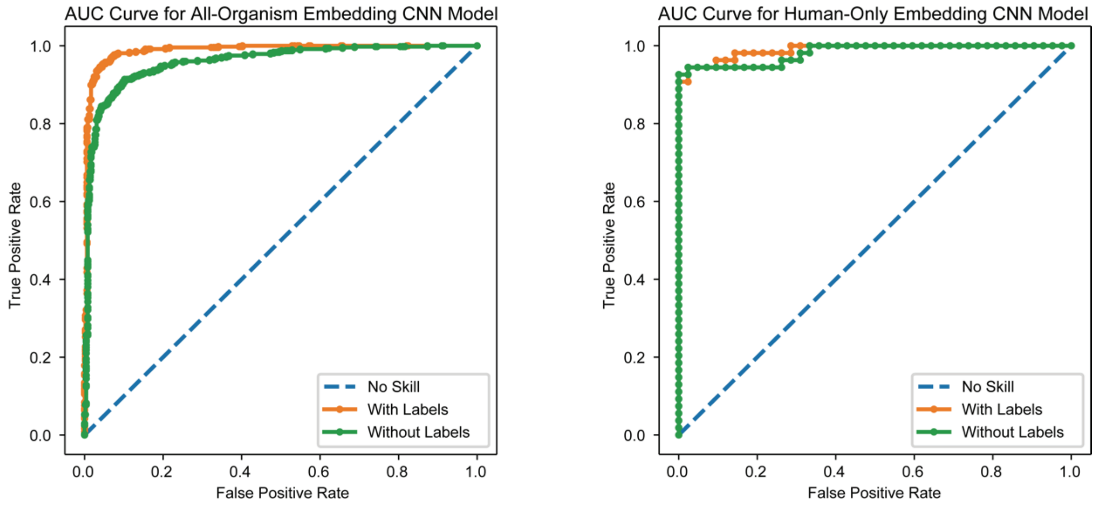


#### Hydroxylation (K)

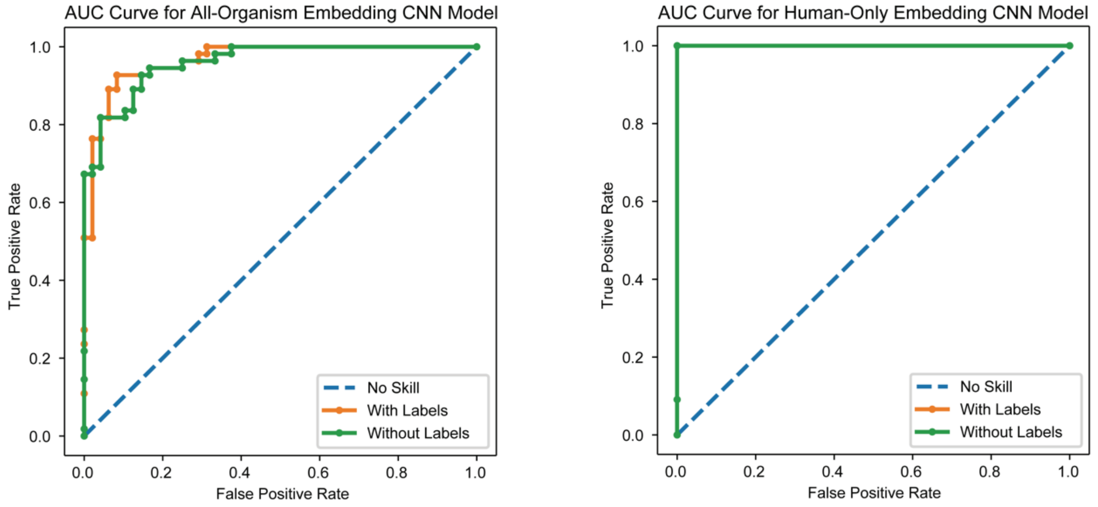

#### 6.2. AUPRC for Models Trained on MusiteDeep Datasets

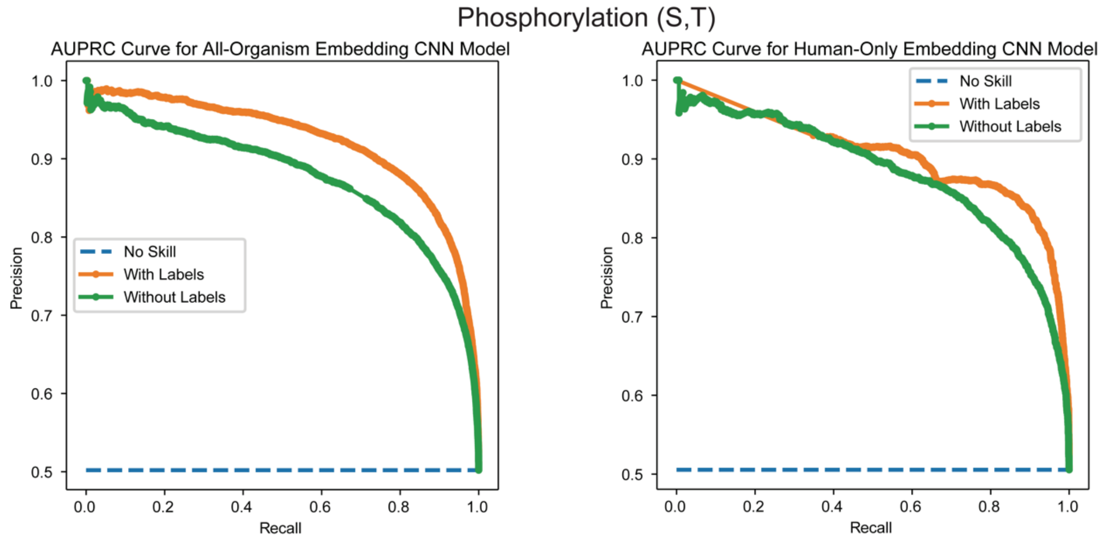


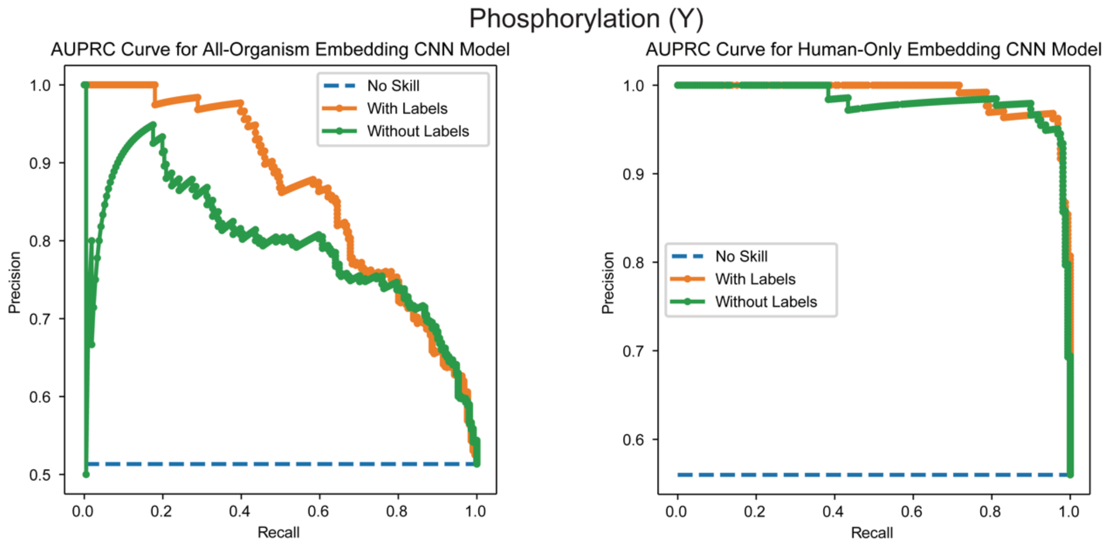

#### N-Glycosylation (N)

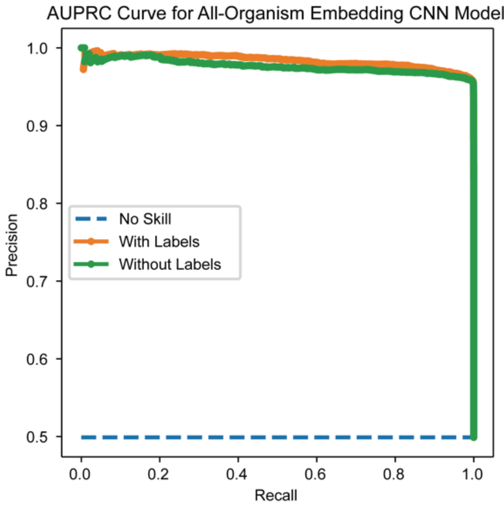


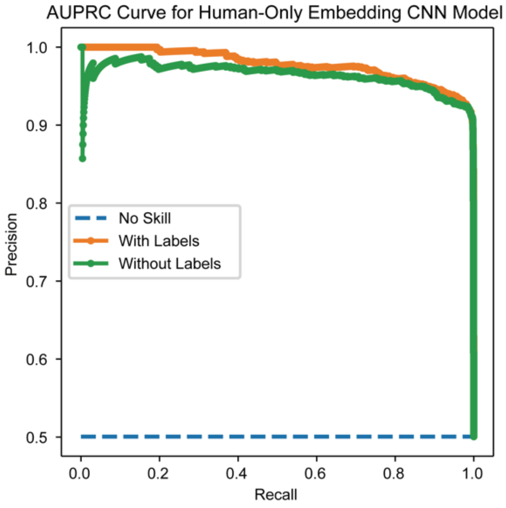


#### O-Glycosylation (S,T)

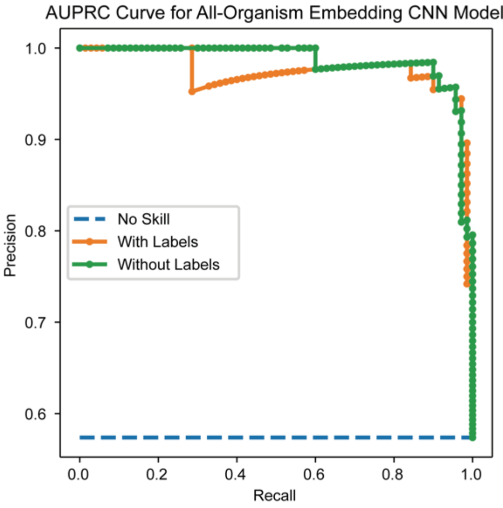


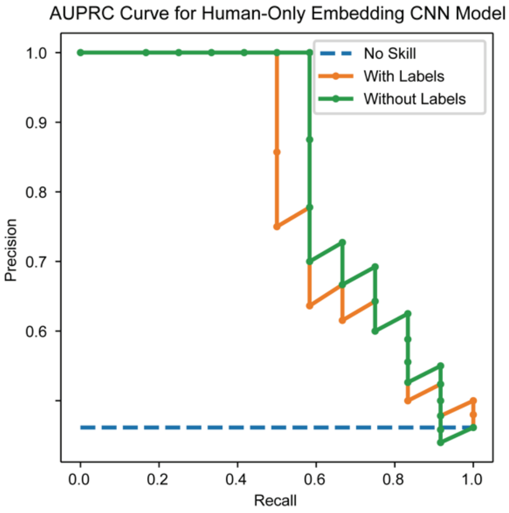


#### Ubiquitination (K)

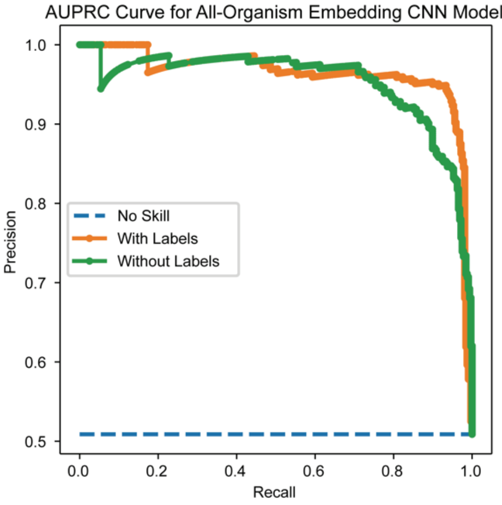


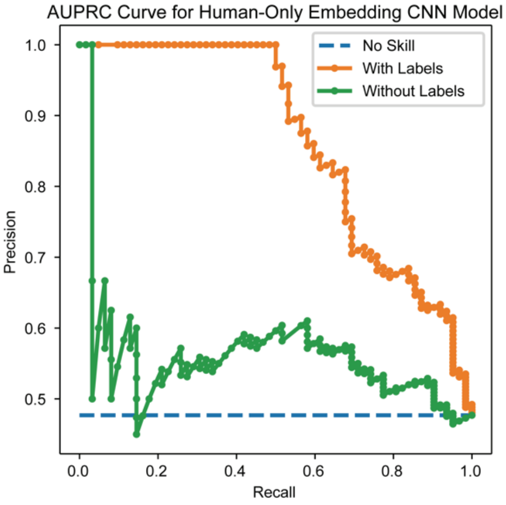

SUMOylation (K)

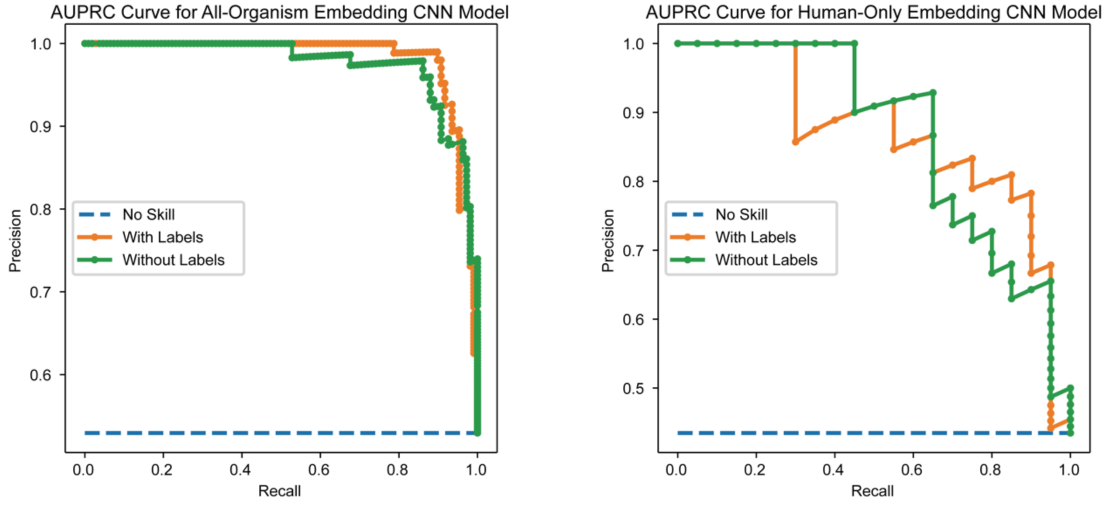


Acetylation (K)

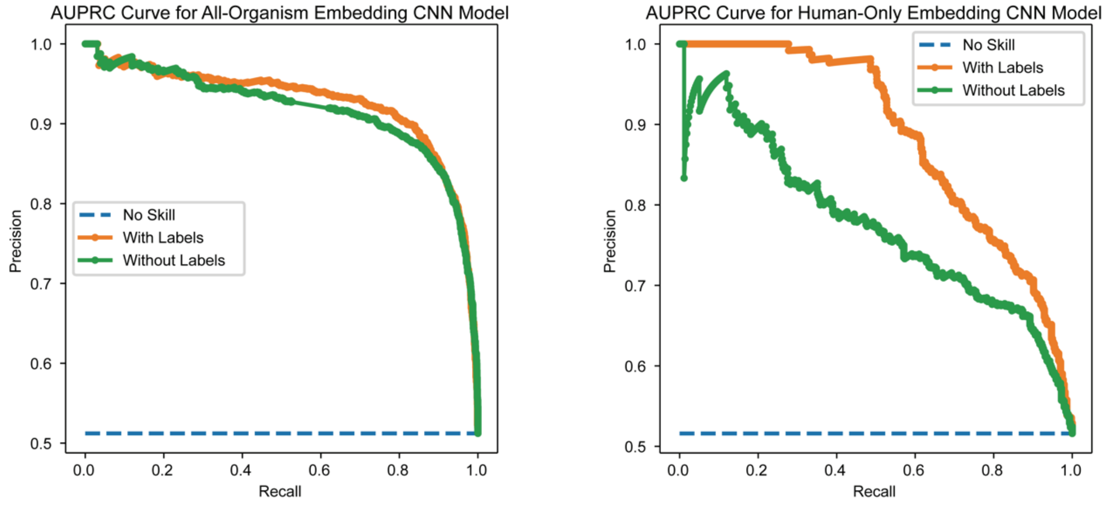


Methylation (K)

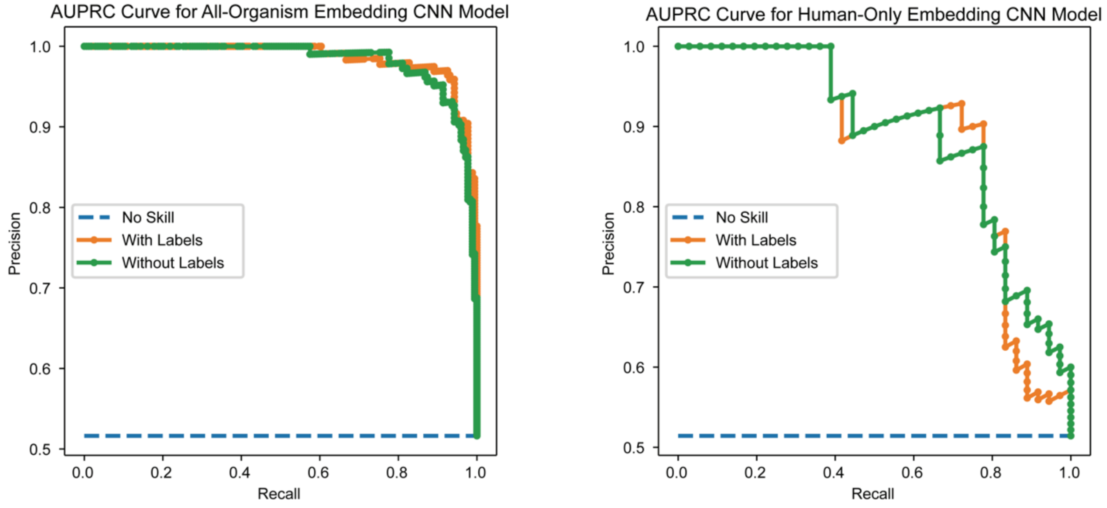

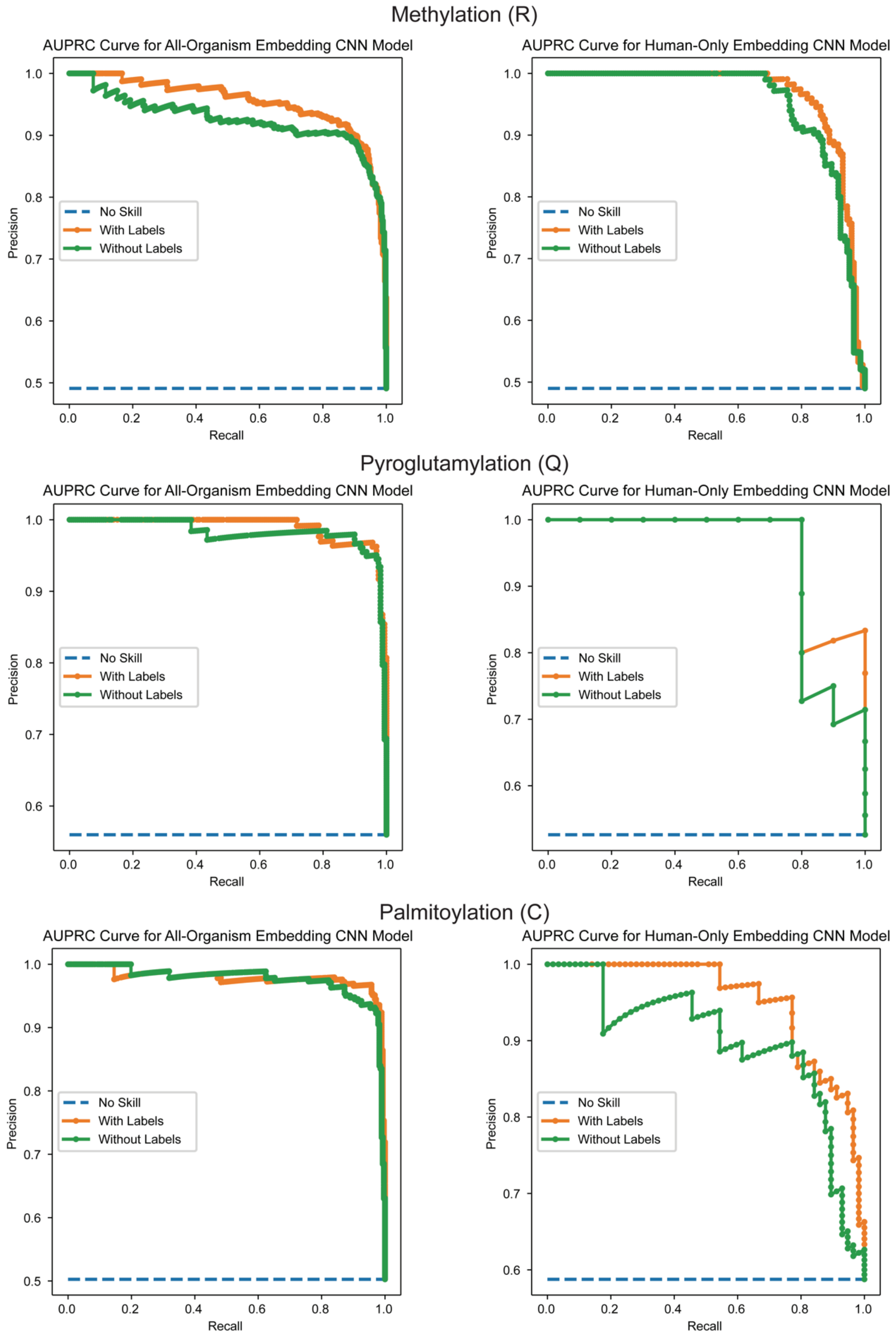
Methylation (R)
AUPRC Curve for All-Organism Embedding CNN Model
AUPRC Curve for Human-Only Embedding CNN Model
Pyroglutamylation (Q)
AUPRC Curve for All-Organism Embedding CNN Model
AUPRC Curve for Human-Only Embedding CNN Model
Palmitoylation (C)
AUPRC Curve for All-Organism Embedding CNN Model
AUPRC Curve for Human-Only Embedding CNN Model
Precision
Recall
No Skill
With Labels
Without Labels

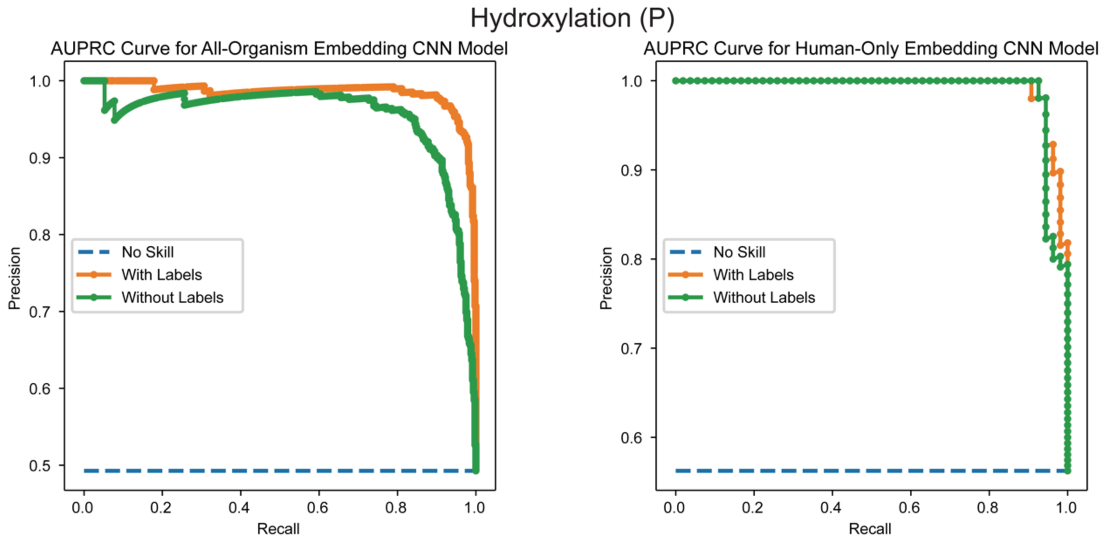
Hydroxylation (P)
AUPRC Curve for All-Organism Embedding CNN Model
AUPRC Curve for Human-Only Embedding CNN Model
Precision
Recall
No Skill
With Labels
Without Labels
0.0
0.2
0.4
0.6
0.8
1.0
0.5
0.6
0.7
0.8
0.9

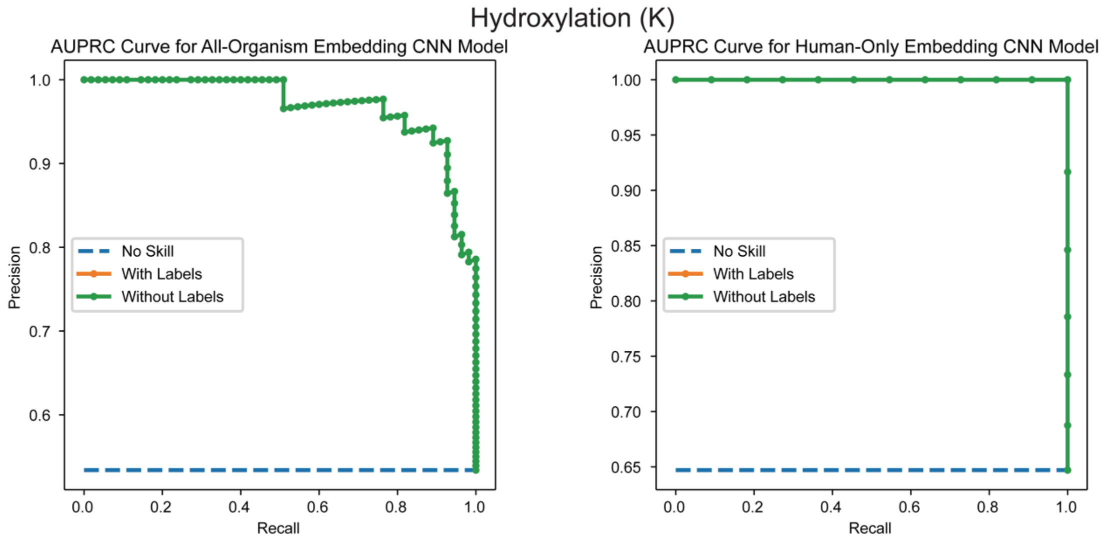
Hydroxylation (K)
AUPRC Curve for All-Organism Embedding CNN Model
AUPRC Curve for Human-Only Embedding CNN Model
Precision
Recall
No Skill
With Labels
Without Labels
0.0
0.2
0.4
0.6
0.8
1.0
0.7
0.9
0.65
0.70
0.75
0.80
0.85
0.90
0.95
1.00

### 7. Additional O-Glycosylation Models

#### 7.1 Model Results

| PTM | | | AUC | AUPRC | Accuracy | Recall | Precision | MCC | F1 | Specificity |
|---|---|---|---|---|---|---|---|---|---|---|
| **OGP Full Dataset, All-Organism** | No labels | Avg | 0.8647 | 0.8670 | 0.7772 | 0.8023 | 0.7624 | 0.5558 | 0.7814 | 0.7525 |
| | | STD | 0.0175 | 0.0190 | 0.0167 | 0.0274 | 0.0194 | 0.0334 | 0.0152 | 0.0276 |
| | Labels | Avg | 0.9574 | 0.9653 | 0.8983 | 0.8637 | 0.9268 | 0.7985 | 0.8939 | 0.9324 |
| | | STD | 0.0065 | 0.0066 | 0.0058 | 0.0162 | 0.0132 | 0.0116 | 0.0073 | 0.0139 |
| | | *P* value | 7.82E−09 | 1.29E−08 | 3.19E−10 | 3.99E−05 | 4.87E−13 | 3.13E−10 | 3.89E−11 | 1.46E−10 |
| **OGP Full Dataset, Human-Only** | No labels | Avg | 0.8544 | 0.8593 | 0.7635 | 0.7751 | 0.7534 | 0.5277 | 0.7635 | 0.7516 |
| | | STD | 0.0163 | 0.0234 | 0.0183 | 0.0423 | 0.0238 | 0.0378 | 0.0261 | 0.0267 |
| | Labels | Avg | 0.9531 | 0.9631 | 0.8970 | 0.8534 | 0.9328 | 0.7968 | 0.8912 | 0.9399 |
| | | STD | 0.0052 | 0.0038 | 0.0079 | 0.0176 | 0.0119 | 0.0147 | 0.0085 | 0.0103 |
| | | *P* value | 3.24E−09 | 2.14E−07 | 1.01E−10 | 2.53E−04 | 2.44E−11 | 2.23E−10 | 2.74E−08 | 2.77E−10 |
| **OGP GalNAc only, All-Organism** | No labels | Avg | 0.8939 | 0.8990 | 0.8060 | 0.8441 | 0.7851 | 0.6141 | 0.8131 | 0.7678 |
| | | STD | 0.0154 | 0.0161 | 0.0155 | 0.0211 | 0.0271 | 0.0295 | 0.0167 | 0.0313 |
| | Labels | Avg | 0.9625 | 0.9682 | 0.8988 | 0.8721 | 0.9218 | 0.7987 | 0.8961 | 0.9251 |
| | | STD | 0.0044 | 0.0037 | 0.0088 | 0.0145 | 0.0134 | 0.0177 | 0.0094 | 0.0158 |
| | | *P* value | 9.51E−08 | 1.95E−07 | 2.17E−10 | 4.79E−03 | 4.07E−09 | 9.39E−11 | 2.99E−09 | 3.97E−09 |
| **OGP GalNAc Only Human-Only** | No labels | Avg | 0.8848 | 0.8781 | 0.7898 | 0.7918 | 0.7889 | 0.5828 | 0.7862 | 0.7865 |
| | | STD | 0.0403 | 0.0618 | 0.0468 | 0.1128 | 0.0344 | 0.0907 | 0.0723 | 0.0464 |
| | Labels | Avg | 0.9721 | 0.9781 | 0.9233 | 0.8849 | 0.9595 | 0.8492 | 0.9206 | 0.9620 |
| | | STD | 0.0035 | 0.0018 | 0.0078 | 0.0109 | 0.0118 | 0.0155 | 0.0071 | 0.0121 |
| | | *P* value | 1.07E−04 | 9.05E−04 | 1.03E−05 | 3.55E−02 | 2.00E−08 | 7.84E−06 | 3.33E−04 | 5.48E−07 |
| **OGP HexNAc Only All-Organism** | No labels | Avg | 0.7961 | 0.7724 | 0.7252 | 0.7654 | 0.7039 | 0.4528 | 0.7327 | 0.6855 |
| | | STD | 0.0193 | 0.0155 | 0.0225 | 0.0406 | 0.0222 | 0.0443 | 0.0246 | 0.0359 |
| | Labels | Avg | 0.9139 | 0.9166 | 0.8319 | 0.8219 | 0.8368 | 0.6652 | 0.8280 | 0.8419 |
| | | STD | 0.0050 | 0.0071 | 0.0096 | 0.0381 | 0.0260 | 0.0175 | 0.0110 | 0.0335 |

| | | | | | | | | | | |
|---|---|---|---|---|---|---|---|---|---|---|
| | | *P* value | 5.38E−09 | 3.38E−12 | 1.59E−08 | 6.99E−03 | 1.08E−09 | 1.82E−08 | 1.31E−07 | 1.90E−08 |
| **OGP HexNAc Only, Human-Only** | No labels | Avg | 0.8463 | 0.8218 | 0.7702 | 0.8104 | 0.7489 | 0.5440 | 0.7774 | 0.7312 |
| | | STD | 0.0165 | 0.0253 | 0.0147 | 0.0446 | 0.0214 | 0.0329 | 0.0180 | 0.0295 |
| | Labels | Avg | 0.9366 | 0.9436 | 0.8619 | 0.8458 | 0.8727 | 0.7238 | 0.8589 | 0.8771 |
| | | STD | 0.0079 | 0.0073 | 0.0108 | 0.0111 | 0.0178 | 0.0217 | 0.0098 | 0.0226 |
| | | *P* value | 1.67E−09 | 4.48E−08 | 4.39E−11 | 4.28E−02 | 1.42E−10 | 4.34E−10 | 1.06E−08 | 1.47E−09 |
| **O-GlcNAc Site Atlas, All-Organism** | No labels | Avg | 0.7744 | 0.7647 | 0.6994 | 0.6704 | 0.7128 | 0.4015 | 0.6888 | 0.7282 |
| | | STD | 0.0135 | 0.0194 | 0.0112 | 0.0570 | 0.0260 | 0.0229 | 0.0235 | 0.0523 |
| | Labels | Avg | 0.8794 | 0.8898 | 0.7965 | 0.7621 | 0.8179 | 0.5946 | 0.7887 | 0.8306 |
| | | STD | 0.0055 | 0.0062 | 0.0073 | 0.0201 | 0.0150 | 0.0147 | 0.0095 | 0.0191 |
| | | *P* value | 6.55E−11 | 1.64E−09 | 4.35E−13 | 7.81E−04 | 3.97E−08 | 7.94E−13 | 6.55E−08 | 1.61E−04 |
| **O-GlcNAc Site Atlas, Human-Only** | No labels | Avg | 0.7670 | 0.7631 | 0.7008 | 0.6154 | 0.7376 | 0.4072 | 0.6693 | 0.7847 |
| | | STD | 0.0311 | 0.0337 | 0.0280 | 0.0620 | 0.0276 | 0.0530 | 0.0421 | 0.0316 |
| | Labels | Avg | 0.9122 | 0.9235 | 0.8281 | 0.7668 | 0.8717 | 0.6612 | 0.8154 | 0.8886 |
| | | STD | 0.0035 | 0.0043 | 0.0061 | 0.0210 | 0.0215 | 0.0128 | 0.0080 | 0.0222 |
| | | *P* value | 1.70E−07 | 1.31E−07 | 1.26E−07 | 2.43E−05 | 1.93E−09 | 6.46E−08 | 1.72E−06 | 4.57E−07 |

#### 7.2 AUC ROC Curves

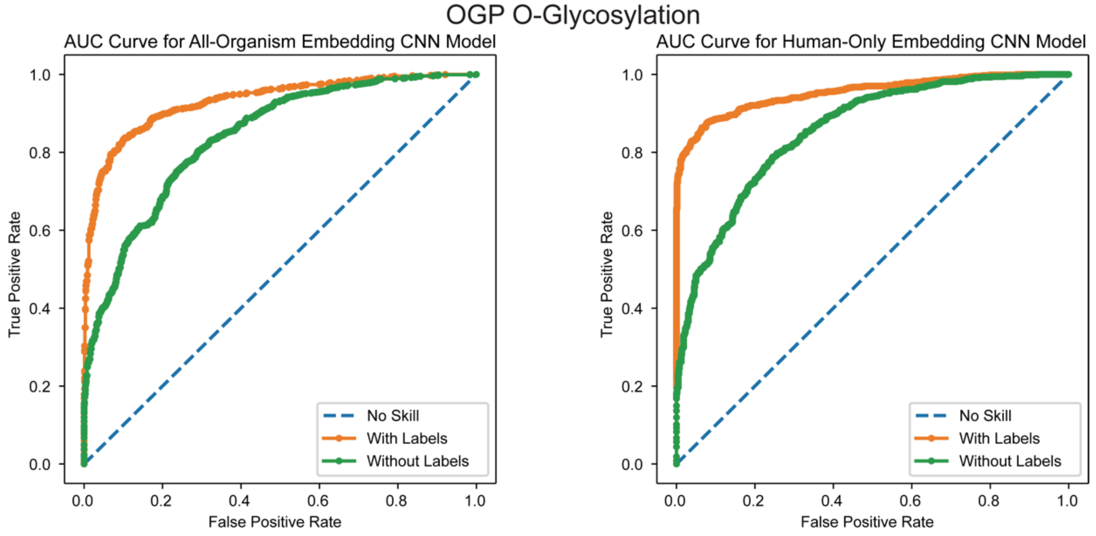


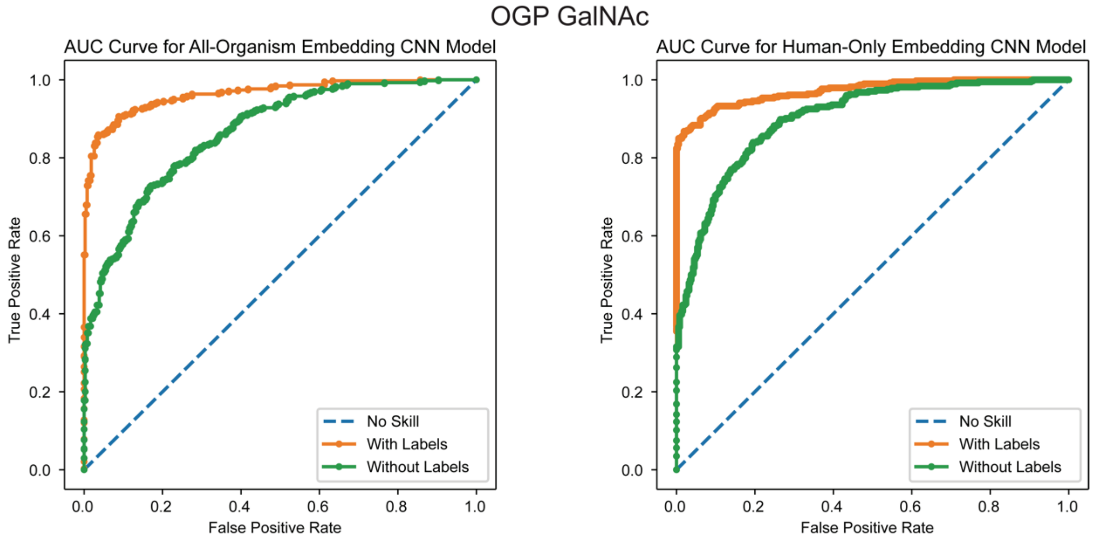

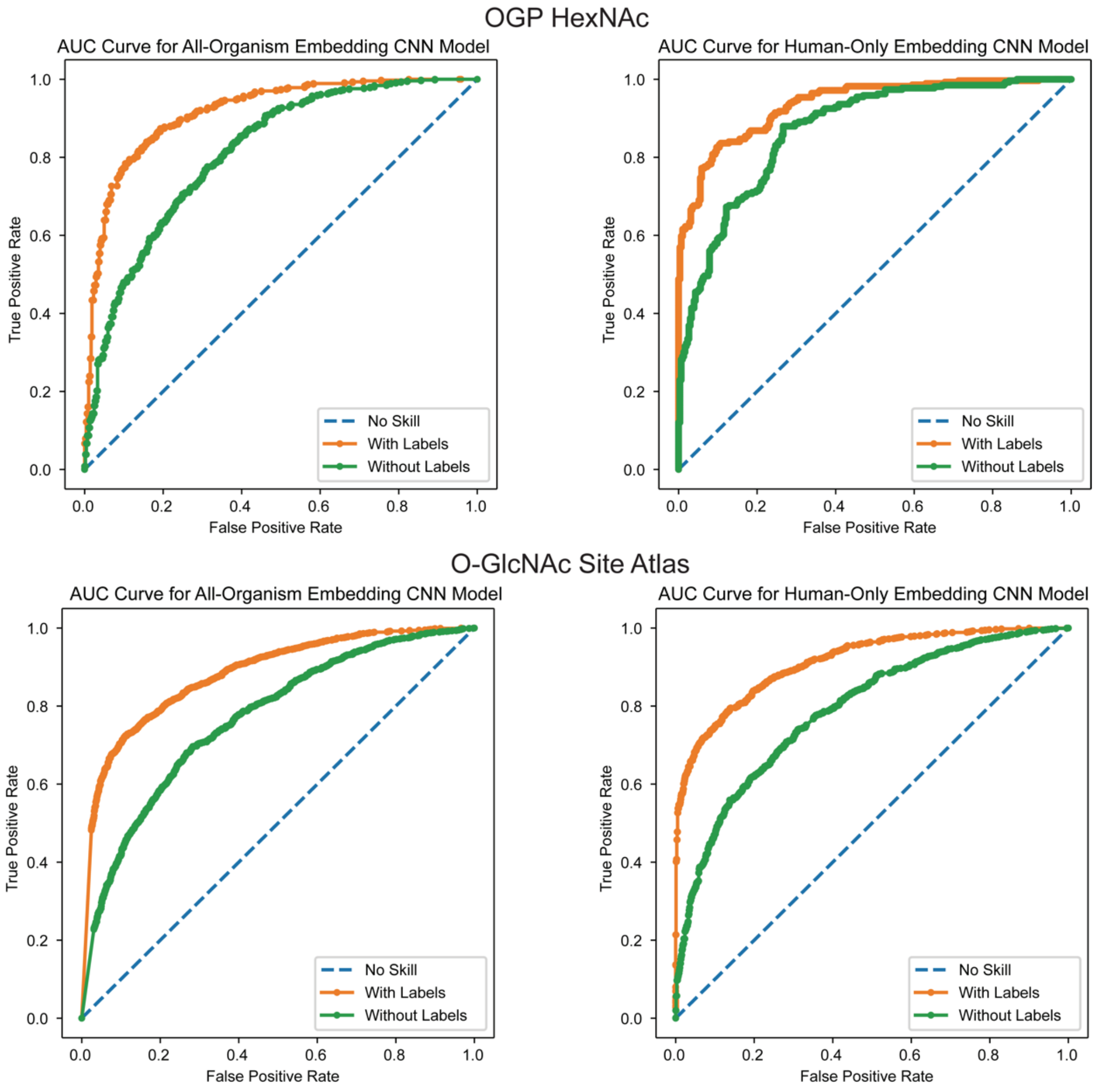
OGP HexNAc
AUC Curve for All-Organism Embedding CNN Model
AUC Curve for Human-Only Embedding CNN Model
O-GlcNAc Site Atlas
AUC Curve for All-Organism Embedding CNN Model
AUC Curve for Human-Only Embedding CNN Model
True Positive Rate
False Positive Rate
No Skill
With Labels
Without Labels

##### 7.3. AUPRC ROC curves

OGP O-Glycosylation

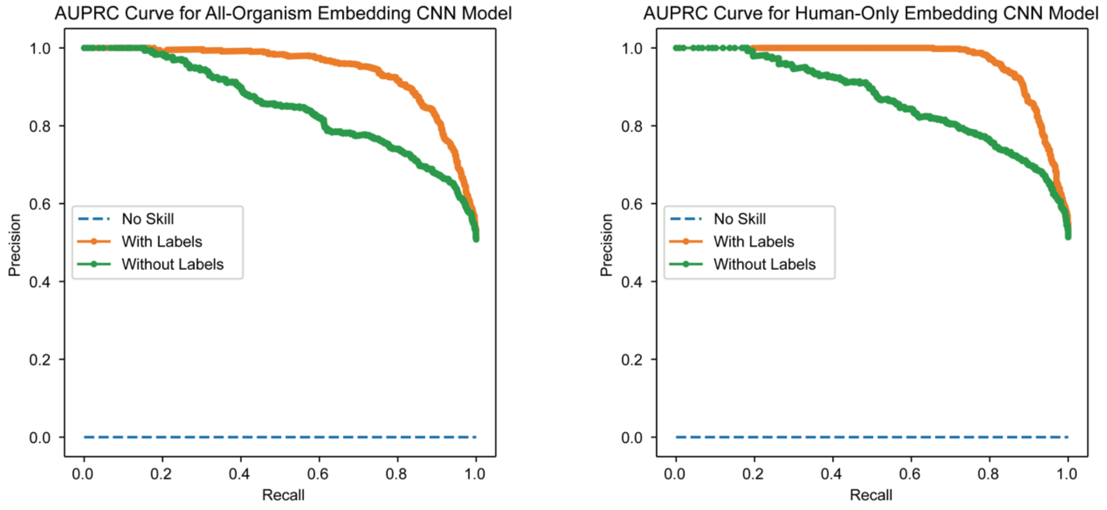


OGP GalNAc

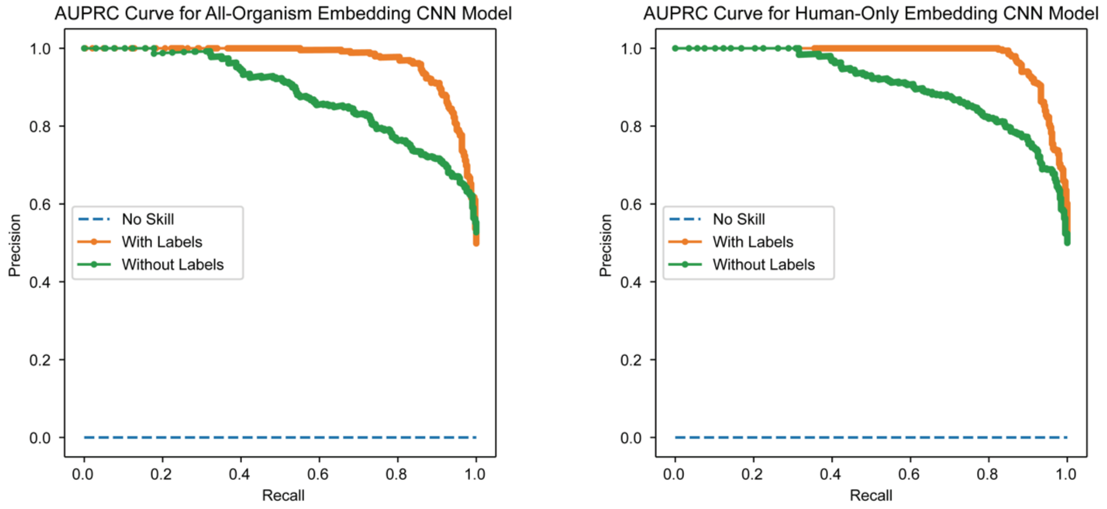

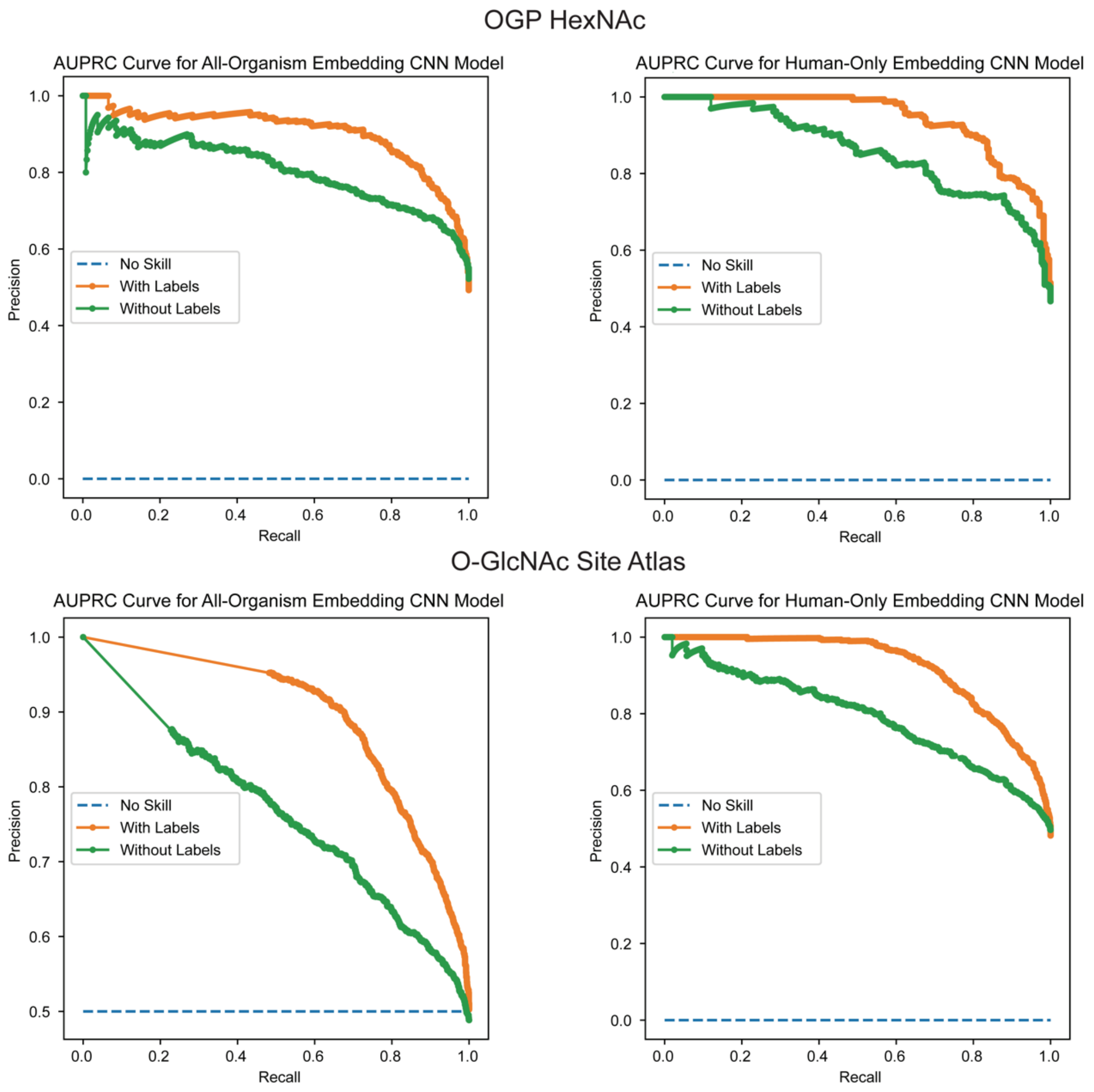
OGP HexNAc
AUPRC Curve for All-Organism Embedding CNN Model
AUPRC Curve for Human-Only Embedding CNN Model
Precision
Recall
No Skill
With Labels
Without Labels
O-GlcNAc Site Atlas
AUPRC Curve for All-Organism Embedding CNN Model
AUPRC Curve for Human-Only Embedding CNN Model
Precision
Recall
No Skill
With Labels
Without Labels

### 8. N-Glycosylation Sequon-Specific Models

#### 8.1. Model Results

| PTM | | | AUC | AUPRC | Accuracy | Recall | Precision | MCC | F1 | Specificity |
|---|---|---|---|---|---|---|---|---|---|---|
| **N-Glycosite Atlas, All-Organism** | No labels | Avg | 0.6557 | 0.4819 | 0.6645 | 0.1751 | 0.5350 | 0.1423 | 0.2577 | 0.6645 |
| | | STD | 0.0114 | 0.0142 | 0.0088 | 0.0634 | 0.0190 | 0.0347 | 0.0752 | 0.0088 |
| | Labels | Avg | 0.6869 | 0.5209 | 0.6792 | 0.2720 | 0.5664 | 0.2086 | 0.3658 | 0.6792 |
| | | STD | 0.0260 | 0.0313 | 0.0136 | 0.0340 | 0.0411 | 0.0320 | 0.0315 | 0.0136 |
| | | *P* value | 0.0013 | 0.0007 | 0.0215 | 0.0000 | 0.0548 | 0.0001 | 0.0001 | 0.0215 |
| **NGlyDE Dataset, Human-Only** | No labels | Avg | 0.6698 | 0.7827 | 0.6913 | 0.9464 | 0.7028 | 0.1838 | 0.8044 | 0.6913 |
| | | STD | 0.0206 | 0.0307 | 0.0261 | 0.0583 | 0.0393 | 0.0884 | 0.0182 | 0.0261 |
| | Labels | Avg | 0.7545 | 0.8369 | 0.7270 | 0.9374 | 0.7260 | 0.3420 | 0.8175 | 0.7270 |
| | | STD | 0.0559 | 0.0477 | 0.0254 | 0.0403 | 0.0279 | 0.0848 | 0.0214 | 0.0254 |
| | | *P* value | 0.0001 | 0.0091 | 0.0077 | 0.7117 | 0.1429 | 0.0000 | 0.2180 | 0.0077 |

#### 8.2 AUC ROC Curves

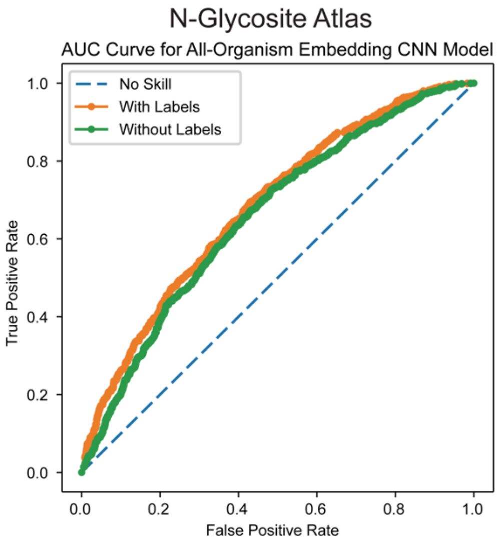


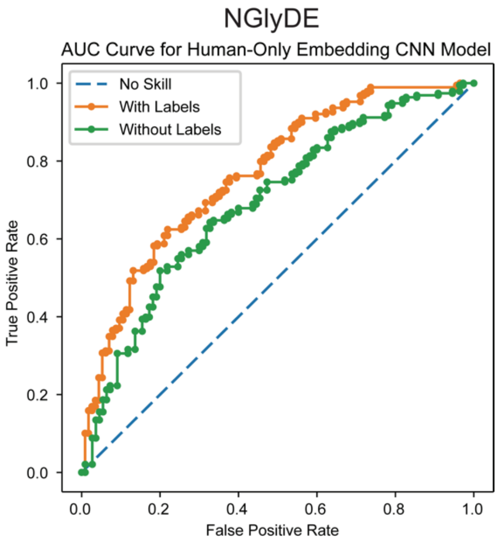

#### 8.3 AUPRC ROC Curves

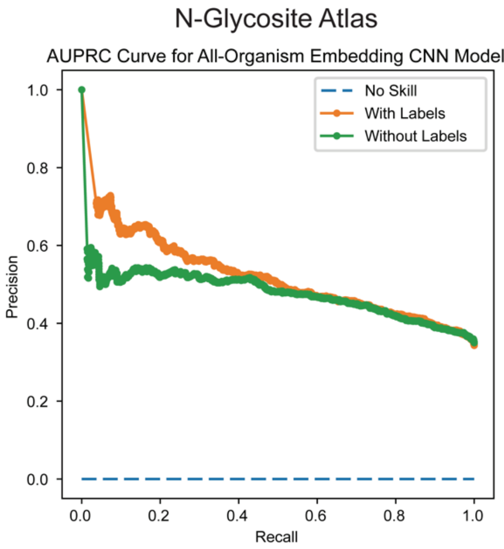


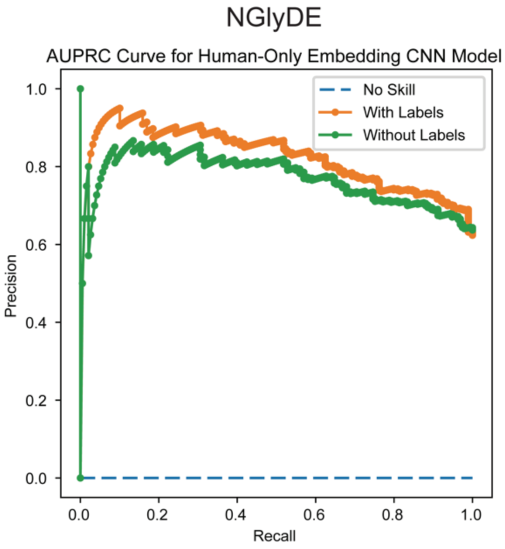


### 9. Additional Kinase-Specific Model Results

#### 9.1. List of Kinases Used

| | | | | | |
|---|---|---|---|---|---|
| AurA | AurC | CDC2/CycB1 | CDK5/p25 | CHK1 | CK1δ |
| CaMK2δ | CaMK2γ | DCLK2 | DYRK2 | DYRK3 | Erk1 |
| Erk2 | Erk5 | GCK | HGK | IKKα | IKKβ |
| IKKε | JNK1 | KHS1 | MAP3K1 | MAPKAPK3 | MAPKAPK5 |
| MARK2 | MARK4 | MINK | MLK1 | MLK2 | MLK3 |
| MSK1 | MST1 | MST2 | MST3 | MST4 | NEK1 |
| NEK2 | NEK4 | NEK6 | NEK7 | NEK9 | NIM1 |
| NLK | p38α | p38β | p38δ | p38γ | p70S6K |
| PAK1 | PAK2 | PAK3 | PHKG2 | PIM1 | PIM3 |
| PKA Cα | PLK1 | PLK3 | QIK | RSK3 | SGK3 |
| SIK | TAK1 | TAO2 | TLK1 | TNIK | TSSK1 |
| TTBK1 | YSK1 | ZAK | | | |

#### 9.2 Kinase-Specific Model Results

| Kinase | | | AUC | AUPRC | Accuracy | Recall | Precision | MCC | F1 | Specificity |
|---|---|---|---|---|---|---|---|---|---|---|
| **AurA** | No labels | Average | 0.8761 | 0.8567 | 0.8188 | 0.8686 | 0.7873 | 0.6428 | 0.8242 | 0.8188 |
| | | STD | 0.0372 | 0.0600 | 0.0457 | 0.0533 | 0.0670 | 0.0880 | 0.0494 | 0.0457 |
| | Labels | Average | 0.9328 | 0.9174 | 0.8594 | 0.8641 | 0.8563 | 0.7187 | 0.8594 | 0.8594 |
| | | STD | 0.0201 | 0.0440 | 0.0239 | 0.0387 | 0.0378 | 0.0491 | 0.0275 | 0.0239 |
| | | *P* value | 0.0013 | 0.0258 | 0.0336 | 0.8415 | 0.0173 | 0.0401 | 0.0826 | 0.0336 |
| **AurC** | No labels | Average | 0.9019 | 0.8788 | 0.8351 | 0.8392 | 0.8430 | 0.6695 | 0.8395 | 0.8351 |
| | | STD | 0.0401 | 0.0504 | 0.0342 | 0.0469 | 0.0599 | 0.0733 | 0.0392 | 0.0342 |
| | Labels | Average | 0.9363 | 0.9299 | 0.8807 | 0.8923 | 0.8793 | 0.7603 | 0.8848 | 0.8807 |
| | | STD | 0.0182 | 0.0308 | 0.0143 | 0.0425 | 0.0276 | 0.0290 | 0.0212 | 0.0143 |
| | | *P* value | 0.0363 | 0.0204 | 0.0030 | 0.0217 | 0.1226 | 0.0049 | 0.0087 | 0.0030 |
| **CDC2/CycB1** | No labels | Average | 0.9256 | 0.9234 | 0.8553 | 0.8461 | 0.8631 | 0.7115 | 0.8529 | 0.8553 |
| | | STD | 0.0198 | 0.0296 | 0.0261 | 0.0466 | 0.0422 | 0.0510 | 0.0246 | 0.0261 |
| | Labels | Average | 0.9605 | 0.9588 | 0.8947 | 0.9020 | 0.8853 | 0.7901 | 0.8917 | 0.8947 |
| | | STD | 0.0116 | 0.0155 | 0.0187 | 0.0610 | 0.0327 | 0.0375 | 0.0293 | 0.0187 |
| | | *P* value | 0.0004 | 0.0069 | 0.0020 | 0.0435 | 0.2283 | 0.0017 | 0.0072 | 0.0020 |
| **CDK5/p25** | No labels | Average | 0.9000 | 0.9016 | 0.8250 | 0.8396 | 0.8133 | 0.6498 | 0.8254 | 0.8250 |
| | | STD | 0.0304 | 0.0337 | 0.0385 | 0.0472 | 0.0542 | 0.0752 | 0.0436 | 0.0385 |
| | Labels | Average | 0.9326 | 0.9310 | 0.8630 | 0.8619 | 0.8797 | 0.7263 | 0.8699 | 0.8630 |
| | | STD | 0.0164 | 0.0183 | 0.0253 | 0.0472 | 0.0157 | 0.0477 | 0.0250 | 0.0253 |
| | | *P* value | 0.0135 | 0.0379 | 0.0254 | 0.3317 | 0.0050 | 0.0209 | 0.0185 | 0.0254 |
| **CHK1** | No labels | Average | 0.8887 | 0.8842 | 0.8000 | 0.8107 | 0.7951 | 0.6079 | 0.7979 | 0.8000 |
| | | STD | 0.0189 | 0.0325 | 0.0327 | 0.0946 | 0.0481 | 0.0627 | 0.0422 | 0.0327 |
| | Labels | Average | 0.9376 | 0.9348 | 0.8663 | 0.8736 | 0.8686 | 0.7343 | 0.8693 | 0.8663 |
| | | STD | 0.0224 | 0.0305 | 0.0356 | 0.0622 | 0.0448 | 0.0700 | 0.0381 | 0.0356 |
| | | *P* value | 0.0001 | 0.0032 | 0.0007 | 0.1158 | 0.0036 | 0.0008 | 0.0014 | 0.0007 |
| **CK1δ** | No labels | Average | 0.9106 | 0.8950 | 0.8383 | 0.8602 | 0.8222 | 0.6781 | 0.8388 | 0.8383 |
| | | STD | 0.0179 | 0.0296 | 0.0215 | 0.0557 | 0.0407 | 0.0411 | 0.0277 | 0.0215 |

| | | | | | | | | | | |
|---|---|---|---|---|---|---|---|---|---|---|
| | Labels | Average | 0.9552 | 0.9546 | 0.8750 | 0.8933 | 0.8608 | 0.7527 | 0.8752 | 0.8750 |
| | | STD | 0.0091 | 0.0127 | 0.0255 | 0.0525 | 0.0464 | 0.0521 | 0.0320 | 0.0255 |
| | | *P* value | 0.0000 | 0.0001 | 0.0041 | 0.2121 | 0.0770 | 0.0036 | 0.0192 | 0.0041 |
| **CaMK2δ** | No labels | Average | 0.8935 | 0.8865 | 0.8144 | 0.8452 | 0.8047 | 0.6299 | 0.8230 | 0.8144 |
| | | STD | 0.0301 | 0.0339 | 0.0445 | 0.0701 | 0.0391 | 0.0905 | 0.0452 | 0.0445 |
| | Labels | Average | 0.9385 | 0.9296 | 0.8536 | 0.8800 | 0.8372 | 0.7105 | 0.8561 | 0.8536 |
| | | STD | 0.0188 | 0.0341 | 0.0302 | 0.0553 | 0.0536 | 0.0589 | 0.0369 | 0.0302 |
| | | *P* value | 0.0017 | 0.0151 | 0.0441 | 0.2579 | 0.1605 | 0.0403 | 0.1065 | 0.0441 |
| **CaMK2γ** | No labels | Average | 0.9034 | 0.8875 | 0.8235 | 0.8432 | 0.8076 | 0.6468 | 0.8234 | 0.8235 |
| | | STD | 0.0250 | 0.0472 | 0.0264 | 0.0527 | 0.0265 | 0.0512 | 0.0219 | 0.0264 |
| | Labels | Average | 0.9466 | 0.9392 | 0.8783 | 0.8734 | 0.8851 | 0.7545 | 0.8787 | 0.8783 |
| | | STD | 0.0228 | 0.0311 | 0.0346 | 0.0346 | 0.0515 | 0.0683 | 0.0382 | 0.0346 |
| | | *P* value | 0.0013 | 0.0147 | 0.0015 | 0.1702 | 0.0014 | 0.0015 | 0.0020 | 0.0015 |
| **DCLK2** | No labels | Average | 0.8869 | 0.8717 | 0.7969 | 0.8308 | 0.7670 | 0.5988 | 0.7951 | 0.7969 |
| | | STD | 0.0338 | 0.0594 | 0.0391 | 0.0560 | 0.0629 | 0.0787 | 0.0410 | 0.0391 |
| | Labels | Average | 0.9365 | 0.9367 | 0.8670 | 0.8709 | 0.8735 | 0.7380 | 0.8693 | 0.8670 |
| | | STD | 0.0241 | 0.0302 | 0.0403 | 0.0839 | 0.0380 | 0.0779 | 0.0439 | 0.0403 |
| | | *P* value | 0.0025 | 0.0115 | 0.0015 | 0.2508 | 0.0006 | 0.0014 | 0.0016 | 0.0015 |
| **DYRK2** | No labels | Average | 0.8718 | 0.8614 | 0.7968 | 0.8224 | 0.7794 | 0.5926 | 0.7995 | 0.7968 |
| | | STD | 0.0195 | 0.0318 | 0.0212 | 0.0358 | 0.0327 | 0.0442 | 0.0221 | 0.0212 |
| | Labels | Average | 0.9284 | 0.9211 | 0.8552 | 0.8693 | 0.8479 | 0.7131 | 0.8569 | 0.8552 |
| | | STD | 0.0220 | 0.0320 | 0.0305 | 0.0450 | 0.0528 | 0.0580 | 0.0334 | 0.0305 |
| | | *P* value | 0.0000 | 0.0009 | 0.0002 | 0.0255 | 0.0048 | 0.0001 | 0.0006 | 0.0002 |
| **DYRK3** | No labels | Average | 0.8831 | 0.8626 | 0.7990 | 0.7969 | 0.7950 | 0.6015 | 0.7920 | 0.7990 |
| | | STD | 0.0294 | 0.0488 | 0.0345 | 0.0660 | 0.0620 | 0.0642 | 0.0375 | 0.0345 |
| | Labels | Average | 0.9305 | 0.9087 | 0.8631 | 0.8918 | 0.8403 | 0.7309 | 0.8628 | 0.8631 |
| | | STD | 0.0286 | 0.0530 | 0.0273 | 0.0416 | 0.0614 | 0.0541 | 0.0243 | 0.0273 |
| | | *P* value | 0.0027 | 0.0712 | 0.0004 | 0.0023 | 0.1367 | 0.0002 | 0.0002 | 0.0004 |
| **Erk1** | No labels | Average | 0.8887 | 0.8773 | 0.8158 | 0.8212 | 0.8117 | 0.6336 | 0.8134 | 0.8158 |
| | | STD | 0.0350 | 0.0409 | 0.0372 | 0.0635 | 0.0601 | 0.0737 | 0.0397 | 0.0372 |

| | | | | | | | | | | |
|---|---|---|---|---|---|---|---|---|---|---|
| | Labels | Average | 0.9361 | 0.9236 | 0.8579 | 0.8860 | 0.8348 | 0.7229 | 0.8566 | 0.8579 |
| | | STD | 0.0254 | 0.0344 | 0.0382 | 0.0543 | 0.0718 | 0.0741 | 0.0396 | 0.0382 |
| | | *P* value | 0.0045 | 0.0185 | 0.0293 | 0.0323 | 0.4684 | 0.0196 | 0.0328 | 0.0293 |
| **Erk2** | No labels | Average | 0.9020 | 0.9061 | 0.8341 | 0.8456 | 0.8338 | 0.6728 | 0.8367 | 0.8341 |
| | | STD | 0.0174 | 0.0158 | 0.0170 | 0.0617 | 0.0420 | 0.0321 | 0.0212 | 0.0170 |
| | Labels | Average | 0.9539 | 0.9475 | 0.8948 | 0.8946 | 0.8905 | 0.7888 | 0.8918 | 0.8948 |
| | | STD | 0.0282 | 0.0304 | 0.0361 | 0.0290 | 0.0525 | 0.0726 | 0.0334 | 0.0361 |
| | | *P* value | 0.0003 | 0.0029 | 0.0005 | 0.0502 | 0.0214 | 0.0008 | 0.0008 | 0.0005 |
| **Erk5** | No labels | Average | 0.8505 | 0.8470 | 0.7802 | 0.7776 | 0.7960 | 0.5639 | 0.7848 | 0.7802 |
| | | STD | 0.0465 | 0.0433 | 0.0420 | 0.0734 | 0.0282 | 0.0805 | 0.0420 | 0.0420 |
| | Labels | Average | 0.9298 | 0.9233 | 0.8490 | 0.8443 | 0.8639 | 0.6997 | 0.8508 | 0.8490 |
| | | STD | 0.0238 | 0.0361 | 0.0376 | 0.0794 | 0.0543 | 0.0738 | 0.0462 | 0.0376 |
| | | *P* value | 0.0005 | 0.0008 | 0.0018 | 0.0810 | 0.0052 | 0.0015 | 0.0053 | 0.0018 |
| **GCK** | No labels | Average | 0.8565 | 0.8520 | 0.7564 | 0.7788 | 0.7489 | 0.5160 | 0.7610 | 0.7564 |
| | | STD | 0.0318 | 0.0396 | 0.0266 | 0.0526 | 0.0520 | 0.0523 | 0.0288 | 0.0266 |
| | Labels | Average | 0.9169 | 0.9107 | 0.8336 | 0.8151 | 0.8505 | 0.6703 | 0.8306 | 0.8336 |
| | | STD | 0.0130 | 0.0260 | 0.0173 | 0.0494 | 0.0464 | 0.0363 | 0.0271 | 0.0173 |
| | | *P* value | 0.0002 | 0.0020 | 0.0000 | 0.1493 | 0.0004 | 0.0000 | 0.0001 | 0.0000 |
| **HGK** | No labels | Average | 0.8761 | 0.8652 | 0.7877 | 0.7797 | 0.7919 | 0.5725 | 0.7852 | 0.7877 |
| | | STD | 0.0256 | 0.0452 | 0.0299 | 0.0494 | 0.0415 | 0.0610 | 0.0400 | 0.0299 |
| | Labels | Average | 0.9416 | 0.9407 | 0.8673 | 0.8666 | 0.8764 | 0.7384 | 0.8692 | 0.8673 |
| | | STD | 0.0201 | 0.0271 | 0.0299 | 0.0673 | 0.0312 | 0.0541 | 0.0307 | 0.0299 |
| | | *P* value | 0.0000 | 0.0007 | 0.0000 | 0.0064 | 0.0001 | 0.0000 | 0.0001 | 0.0000 |
| **IKKα** | No labels | Average | 0.8519 | 0.8389 | 0.7826 | 0.7970 | 0.7650 | 0.5711 | 0.7771 | 0.7826 |
| | | STD | 0.0367 | 0.0422 | 0.0466 | 0.1010 | 0.0354 | 0.0928 | 0.0566 | 0.0466 |
| | Labels | Average | 0.9337 | 0.9214 | 0.8554 | 0.8708 | 0.8372 | 0.7131 | 0.8521 | 0.8554 |
| | | STD | 0.0255 | 0.0338 | 0.0367 | 0.0667 | 0.0401 | 0.0756 | 0.0395 | 0.0367 |
| | | *P* value | 0.0000 | 0.0003 | 0.0018 | 0.0866 | 0.0008 | 0.0024 | 0.0049 | 0.0018 |
| **IKKβ** | No labels | Average | 0.8565 | 0.8291 | 0.7757 | 0.7968 | 0.7579 | 0.5592 | 0.7728 | 0.7757 |
| | | STD | 0.0191 | 0.0337 | 0.0308 | 0.0847 | 0.0502 | 0.0626 | 0.0398 | 0.0308 |

| | | | | | | | | | | |
|---|---|---|---|---|---|---|---|---|---|---|
| | Labels | Average | 0.9442 | 0.9410 | 0.8739 | 0.8884 | 0.8614 | 0.7485 | 0.8741 | 0.8739 |
| | | STD | 0.0095 | 0.0157 | 0.0169 | 0.0267 | 0.0282 | 0.0339 | 0.0150 | 0.0169 |
| | | *P* value | 0.0000 | 0.0000 | 0.0000 | 0.0104 | 0.0001 | 0.0000 | 0.0000 | 0.0000 |
| **IKKε** | No labels | Average | 0.8719 | 0.8609 | 0.7940 | 0.7843 | 0.7982 | 0.5895 | 0.7901 | 0.7940 |
| | | STD | 0.0240 | 0.0360 | 0.0341 | 0.0565 | 0.0419 | 0.0676 | 0.0406 | 0.0341 |
| | Labels | Average | 0.9276 | 0.9247 | 0.8500 | 0.8717 | 0.8405 | 0.7020 | 0.8545 | 0.8500 |
| | | STD | 0.0241 | 0.0244 | 0.0365 | 0.0384 | 0.0494 | 0.0725 | 0.0293 | 0.0365 |
| | | *P* value | 0.0001 | 0.0005 | 0.0035 | 0.0015 | 0.0661 | 0.0032 | 0.0013 | 0.0035 |
| **JNK1** | No labels | Average | 0.9040 | 0.8990 | 0.8357 | 0.8332 | 0.8449 | 0.6750 | 0.8368 | 0.8357 |
| | | STD | 0.0272 | 0.0349 | 0.0324 | 0.0383 | 0.0651 | 0.0642 | 0.0318 | 0.0324 |
| | Labels | Average | 0.9499 | 0.9361 | 0.8818 | 0.9196 | 0.8481 | 0.7676 | 0.8814 | 0.8818 |
| | | STD | 0.0137 | 0.0233 | 0.0178 | 0.0241 | 0.0411 | 0.0331 | 0.0191 | 0.0178 |
| | | *P* value | 0.0005 | 0.0176 | 0.0022 | 0.0000 | 0.9042 | 0.0019 | 0.0027 | 0.0022 |
| **KHS1** | No labels | Average | 0.8514 | 0.8280 | 0.7708 | 0.7898 | 0.7620 | 0.5452 | 0.7737 | 0.7708 |
| | | STD | 0.0183 | 0.0410 | 0.0178 | 0.0384 | 0.0461 | 0.0340 | 0.0175 | 0.0178 |
| | Labels | Average | 0.9401 | 0.9315 | 0.8639 | 0.8801 | 0.8523 | 0.7294 | 0.8647 | 0.8639 |
| | | STD | 0.0185 | 0.0257 | 0.0305 | 0.0407 | 0.0440 | 0.0585 | 0.0275 | 0.0305 |
| | | *P* value | 0.0000 | 0.0000 | 0.0000 | 0.0001 | 0.0005 | 0.0000 | 0.0000 | 0.0000 |
| **MAP3K1** | No labels | Average | 0.8680 | 0.8449 | 0.7827 | 0.8001 | 0.7549 | 0.5689 | 0.7746 | 0.7827 |
| | | STD | 0.0235 | 0.0447 | 0.0283 | 0.0490 | 0.0676 | 0.0576 | 0.0423 | 0.0283 |
| | Labels | Average | 0.9182 | 0.9075 | 0.8327 | 0.8256 | 0.8339 | 0.6672 | 0.8274 | 0.8327 |
| | | STD | 0.0270 | 0.0428 | 0.0328 | 0.0450 | 0.0684 | 0.0634 | 0.0376 | 0.0328 |
| | | *P* value | 0.0005 | 0.0072 | 0.0028 | 0.2652 | 0.0239 | 0.0029 | 0.0120 | 0.0028 |
| **MAPKAPK3** | No labels | Average | 0.8773 | 0.8784 | 0.7985 | 0.7854 | 0.8164 | 0.5995 | 0.7979 | 0.7985 |
| | | STD | 0.0179 | 0.0347 | 0.0260 | 0.0624 | 0.0509 | 0.0508 | 0.0344 | 0.0260 |
| | Labels | Average | 0.9192 | 0.9157 | 0.8466 | 0.8574 | 0.8380 | 0.6946 | 0.8460 | 0.8466 |
| | | STD | 0.0245 | 0.0292 | 0.0367 | 0.0662 | 0.0440 | 0.0729 | 0.0428 | 0.0367 |
| | | *P* value | 0.0007 | 0.0245 | 0.0054 | 0.0289 | 0.3481 | 0.0054 | 0.0174 | 0.0054 |
| **MAPKAPK5** | No labels | Average | 0.8106 | 0.7890 | 0.7292 | 0.7674 | 0.7066 | 0.4652 | 0.7326 | 0.7292 |
| | | STD | 0.0445 | 0.0741 | 0.0411 | 0.0887 | 0.0521 | 0.0873 | 0.0529 | 0.0411 |

| | | | | | | | | | | |
|---|---|---|---|---|---|---|---|---|---|---|
| | Labels | Average | 0.8845 | 0.8879 | 0.8104 | 0.8258 | 0.8149 | 0.6226 | 0.8180 | 0.8104 |
| | | STD | 0.0395 | 0.0408 | 0.0596 | 0.0729 | 0.0758 | 0.1189 | 0.0602 | 0.0596 |
| | | *P* value | 0.0016 | 0.0035 | 0.0039 | 0.1450 | 0.0028 | 0.0054 | 0.0051 | 0.0039 |
| **MARK2** | No labels | Average | 0.8486 | 0.8293 | 0.7648 | 0.7531 | 0.7778 | 0.5299 | 0.7635 | 0.7648 |
| | | STD | 0.0402 | 0.0581 | 0.0467 | 0.0729 | 0.0588 | 0.0955 | 0.0546 | 0.0467 |
| | Labels | Average | 0.9363 | 0.9300 | 0.8648 | 0.8841 | 0.8453 | 0.7316 | 0.8617 | 0.8648 |
| | | STD | 0.0199 | 0.0234 | 0.0236 | 0.0456 | 0.0557 | 0.0442 | 0.0215 | 0.0236 |
| | | *P* value | 0.0001 | 0.0004 | 0.0001 | 0.0004 | 0.0223 | 0.0001 | 0.0003 | 0.0001 |
| **MARK4** | No labels | Average | 0.8425 | 0.8266 | 0.7645 | 0.7587 | 0.7700 | 0.5270 | 0.7635 | 0.7645 |
| | | STD | 0.0366 | 0.0539 | 0.0365 | 0.0380 | 0.0541 | 0.0756 | 0.0385 | 0.0365 |
| | Labels | Average | 0.9062 | 0.8895 | 0.8255 | 0.8469 | 0.8082 | 0.6573 | 0.8232 | 0.8255 |
| | | STD | 0.0346 | 0.0477 | 0.0383 | 0.0595 | 0.0711 | 0.0733 | 0.0365 | 0.0383 |
| | | *P* value | 0.0013 | 0.0176 | 0.0028 | 0.0019 | 0.2172 | 0.0016 | 0.0033 | 0.0028 |
| **MINK** | No labels | Average | 0.8495 | 0.8345 | 0.7732 | 0.7726 | 0.7880 | 0.5488 | 0.7784 | 0.7732 |
| | | STD | 0.0327 | 0.0577 | 0.0252 | 0.0497 | 0.0470 | 0.0495 | 0.0299 | 0.0252 |
| | Labels | Average | 0.9415 | 0.9391 | 0.8631 | 0.8615 | 0.8595 | 0.7263 | 0.8600 | 0.8631 |
| | | STD | 0.0230 | 0.0252 | 0.0394 | 0.0515 | 0.0383 | 0.0786 | 0.0404 | 0.0394 |
| | | *P* value | 0.0000 | 0.0003 | 0.0000 | 0.0015 | 0.0025 | 0.0000 | 0.0002 | 0.0000 |
| **MLK1** | No labels | Average | 0.8655 | 0.8473 | 0.7880 | 0.7818 | 0.7861 | 0.5781 | 0.7820 | 0.7880 |
| | | STD | 0.0208 | 0.0356 | 0.0254 | 0.0542 | 0.0405 | 0.0503 | 0.0288 | 0.0254 |
| | Labels | Average | 0.9446 | 0.9400 | 0.8770 | 0.8845 | 0.8659 | 0.7553 | 0.8743 | 0.8770 |
| | | STD | 0.0186 | 0.0178 | 0.0310 | 0.0532 | 0.0261 | 0.0622 | 0.0328 | 0.0310 |
| | | *P* value | 0.0000 | 0.0000 | 0.0000 | 0.0007 | 0.0002 | 0.0000 | 0.0000 | 0.0000 |
| **MLK2** | No labels | Average | 0.8853 | 0.8768 | 0.8009 | 0.8502 | 0.7698 | 0.6041 | 0.8065 | 0.8009 |
| | | STD | 0.0196 | 0.0268 | 0.0376 | 0.0483 | 0.0660 | 0.0712 | 0.0477 | 0.0376 |
| | Labels | Average | 0.9453 | 0.9506 | 0.8698 | 0.8827 | 0.8806 | 0.7409 | 0.8794 | 0.8698 |
| | | STD | 0.0166 | 0.0193 | 0.0352 | 0.0425 | 0.0639 | 0.0696 | 0.0329 | 0.0352 |
| | | *P* value | 0.0000 | 0.0000 | 0.0008 | 0.1473 | 0.0020 | 0.0006 | 0.0016 | 0.0008 |
| **MLK3** | No labels | Average | 0.8329 | 0.8089 | 0.7563 | 0.7458 | 0.7745 | 0.5141 | 0.7569 | 0.7563 |
| | | STD | 0.0345 | 0.0512 | 0.0414 | 0.0842 | 0.0464 | 0.0797 | 0.0530 | 0.0414 |

| | | | | | | | | | | |
|---|---|---|---|---|---|---|---|---|---|---|
| | Labels | Average | 0.9155 | 0.9116 | 0.8373 | 0.8343 | 0.8349 | 0.6758 | 0.8328 | 0.8373 |
| | | STD | 0.0200 | 0.0252 | 0.0257 | 0.0504 | 0.0487 | 0.0508 | 0.0312 | 0.0257 |
| | | *P* value | 0.0000 | 0.0001 | 0.0002 | 0.0165 | 0.0148 | 0.0001 | 0.0022 | 0.0002 |
| **MSK1** | No labels | Average | 0.8951 | 0.8925 | 0.8237 | 0.7979 | 0.8378 | 0.6498 | 0.8134 | 0.8237 |
| | | STD | 0.0315 | 0.0446 | 0.0403 | 0.0781 | 0.0655 | 0.0832 | 0.0459 | 0.0403 |
| | Labels | Average | 0.9335 | 0.9329 | 0.8535 | 0.8398 | 0.8684 | 0.7092 | 0.8523 | 0.8535 |
| | | STD | 0.0191 | 0.0278 | 0.0204 | 0.0367 | 0.0458 | 0.0409 | 0.0194 | 0.0204 |
| | | *P* value | 0.0070 | 0.0356 | 0.0686 | 0.1693 | 0.2672 | 0.0765 | 0.0369 | 0.0686 |
| **MST1** | No labels | Average | 0.8858 | 0.8665 | 0.8101 | 0.8249 | 0.8026 | 0.6216 | 0.8122 | 0.8101 |
| | | STD | 0.0304 | 0.0406 | 0.0274 | 0.0465 | 0.0449 | 0.0544 | 0.0313 | 0.0274 |
| | Labels | Average | 0.9536 | 0.9538 | 0.8799 | 0.8756 | 0.8874 | 0.7606 | 0.8804 | 0.8799 |
| | | STD | 0.0122 | 0.0141 | 0.0227 | 0.0376 | 0.0358 | 0.0460 | 0.0206 | 0.0227 |
| | | *P* value | 0.0000 | 0.0001 | 0.0000 | 0.0209 | 0.0004 | 0.0000 | 0.0001 | 0.0000 |
| **MST2** | No labels | Average | 0.8727 | 0.8411 | 0.7856 | 0.7971 | 0.7691 | 0.5718 | 0.7808 | 0.7856 |
| | | STD | 0.0203 | 0.0364 | 0.0306 | 0.0699 | 0.0446 | 0.0632 | 0.0421 | 0.0306 |
| | Labels | Average | 0.9470 | 0.9459 | 0.8673 | 0.8760 | 0.8572 | 0.7354 | 0.8657 | 0.8673 |
| | | STD | 0.0115 | 0.0138 | 0.0209 | 0.0359 | 0.0298 | 0.0406 | 0.0215 | 0.0209 |
| | | *P* value | 0.0000 | 0.0000 | 0.0000 | 0.0097 | 0.0002 | 0.0000 | 0.0001 | 0.0000 |
| **MST3** | No labels | Average | 0.8925 | 0.8760 | 0.8119 | 0.8221 | 0.8034 | 0.6234 | 0.8116 | 0.8119 |
| | | STD | 0.0202 | 0.0302 | 0.0260 | 0.0555 | 0.0268 | 0.0532 | 0.0311 | 0.0260 |
| | Labels | Average | 0.9558 | 0.9490 | 0.8863 | 0.9046 | 0.8735 | 0.7743 | 0.8877 | 0.8863 |
| | | STD | 0.0107 | 0.0229 | 0.0266 | 0.0371 | 0.0480 | 0.0533 | 0.0306 | 0.0266 |
| | | *P* value | 0.0000 | 0.0000 | 0.0000 | 0.0020 | 0.0018 | 0.0000 | 0.0001 | 0.0000 |
| **MST4** | No labels | Average | 0.8842 | 0.8712 | 0.8098 | 0.8365 | 0.7917 | 0.6209 | 0.8124 | 0.8098 |
| | | STD | 0.0249 | 0.0383 | 0.0351 | 0.0264 | 0.0608 | 0.0677 | 0.0381 | 0.0351 |
| | Labels | Average | 0.9218 | 0.9208 | 0.8434 | 0.8634 | 0.8312 | 0.6870 | 0.8466 | 0.8434 |
| | | STD | 0.0318 | 0.0309 | 0.0334 | 0.0309 | 0.0376 | 0.0673 | 0.0286 | 0.0334 |
| | | *P* value | 0.0122 | 0.0076 | 0.0523 | 0.0627 | 0.1177 | 0.0524 | 0.0467 | 0.0523 |
| **NEK1** | No labels | Average | 0.8532 | 0.8505 | 0.7599 | 0.7892 | 0.7498 | 0.5226 | 0.7665 | 0.7599 |
| | | STD | 0.0301 | 0.0372 | 0.0381 | 0.0609 | 0.0650 | 0.0711 | 0.0473 | 0.0381 |

| | | | | | | | | | | |
|---|---|---|---|---|---|---|---|---|---|---|
| | Labels | Average | 0.9289 | 0.9333 | 0.8472 | 0.8482 | 0.8538 | 0.6940 | 0.8493 | 0.8472 |
| | | STD | 0.0150 | 0.0191 | 0.0223 | 0.0590 | 0.0326 | 0.0450 | 0.0298 | 0.0223 |
| | | *P* value | 0.0000 | 0.0000 | 0.0000 | 0.0516 | 0.0008 | 0.0000 | 0.0005 | 0.0000 |
| **NEK2** | No labels | Average | 0.8489 | 0.8246 | 0.7695 | 0.8187 | 0.7519 | 0.5471 | 0.7801 | 0.7695 |
| | | STD | 0.0292 | 0.0494 | 0.0237 | 0.0664 | 0.0519 | 0.0509 | 0.0221 | 0.0237 |
| | Labels | Average | 0.9383 | 0.9362 | 0.8715 | 0.8766 | 0.8690 | 0.7433 | 0.8715 | 0.8715 |
| | | STD | 0.0207 | 0.0282 | 0.0273 | 0.0351 | 0.0469 | 0.0559 | 0.0261 | 0.0273 |
| | | *P* value | 0.0000 | 0.0000 | 0.0000 | 0.0369 | 0.0001 | 0.0000 | 0.0000 | 0.0000 |
| **NEK4** | No labels | Average | 0.8479 | 0.8070 | 0.7667 | 0.8013 | 0.7499 | 0.5355 | 0.7731 | 0.7667 |
| | | STD | 0.0295 | 0.0603 | 0.0259 | 0.0597 | 0.0389 | 0.0544 | 0.0335 | 0.0259 |
| | Labels | Average | 0.9494 | 0.9497 | 0.8761 | 0.8830 | 0.8729 | 0.7547 | 0.8765 | 0.8761 |
| | | STD | 0.0144 | 0.0150 | 0.0278 | 0.0528 | 0.0370 | 0.0548 | 0.0287 | 0.0278 |
| | | *P* value | 0.0000 | 0.0000 | 0.0000 | 0.0066 | 0.0000 | 0.0000 | 0.0000 | 0.0000 |
| **NEK6** | No labels | Average | 0.8718 | 0.8614 | 0.8000 | 0.8223 | 0.7941 | 0.6028 | 0.8063 | 0.8000 |
| | | STD | 0.0178 | 0.0330 | 0.0239 | 0.0517 | 0.0357 | 0.0483 | 0.0257 | 0.0239 |
| | Labels | Average | 0.9309 | 0.9287 | 0.8576 | 0.8727 | 0.8476 | 0.7160 | 0.8590 | 0.8576 |
| | | STD | 0.0170 | 0.0231 | 0.0274 | 0.0438 | 0.0379 | 0.0540 | 0.0292 | 0.0274 |
| | | *P* value | 0.0000 | 0.0001 | 0.0002 | 0.0389 | 0.0065 | 0.0002 | 0.0008 | 0.0002 |
| **NEK7** | No labels | Average | 0.8790 | 0.8562 | 0.7876 | 0.8031 | 0.7703 | 0.5776 | 0.7844 | 0.7876 |
| | | STD | 0.0115 | 0.0273 | 0.0153 | 0.0606 | 0.0275 | 0.0346 | 0.0244 | 0.0153 |
| | Labels | Average | 0.9214 | 0.9158 | 0.8444 | 0.8587 | 0.8316 | 0.6889 | 0.8442 | 0.8444 |
| | | STD | 0.0264 | 0.0315 | 0.0263 | 0.0465 | 0.0251 | 0.0543 | 0.0274 | 0.0263 |
| | | *P* value | 0.0008 | 0.0005 | 0.0001 | 0.0436 | 0.0001 | 0.0001 | 0.0001 | 0.0001 |
| **NEK9** | No labels | Average | 0.8665 | 0.8406 | 0.7963 | 0.8228 | 0.7546 | 0.5943 | 0.7848 | 0.7963 |
| | | STD | 0.0336 | 0.0531 | 0.0301 | 0.0605 | 0.0611 | 0.0610 | 0.0418 | 0.0301 |
| | Labels | Average | 0.9423 | 0.9369 | 0.8713 | 0.8642 | 0.8700 | 0.7431 | 0.8654 | 0.8713 |
| | | STD | 0.0193 | 0.0178 | 0.0250 | 0.0454 | 0.0380 | 0.0520 | 0.0189 | 0.0250 |
| | | *P* value | 0.0000 | 0.0003 | 0.0000 | 0.1201 | 0.0002 | 0.0000 | 0.0002 | 0.0000 |
| **NIM1** | No labels | Average | 0.8576 | 0.8536 | 0.7733 | 0.7803 | 0.7780 | 0.5513 | 0.7764 | 0.7733 |
| | | STD | 0.0450 | 0.0423 | 0.0552 | 0.0915 | 0.0444 | 0.1083 | 0.0571 | 0.0552 |

| | | | | | | | | | | |
|---|---|---|---|---|---|---|---|---|---|---|
| | Labels | Average | 0.9063 | 0.8926 | 0.8305 | 0.8764 | 0.8115 | 0.6677 | 0.8403 | 0.8305 |
| | | STD | 0.0270 | 0.0427 | 0.0204 | 0.0567 | 0.0413 | 0.0407 | 0.0202 | 0.0204 |
| | | *P* value | 0.0141 | 0.0668 | 0.0134 | 0.0172 | 0.1146 | 0.0111 | 0.0088 | 0.0134 |
| **NLK** | No labels | Average | 0.8582 | 0.8309 | 0.7756 | 0.7539 | 0.7646 | 0.5544 | 0.7545 | 0.7756 |
| | | STD | 0.0363 | 0.0628 | 0.0449 | 0.0770 | 0.0870 | 0.0912 | 0.0555 | 0.0449 |
| | Labels | Average | 0.9261 | 0.9176 | 0.8444 | 0.8676 | 0.8259 | 0.6931 | 0.8438 | 0.8444 |
| | | STD | 0.0183 | 0.0199 | 0.0268 | 0.0564 | 0.0427 | 0.0529 | 0.0226 | 0.0268 |
| | | *P* value | 0.0002 | 0.0024 | 0.0013 | 0.0024 | 0.0800 | 0.0014 | 0.0008 | 0.0013 |
| **PAK1** | No labels | Average | 0.9184 | 0.9181 | 0.8207 | 0.8324 | 0.8233 | 0.6530 | 0.8207 | 0.8207 |
| | | STD | 0.0237 | 0.0243 | 0.0358 | 0.0847 | 0.0787 | 0.0524 | 0.0360 | 0.0358 |
| | Labels | Average | 0.9335 | 0.9337 | 0.8565 | 0.8371 | 0.8783 | 0.7171 | 0.8551 | 0.8565 |
| | | STD | 0.0327 | 0.0370 | 0.0256 | 0.0546 | 0.0415 | 0.0494 | 0.0262 | 0.0256 |
| | | *P* value | 0.2800 | 0.3037 | 0.0263 | 0.8905 | 0.0853 | 0.0155 | 0.0337 | 0.0263 |
| **PAK2** | No labels | Average | 0.8856 | 0.8670 | 0.8051 | 0.7974 | 0.7975 | 0.6107 | 0.7949 | 0.8051 |
| | | STD | 0.0288 | 0.0478 | 0.0501 | 0.0726 | 0.0722 | 0.0982 | 0.0583 | 0.0501 |
| | Labels | Average | 0.9351 | 0.9274 | 0.8500 | 0.8524 | 0.8349 | 0.7006 | 0.8419 | 0.8500 |
| | | STD | 0.0220 | 0.0267 | 0.0401 | 0.0570 | 0.0600 | 0.0800 | 0.0448 | 0.0401 |
| | | *P* value | 0.0008 | 0.0051 | 0.0508 | 0.0915 | 0.2478 | 0.0477 | 0.0722 | 0.0508 |
| **PAK3** | No labels | Average | 0.8996 | 0.8990 | 0.7976 | 0.7916 | 0.8043 | 0.5921 | 0.7972 | 0.7976 |
| | | STD | 0.0210 | 0.0277 | 0.0248 | 0.0395 | 0.0444 | 0.0489 | 0.0346 | 0.0248 |
| | Labels | Average | 0.9306 | 0.9280 | 0.8532 | 0.8602 | 0.8479 | 0.7078 | 0.8529 | 0.8532 |
| | | STD | 0.0266 | 0.0288 | 0.0344 | 0.0571 | 0.0385 | 0.0704 | 0.0362 | 0.0344 |
| | | *P* value | 0.0139 | 0.0432 | 0.0011 | 0.0092 | 0.0394 | 0.0009 | 0.0037 | 0.0011 |
| **PHKG2** | No labels | Average | 0.8648 | 0.8361 | 0.7779 | 0.7702 | 0.7637 | 0.5529 | 0.7657 | 0.7779 |
| | | STD | 0.0368 | 0.0592 | 0.0395 | 0.0441 | 0.0674 | 0.0809 | 0.0475 | 0.0395 |
| | Labels | Average | 0.9362 | 0.9326 | 0.8654 | 0.8866 | 0.8524 | 0.7338 | 0.8674 | 0.8654 |
| | | STD | 0.0277 | 0.0386 | 0.0325 | 0.0626 | 0.0447 | 0.0630 | 0.0386 | 0.0325 |
| | | *P* value | 0.0002 | 0.0009 | 0.0001 | 0.0003 | 0.0048 | 0.0001 | 0.0001 | 0.0001 |
| **PIM1** | No labels | Average | 0.8863 | 0.8642 | 0.8048 | 0.8207 | 0.7963 | 0.6102 | 0.8060 | 0.8048 |
| | | STD | 0.0310 | 0.0493 | 0.0404 | 0.0482 | 0.0730 | 0.0810 | 0.0471 | 0.0404 |

| | | | | | | | | | | |
|---|---|---|---|---|---|---|---|---|---|---|
| | Labels | Average | 0.9347 | 0.9265 | 0.8587 | 0.8733 | 0.8536 | 0.7184 | 0.8623 | 0.8587 |
| | | STD | 0.0242 | 0.0439 | 0.0352 | 0.0353 | 0.0568 | 0.0690 | 0.0367 | 0.0352 |
| | | *P* value | 0.0018 | 0.0112 | 0.0075 | 0.0176 | 0.0803 | 0.0071 | 0.0117 | 0.0075 |
| **PIM3** | No labels | Average | 0.8731 | 0.8525 | 0.7862 | 0.8517 | 0.7481 | 0.5801 | 0.7943 | 0.7862 |
| | | STD | 0.0366 | 0.0608 | 0.0472 | 0.0552 | 0.0671 | 0.0918 | 0.0475 | 0.0472 |
| | Labels | Average | 0.9285 | 0.9276 | 0.8500 | 0.8742 | 0.8431 | 0.7020 | 0.8569 | 0.8500 |
| | | STD | 0.0340 | 0.0348 | 0.0467 | 0.0486 | 0.0598 | 0.0925 | 0.0429 | 0.0467 |
| | | *P* value | 0.0038 | 0.0060 | 0.0099 | 0.3718 | 0.0054 | 0.0117 | 0.0089 | 0.0099 |
| **PKA Cα** | No labels | Average | 0.9192 | 0.9095 | 0.8407 | 0.8463 | 0.8442 | 0.6831 | 0.8435 | 0.8407 |
| | | STD | 0.0302 | 0.0355 | 0.0387 | 0.0546 | 0.0533 | 0.0765 | 0.0387 | 0.0387 |
| | Labels | Average | 0.9316 | 0.9188 | 0.8626 | 0.8741 | 0.8590 | 0.7276 | 0.8646 | 0.8626 |
| | | STD | 0.0190 | 0.0342 | 0.0276 | 0.0483 | 0.0547 | 0.0529 | 0.0346 | 0.0276 |
| | | *P* value | 0.3120 | 0.5801 | 0.1843 | 0.2668 | 0.5687 | 0.1701 | 0.2375 | 0.1843 |
| **PLK1** | No labels | Average | 0.9109 | 0.9067 | 0.8398 | 0.8502 | 0.8456 | 0.6814 | 0.8455 | 0.8398 |
| | | STD | 0.0311 | 0.0429 | 0.0299 | 0.0553 | 0.0594 | 0.0614 | 0.0355 | 0.0299 |
| | Labels | Average | 0.9581 | 0.9483 | 0.8935 | 0.9128 | 0.8776 | 0.7881 | 0.8940 | 0.8935 |
| | | STD | 0.0154 | 0.0194 | 0.0313 | 0.0370 | 0.0441 | 0.0624 | 0.0314 | 0.0313 |
| | | *P* value | 0.0013 | 0.0205 | 0.0016 | 0.0124 | 0.2120 | 0.0018 | 0.0066 | 0.0016 |
| **PLK3** | No labels | Average | 0.9159 | 0.9154 | 0.8333 | 0.8317 | 0.8458 | 0.6653 | 0.8375 | 0.8333 |
| | | STD | 0.0303 | 0.0299 | 0.0401 | 0.0669 | 0.0369 | 0.0789 | 0.0449 | 0.0401 |
| | Labels | Average | 0.9574 | 0.9442 | 0.9000 | 0.9334 | 0.8706 | 0.8027 | 0.8999 | 0.9000 |
| | | STD | 0.0179 | 0.0361 | 0.0215 | 0.0395 | 0.0342 | 0.0430 | 0.0227 | 0.0215 |
| | | *P* value | 0.0031 | 0.0825 | 0.0006 | 0.0014 | 0.1573 | 0.0004 | 0.0025 | 0.0006 |
| **QIK** | No labels | Average | 0.8286 | 0.8240 | 0.7562 | 0.7804 | 0.7571 | 0.5122 | 0.7670 | 0.7562 |
| | | STD | 0.0222 | 0.0292 | 0.0317 | 0.0478 | 0.0436 | 0.0666 | 0.0296 | 0.0317 |
| | Labels | Average | 0.8982 | 0.8821 | 0.8226 | 0.8549 | 0.8019 | 0.6484 | 0.8260 | 0.8226 |
| | | STD | 0.0126 | 0.0349 | 0.0244 | 0.0466 | 0.0361 | 0.0479 | 0.0212 | 0.0244 |
| | | *P* value | 0.0000 | 0.0013 | 0.0001 | 0.0036 | 0.0289 | 0.0001 | 0.0002 | 0.0001 |
| **RSK3** | No labels | Average | 0.8622 | 0.8573 | 0.7693 | 0.7986 | 0.7628 | 0.5433 | 0.7772 | 0.7693 |
| | | STD | 0.0321 | 0.0451 | 0.0427 | 0.0629 | 0.0677 | 0.0813 | 0.0450 | 0.0427 |

| | | | | | | | | | | |
|---|---|---|---|---|---|---|---|---|---|---|
| | Labels | Average | 0.9140 | 0.9062 | 0.8336 | 0.8516 | 0.8210 | 0.6676 | 0.8347 | 0.8336 |
| | | STD | 0.0190 | 0.0309 | 0.0313 | 0.0432 | 0.0518 | 0.0618 | 0.0354 | 0.0313 |
| | | *P* value | 0.0009 | 0.0164 | 0.0021 | 0.0535 | 0.0565 | 0.0020 | 0.0078 | 0.0021 |
| **SGK3** | No labels | Average | 0.8801 | 0.8625 | 0.7990 | 0.8338 | 0.7675 | 0.6048 | 0.7963 | 0.7990 |
| | | STD | 0.0244 | 0.0406 | 0.0351 | 0.0692 | 0.0635 | 0.0728 | 0.0441 | 0.0351 |
| | Labels | Average | 0.9330 | 0.9288 | 0.8519 | 0.8400 | 0.8669 | 0.7062 | 0.8511 | 0.8519 |
| | | STD | 0.0317 | 0.0364 | 0.0385 | 0.0429 | 0.0664 | 0.0784 | 0.0359 | 0.0385 |
| | | *P* value | 0.0010 | 0.0018 | 0.0070 | 0.8238 | 0.0045 | 0.0109 | 0.0100 | 0.0070 |
| **SIK** | No labels | Average | 0.8601 | 0.8503 | 0.7695 | 0.7584 | 0.7695 | 0.5447 | 0.7596 | 0.7695 |
| | | STD | 0.0309 | 0.0323 | 0.0321 | 0.0838 | 0.0475 | 0.0614 | 0.0376 | 0.0321 |
| | Labels | Average | 0.8943 | 0.8820 | 0.8158 | 0.8315 | 0.8137 | 0.6344 | 0.8198 | 0.8158 |
| | | STD | 0.0365 | 0.0283 | 0.0512 | 0.0775 | 0.0379 | 0.0991 | 0.0418 | 0.0512 |
| | | *P* value | 0.0463 | 0.0397 | 0.0361 | 0.0708 | 0.0434 | 0.0357 | 0.0049 | 0.0361 |
| **TAK1** | No labels | Average | 0.8779 | 0.9014 | 0.7823 | 0.7789 | 0.8111 | 0.5631 | 0.7924 | 0.7823 |
| | | STD | 0.0429 | 0.0285 | 0.0383 | 0.0713 | 0.0394 | 0.0844 | 0.0366 | 0.0383 |
| | Labels | Average | 0.9332 | 0.9133 | 0.8532 | 0.8662 | 0.8499 | 0.7146 | 0.8526 | 0.8532 |
| | | STD | 0.0254 | 0.0549 | 0.0522 | 0.0985 | 0.0664 | 0.0982 | 0.0552 | 0.0522 |
| | | *P* value | 0.0047 | 0.5755 | 0.0045 | 0.0466 | 0.1532 | 0.0026 | 0.0151 | 0.0045 |
| **TAO2** | No labels | Average | 0.8511 | 0.8316 | 0.7693 | 0.7817 | 0.7714 | 0.5408 | 0.7746 | 0.7693 |
| | | STD | 0.0340 | 0.0611 | 0.0466 | 0.0553 | 0.0727 | 0.0919 | 0.0525 | 0.0466 |
| | Labels | Average | 0.9414 | 0.9390 | 0.8669 | 0.8681 | 0.8659 | 0.7352 | 0.8654 | 0.8669 |
| | | STD | 0.0205 | 0.0180 | 0.0365 | 0.0593 | 0.0411 | 0.0714 | 0.0369 | 0.0365 |
| | | *P* value | 0.0000 | 0.0004 | 0.0001 | 0.0050 | 0.0043 | 0.0001 | 0.0006 | 0.0001 |
| **TLK1** | No labels | Average | 0.9137 | 0.8937 | 0.8214 | 0.7929 | 0.8476 | 0.6481 | 0.8154 | 0.8214 |
| | | STD | 0.0300 | 0.0485 | 0.0300 | 0.0643 | 0.0727 | 0.0612 | 0.0404 | 0.0300 |
| | Labels | Average | 0.9353 | 0.9215 | 0.8643 | 0.8631 | 0.8673 | 0.7271 | 0.8637 | 0.8643 |
| | | STD | 0.0344 | 0.0421 | 0.0365 | 0.0608 | 0.0418 | 0.0753 | 0.0378 | 0.0365 |
| | | *P* value | 0.1739 | 0.2106 | 0.0144 | 0.0286 | 0.4905 | 0.0257 | 0.0173 | 0.0144 |
| **TNIK** | No labels | Average | 0.8711 | 0.8445 | 0.8000 | 0.8250 | 0.7723 | 0.6035 | 0.7959 | 0.8000 |
| | | STD | 0.0396 | 0.0494 | 0.0332 | 0.0712 | 0.0373 | 0.0691 | 0.0396 | 0.0332 |

| | | | | | | | | | | |
|---|---|---|---|---|---|---|---|---|---|---|
| | Labels | Average | 0.9591 | 0.9547 | 0.8936 | 0.8791 | 0.9021 | 0.7870 | 0.8897 | 0.8936 |
| | | STD | 0.0154 | 0.0250 | 0.0220 | 0.0337 | 0.0392 | 0.0446 | 0.0271 | 0.0220 |
| | | *P* value | 0.0001 | 0.0000 | 0.0000 | 0.0604 | 0.0000 | 0.0000 | 0.0000 | 0.0000 |
| **TSSK1** | No labels | Average | 0.8705 | 0.8472 | 0.7941 | 0.8023 | 0.7934 | 0.5948 | 0.7937 | 0.7941 |
| | | STD | 0.0326 | 0.0606 | 0.0423 | 0.0873 | 0.0661 | 0.0847 | 0.0511 | 0.0423 |
| | Labels | Average | 0.8838 | 0.8737 | 0.8049 | 0.7820 | 0.8400 | 0.6157 | 0.8068 | 0.8049 |
| | | STD | 0.0343 | 0.0426 | 0.0373 | 0.0724 | 0.0520 | 0.0695 | 0.0414 | 0.0373 |
| | | *P* value | 0.4084 | 0.2979 | 0.5734 | 0.5973 | 0.1144 | 0.5735 | 0.5574 | 0.5734 |
| **TTBK1** | No labels | Average | 0.8807 | 0.8607 | 0.8094 | 0.8399 | 0.7882 | 0.6224 | 0.8118 | 0.8094 |
| | | STD | 0.0142 | 0.0258 | 0.0219 | 0.0490 | 0.0286 | 0.0441 | 0.0219 | 0.0219 |
| | Labels | Average | 0.9620 | 0.9577 | 0.8981 | 0.9110 | 0.8857 | 0.7967 | 0.8977 | 0.8981 |
| | | STD | 0.0111 | 0.0190 | 0.0168 | 0.0298 | 0.0273 | 0.0341 | 0.0173 | 0.0168 |
| | | *P* value | 0.0000 | 0.0000 | 0.0000 | 0.0021 | 0.0000 | 0.0000 | 0.0000 | 0.0000 |
| **YSK1** | No labels | Average | 0.8873 | 0.8684 | 0.7993 | 0.8274 | 0.7819 | 0.6034 | 0.8018 | 0.7993 |
| | | STD | 0.0348 | 0.0727 | 0.0423 | 0.0643 | 0.0686 | 0.0858 | 0.0508 | 0.0423 |
| | Labels | Average | 0.9504 | 0.9521 | 0.8787 | 0.9016 | 0.8671 | 0.7574 | 0.8835 | 0.8787 |
| | | STD | 0.0196 | 0.0194 | 0.0255 | 0.0318 | 0.0389 | 0.0516 | 0.0278 | 0.0255 |
| | | *P* value | 0.0003 | 0.0072 | 0.0002 | 0.0083 | 0.0058 | 0.0004 | 0.0008 | 0.0002 |
| **ZAK** | No labels | Average | 0.8985 | 0.8756 | 0.8234 | 0.8299 | 0.8181 | 0.6516 | 0.8210 | 0.8234 |
| | | STD | 0.0253 | 0.0452 | 0.0340 | 0.0464 | 0.0737 | 0.0669 | 0.0379 | 0.0340 |
| | Labels | Average | 0.9579 | 0.9471 | 0.8928 | 0.9092 | 0.8786 | 0.7893 | 0.8915 | 0.8928 |
| | | STD | 0.0222 | 0.0414 | 0.0341 | 0.0535 | 0.0583 | 0.0655 | 0.0382 | 0.0341 |
| | | *P* value | 0.0001 | 0.0026 | 0.0004 | 0.0036 | 0.0703 | 0.0003 | 0.0010 | 0.0004 |
| **p38α** | No labels | Average | 0.8764 | 0.8602 | 0.7906 | 0.7677 | 0.8069 | 0.5825 | 0.7846 | 0.7906 |
| | | STD | 0.0274 | 0.0514 | 0.0271 | 0.0580 | 0.0606 | 0.0544 | 0.0423 | 0.0271 |
| | Labels | Average | 0.9362 | 0.9319 | 0.8724 | 0.8952 | 0.8658 | 0.7444 | 0.8796 | 0.8724 |
| | | STD | 0.0188 | 0.0285 | 0.0317 | 0.0418 | 0.0355 | 0.0638 | 0.0312 | 0.0317 |
| | | *P* value | 0.0001 | 0.0026 | 0.0000 | 0.0001 | 0.0242 | 0.0000 | 0.0001 | 0.0000 |
| **p38β** | No labels | Average | 0.8728 | 0.8695 | 0.8044 | 0.8160 | 0.8079 | 0.6115 | 0.8092 | 0.8044 |
| | | STD | 0.0331 | 0.0565 | 0.0380 | 0.0706 | 0.0671 | 0.0813 | 0.0470 | 0.0380 |

| | | | | | | | | | | |
|---|---|---|---|---|---|---|---|---|---|---|
| | Labels | Average | 0.9323 | 0.9241 | 0.8626 | 0.8834 | 0.8467 | 0.7299 | 0.8616 | 0.8626 |
| | | STD | 0.0202 | 0.0195 | 0.0204 | 0.0637 | 0.0475 | 0.0448 | 0.0212 | 0.0204 |
| | | *P* value | 0.0003 | 0.0192 | 0.0012 | 0.0476 | 0.1763 | 0.0018 | 0.0097 | 0.0012 |
| **p38δ** | No labels | Average | 0.9001 | 0.8736 | 0.8269 | 0.8224 | 0.8181 | 0.6523 | 0.8192 | 0.8269 |
| | | STD | 0.0161 | 0.0293 | 0.0215 | 0.0365 | 0.0336 | 0.0416 | 0.0200 | 0.0215 |
| | Labels | Average | 0.9413 | 0.9323 | 0.8731 | 0.8740 | 0.8712 | 0.7471 | 0.8713 | 0.8731 |
| | | STD | 0.0238 | 0.0268 | 0.0295 | 0.0468 | 0.0467 | 0.0592 | 0.0328 | 0.0295 |
| | | *P* value | 0.0006 | 0.0003 | 0.0015 | 0.0182 | 0.0135 | 0.0012 | 0.0010 | 0.0015 |
| **p38γ** | No labels | Average | 0.9036 | 0.8966 | 0.8379 | 0.8269 | 0.8432 | 0.6744 | 0.8341 | 0.8379 |
| | | STD | 0.0204 | 0.0303 | 0.0288 | 0.0440 | 0.0289 | 0.0552 | 0.0260 | 0.0288 |
| | Labels | Average | 0.9440 | 0.9421 | 0.8697 | 0.8818 | 0.8638 | 0.7380 | 0.8709 | 0.8697 |
| | | STD | 0.0226 | 0.0335 | 0.0374 | 0.0530 | 0.0572 | 0.0710 | 0.0407 | 0.0374 |
| | | *P* value | 0.0009 | 0.0074 | 0.0592 | 0.0283 | 0.3518 | 0.0491 | 0.0367 | 0.0592 |
| **p70S6K** | No labels | Average | 0.9115 | 0.8939 | 0.8268 | 0.8094 | 0.8248 | 0.6541 | 0.8138 | 0.8268 |
| | | STD | 0.0351 | 0.0421 | 0.0456 | 0.0868 | 0.0547 | 0.0900 | 0.0535 | 0.0456 |
| | Labels | Average | 0.9488 | 0.9412 | 0.8928 | 0.8926 | 0.8944 | 0.7858 | 0.8924 | 0.8928 |
| | | STD | 0.0188 | 0.0264 | 0.0361 | 0.0409 | 0.0536 | 0.0711 | 0.0368 | 0.0361 |
| | | *P* value | 0.0140 | 0.0119 | 0.0034 | 0.0221 | 0.0138 | 0.0031 | 0.0023 | 0.0034 |

#### 10. Additional Cross-Model Figures

##### 10.1. Human O-GlcNAc Prediction with Phosphorylation (S/T) Sites

| PTM | | | AUC | AUPRC | Accuracy | Recall | Precision | MCC | F1 | Specificity |
|---|---|---|---|---|---|---|---|---|---|---|
| **O-GlcNAc Atlas, Human-Only** | No labels | Avg | 0.8099 | 0.8161 | 0.7267 | 0.6940 | 0.7474 | 0.4563 | 0.7186 | 0.7267 |
| | | STD | 0.0158 | 0.0167 | 0.0136 | 0.0363 | 0.0280 | 0.0270 | 0.0159 | 0.0136 |
| | GlcNAc Labels | Avg | 0.9122 | 0.9235 | 0.8281 | 0.7668 | 0.8717 | 0.6612 | 0.8154 | 0.8281 |
| | | STD | 0.0037 | 0.0046 | 0.0064 | 0.0221 | 0.0226 | 0.0135 | 0.0084 | 0.0064 |
| | | *P* value | 0.0000 | 0.0000 | 0.0000 | 0.0001 | 0.0000 | 0.0000 | 0.0000 | 0.0000 |
| | Phos Labels | Avg | 0.8157 | 0.8194 | 0.7322 | 0.6919 | 0.7549 | 0.4667 | 0.7215 | 0.7322 |
| | | STD | 0.0098 | 0.0131 | 0.0097 | 0.0350 | 0.0113 | 0.0183 | 0.0173 | 0.0097 |
| | | *P* value | 0.3422 | 0.6227 | 0.3310 | 0.8972 | 0.4463 | 0.3340 | 0.7042 | 0.3310 |

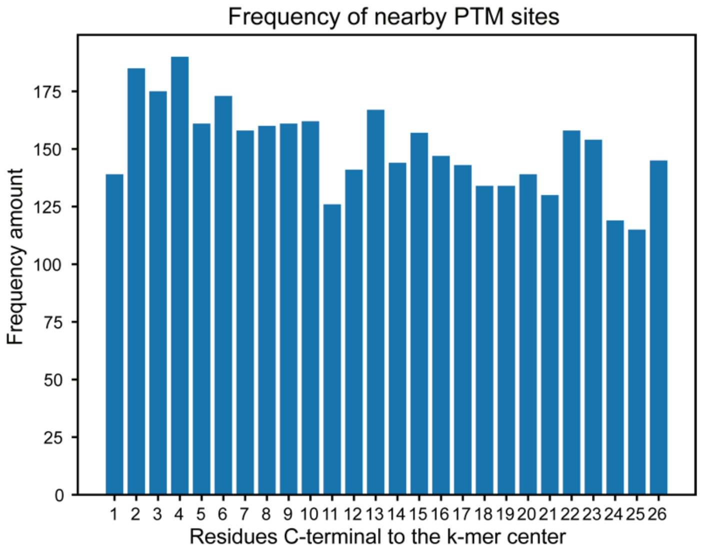

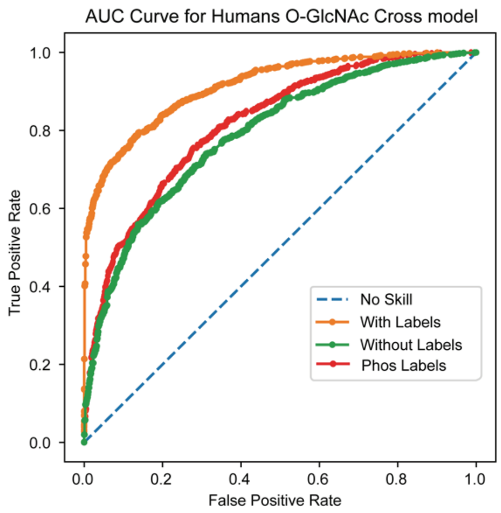
AUC Curve for Humans O-GlcNAc Cross model
True Positive Rate
False Positive Rate
0.0
0.2
0.4
0.6
0.8
1.0
No Skill
With Labels
Without Labels
Phos Labels

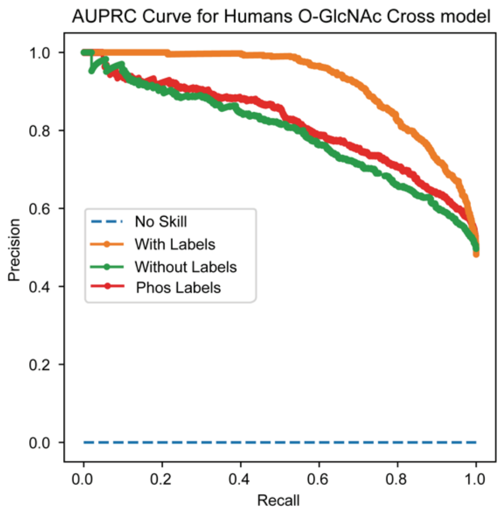
AUPRC Curve for Humans O-GlcNAc Cross model
Precision
Recall
0.0
0.2
0.4
0.6
0.8
1.0
No Skill
With Labels
Without Labels
Phos Labels

#### 11. Effect of Homology on Model Performance

##### 11.1. Evaluation of Sequence Identity

To accurately evaluate the performance of a sequence-based deep learning model, it is important to consider sequence similarity between the training data and independent test sets. Given that we are evaluating enzymatic PTMs, some level of sequence similarity is expected in some cases because enzymes typically recognize particular sequences. For every model we ensured that no k-mers present in the training and validation data were present in the test data (*i.e.,* a sequence similarity of 100%). To further evaluate how similarities impact our datasets, we then used CD-Hit[8] to remove sequences from the test sets that shared different levels of sequence similarity down to a cutoff of 40% sequence identity. The results of this evaluation are shown in Supplementary Fig. 11.2–11.5. This evaluation was not done for kinase-specific datasets because they are small datasets already and because particular kinases likely have specific recognition sites that might require high sequence identity and we want our models to capture that.

For some modifications, specifically the hydroxylation of lysine and proline, there were no instances of sequences below a certain identity cutoff, which indicates that proteins containing these modifications are likely to be homologous (which makes sense because the proteins having these PTMs are often collagens). For most modifications, we saw a slight decrease in performance as we decreased sequence identity. More interestingly, the labeled model performance decreased more than did the unlabeled models. This dichotomy is likely due, in part, to a technical limitation, as k-mers with modified amino acids had to be converted to canonical amino acids for comparison using CD-Hit and then converted back. Inclusion of modified amino acids generally lowers the sequence identity, so those sets in our model likely have lower levels of sequence identity than do the unlabeled sets. Additionally, this decrease could also indicate that incorporating PTM labels allows a model to distinguish between high-identity sequences and would likely increase its performance in context-specific prediction (where some potential modification sites are modified whereas others are not). Regardless, labeled models that perform significantly better than their unlabeled counterpart generally maintain this advantage at lower levels of sequence identity though to a lesser extent.

Finally, it is important to note that sequence identity greatly depends on whether an analysis is performed at a protein or peptide level (*e.g.*, our 53 k-mer) and, in the latter case, on the length of the peptide fragment.

#### 11.2. Using CD-Hit to Lower Test-Set Homology in Musite Deep Datasets

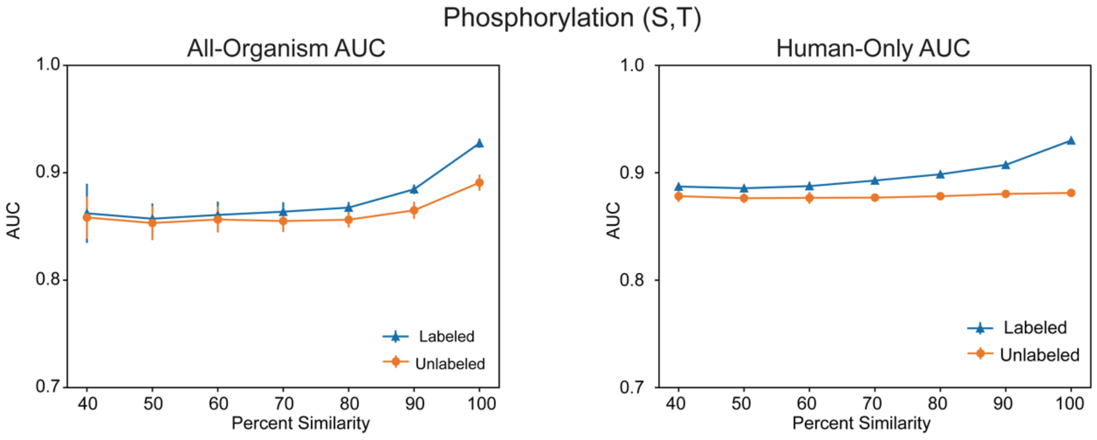


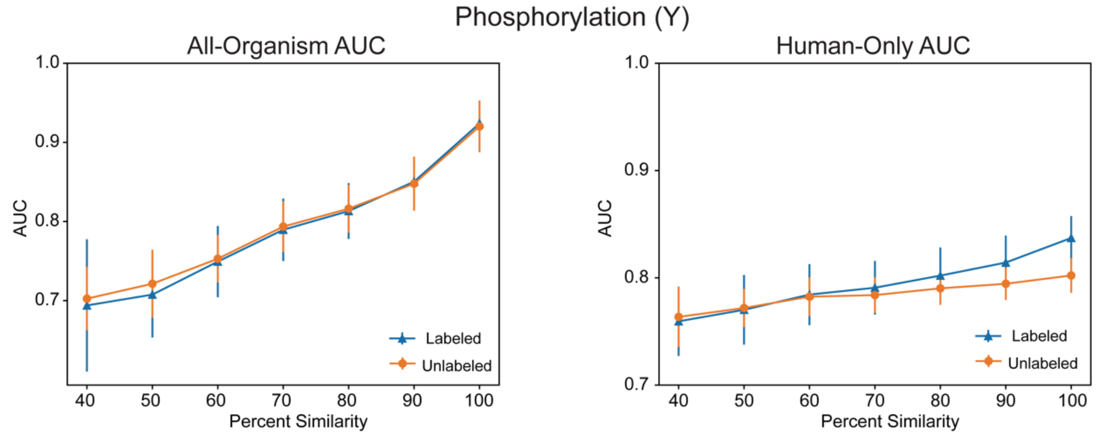


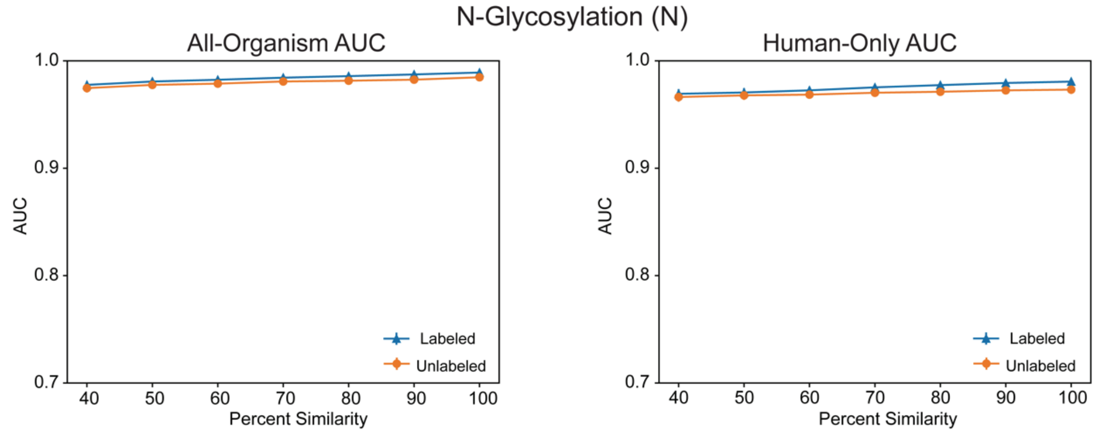

O-Glycosylation (S,T)

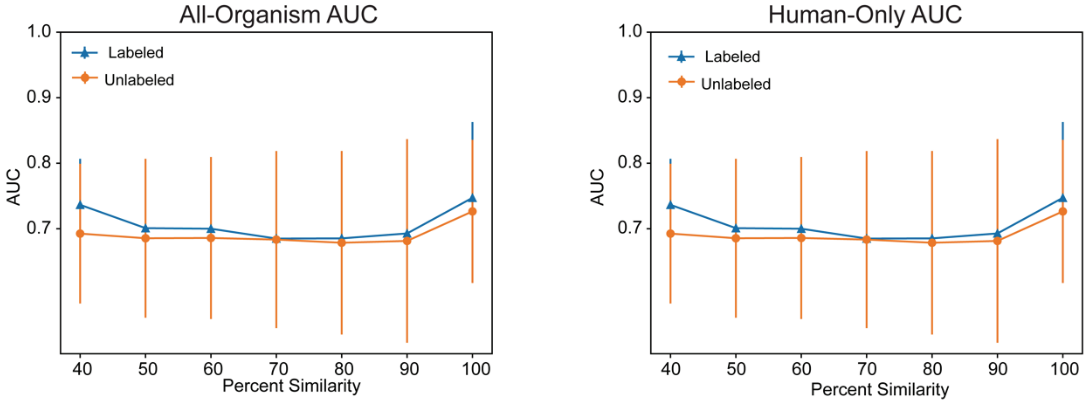


Ubiquitination (K)

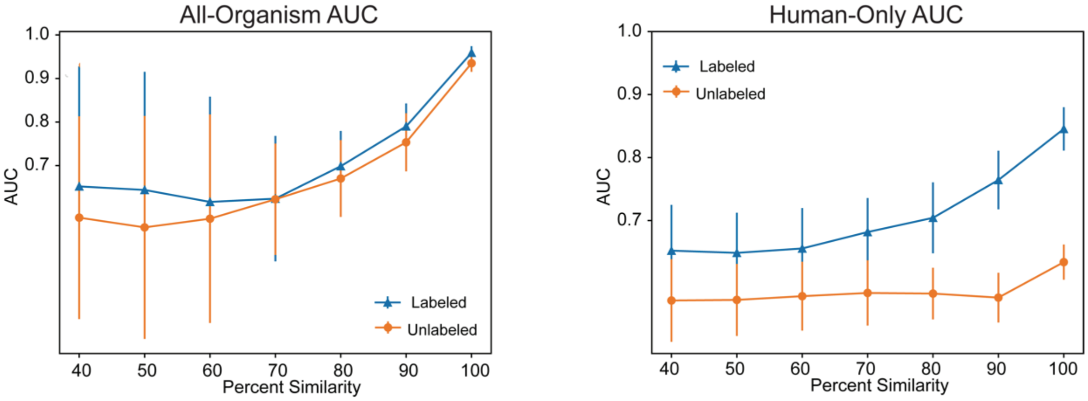


SUMOylation (K)

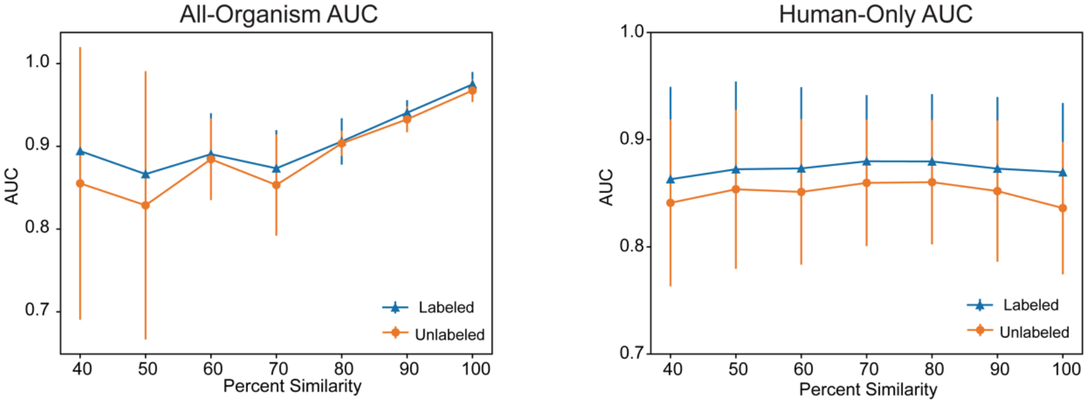

#### Acetylation (K)

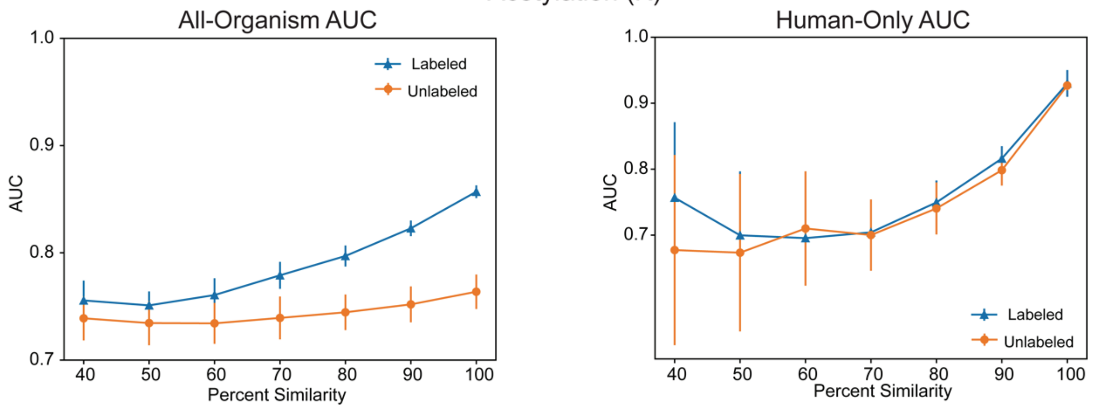


#### Methylation (K)

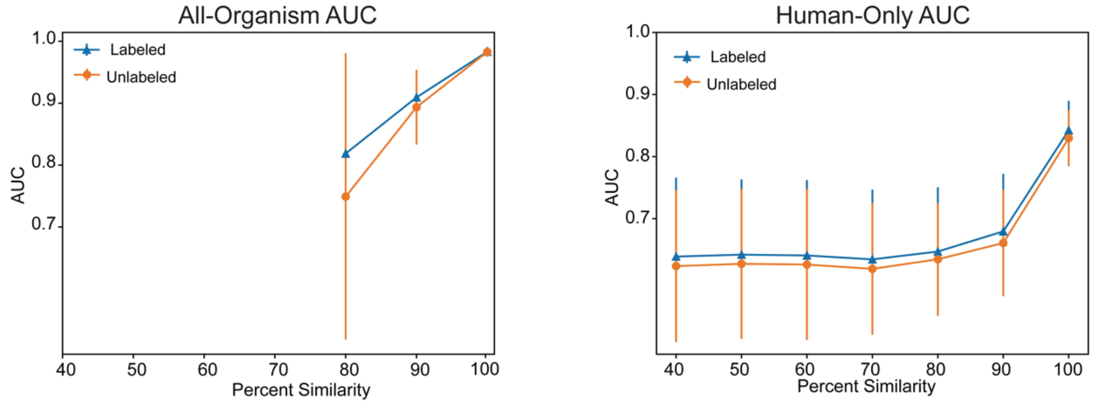


#### Methylation (R)

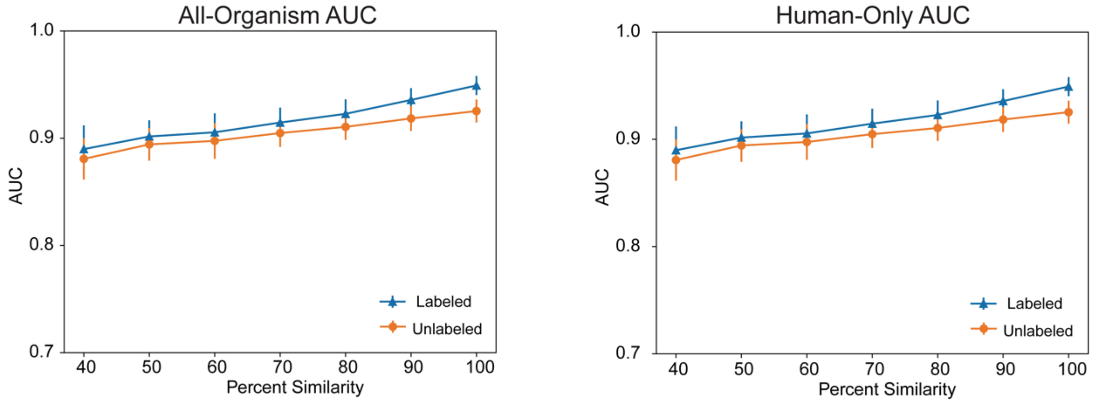

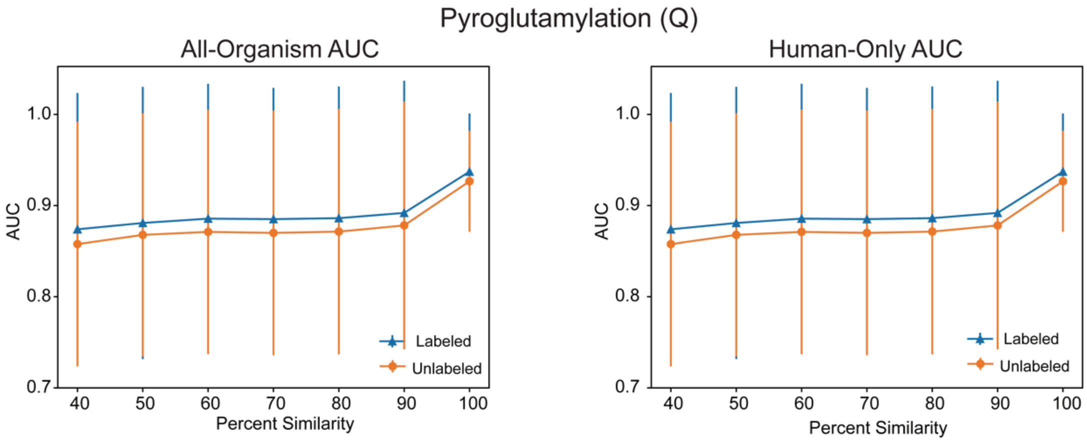
Pyroglutamylation (Q)
All-Organism AUC
Human-Only AUC
AUC
Percent Similarity
Labeled
Unlabeled
1.0
0.9
0.8
0.7
40
50
60
70
80
90
100

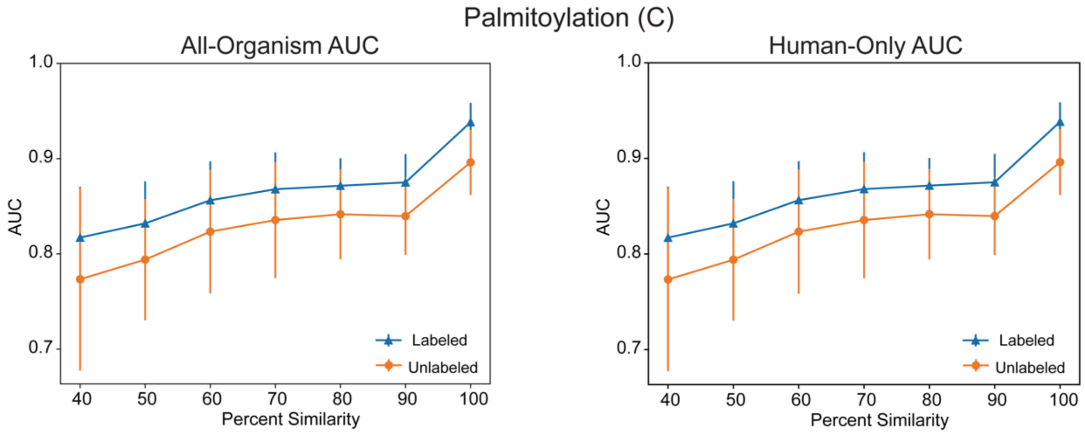
Palmitoylation (C)
All-Organism AUC
Human-Only AUC
AUC
Percent Similarity
Labeled
Unlabeled
1.0
0.9
0.8
0.7
40
50
60
70
80
90
100

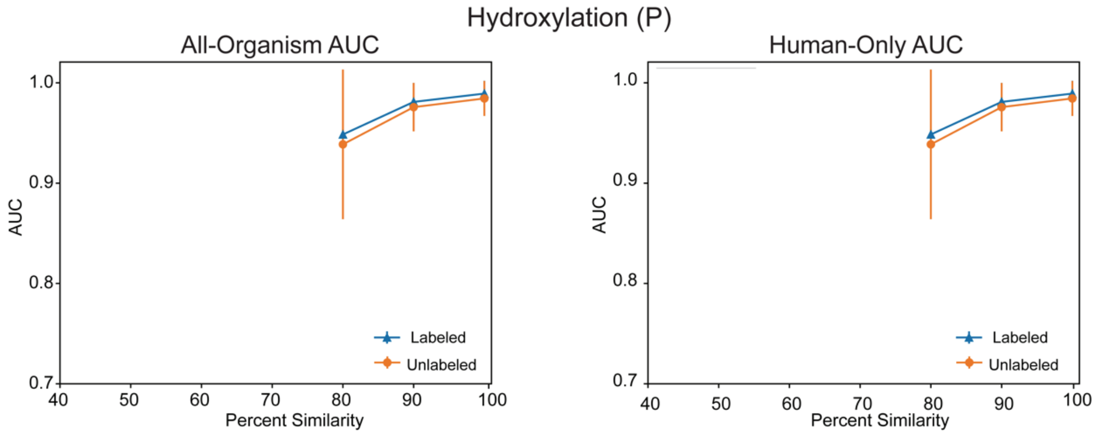
Hydroxylation (P)
All-Organism AUC
Human-Only AUC
AUC
Percent Similarity
Labeled
Unlabeled
1.0
0.9
0.8
0.7
40
50
60
70
80
90
100

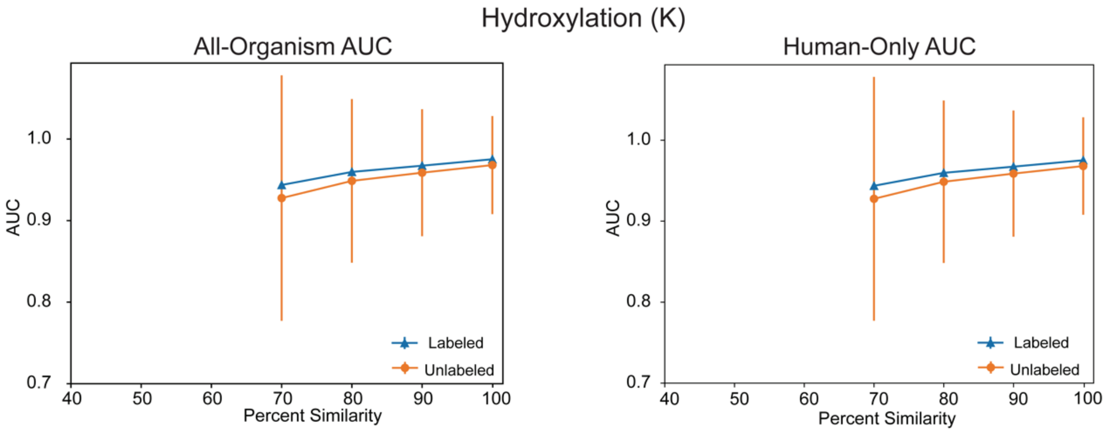

Hydroxylation (K)
All-Organism AUC
Human-Only AUC
AUC
1.0
0.9
0.8
0.7
40
50
60
70
80
90
100
Percent Similarity
Labeled
Unlabeled

#### 11.3. Using CD-Hit to Lower Test-Set Homology in OGP Datasets

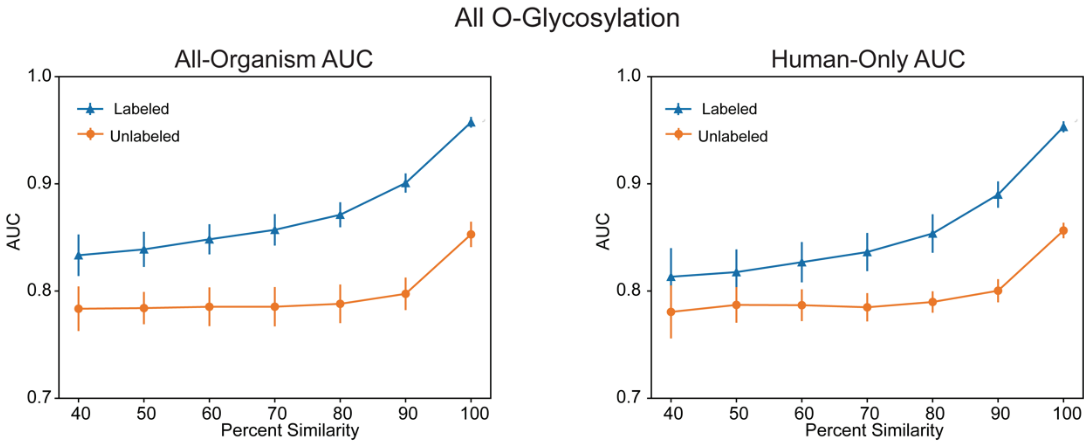


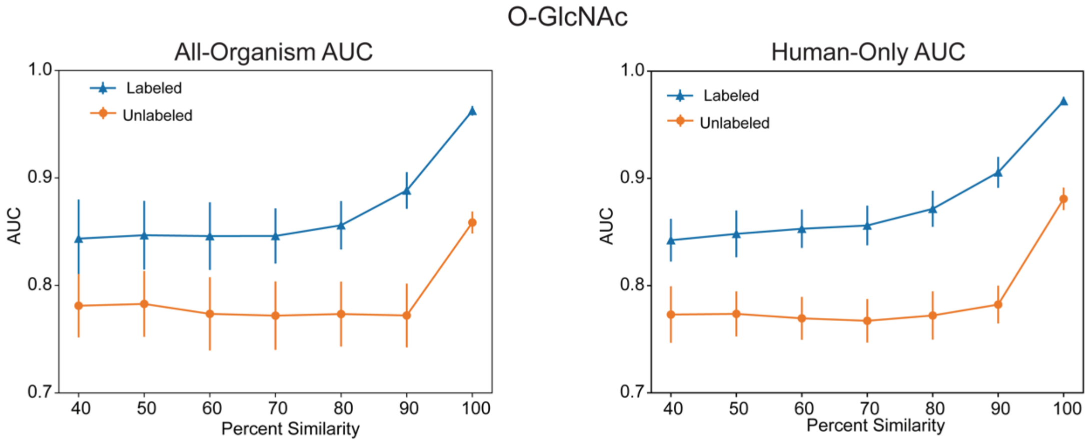


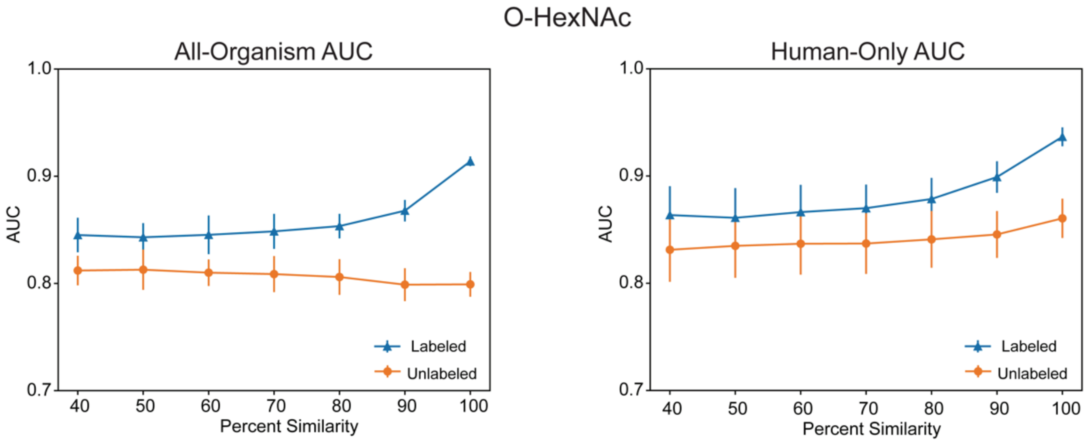

##### 11.4. Using CD-Hit to Lower Test-Set Homology in N-Glycosylation Sequon-Specific Datasets

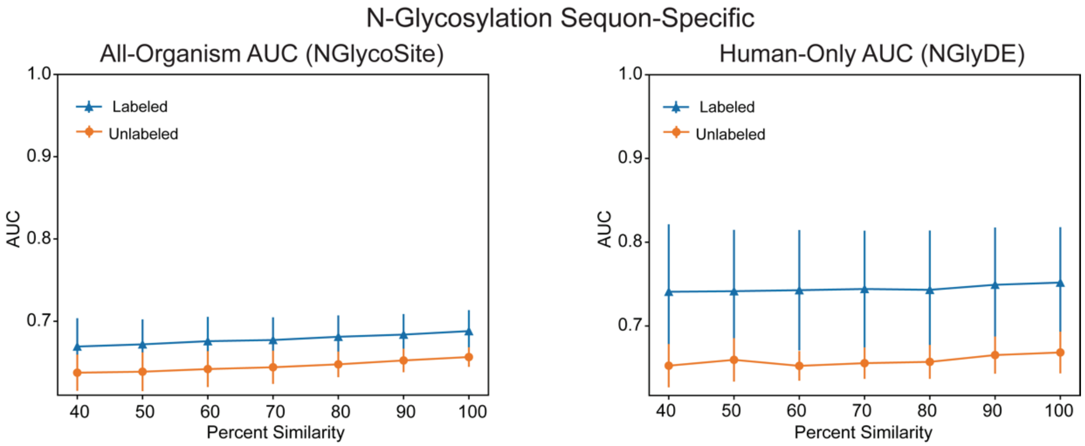


##### 11.5. Using CD-Hit to Lower Test-Set Homology in O-GlcNAc Atlas Datasets

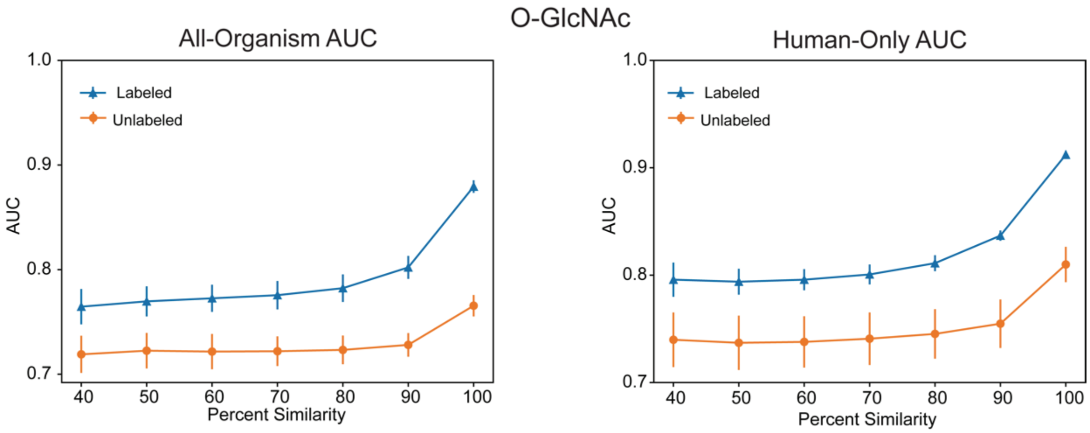
